# Identification and binding-site mapping of RNA-binding proteins interacting with microRNA precursors in Arabidopsis

**DOI:** 10.64898/2026.07.28.741191

**Authors:** Andreas R. Winkel, Valentin W. Bitterer, Martin Lewinski, Marlene Reichel, Janina Lüders, Marion Scheibe, Maria Kalyna, Tom Laloum, Paula Duque, Dorothee Staiger, Falk Butter, Tino Köster

**Affiliations:** RNA Biology and Molecular Physiology, Faculty of Biology, Bielefeld University, 33615 Bielefeld, Germany; Institute of Molecular Biology (IMB), Ackermannweg 4, 55128 Mainz, Germany; Friedrich-Löffler-Institut (FLI), Südufer 10, 17493 Greifswald, Germany; Institute of Molecular Plant Biology, Department of Biotechnology and Food Science, University of Natural Resources and Life Sciences (BOKU), 1190 Vienna, Austria; GIMM - Gulbenkian Institute for Molecular Medicine, Lisbon, Portugal

**Author notes:** Correspondence: Dr. Tino Köster. Authors contributed equally to this work (Andreas R. Winkel, Valentin W. Bitterer). Section for Computational and RNA Biology, Department of Biology, University of Copenhagen, 2200 Copenhagen N, Denmark. Université Côte d’Azur, Nice, France (Tom Laloum).

**Keywords:** miRNAs, SR proteins, *Arabidopsis thaliana*, pri-miRNA processing, RNA pulldown, RNA-binding protein, iCLIP

## Abstract

**Background:** MicroRNAs (miRNAs) play key roles in modulating gene expression. Upon transcription, primary miRNA transcripts (pri-miRNAs or *MIRNAs*) fold into stem-loop structures. Endonucleolytic cleavage releases a miRNA/miRNA* duplex from the stem, which is subsequently matured. In higher plants, the pri-miRNA hairpins vary widely in length and structure. Pri-miRNA processing is extensively regulated by RNA-binding proteins (RBPs) and the range of underlying mechanisms continues to expand.

**Results:** Here, we performed an unbiased screen to identify *Arabidopisis thaliana* RBPs that interact with pri-miRNAs, using *pri-miRNA159a* and *pri-miRNA398b* as paradigms. Nucleoplasmic proteins were significantly enriched in pulldowns with bead-immobilized *in vitro* transcripts of the stem-loop regions of these pri-miRNAs compared with pulldowns using empty beads. The enriched proteins included several members of the family of the serine/arginine-rich (SR) splicing factor family, including RS31 and SR34a. Mutant analysis indicated that selected candidate interactors affect pri-miRNA and/or mature miRNA accumulation. To validate interactions with pri-miRNAs *in vivo*, we performed individual-nucleotide resolution crosslinking and immunoprecipitation (iCLIP). Whereas our conventional plant iCLIP procedure detected only a few crosslink sites on pri-miRNAs, likely because of the low abundance of these transcripts, our improved plant iCLIP2 protocol enabled the identification of binding sites on numerous pri-miRNAs. RS31 and SR34a bound overlapping sets of pri-miRNAs and contacted both stem-loop regions and flanking regions upstream and downstream of the stem-loop. RS31 and SR34 bound both overlapping and distinct regions of pri-miRNAs.

**Conclusion:** *In vitro* pulldowns of nucleoplasmic proteins identified RBPs associated with pri-miRNA stem-loops, and the improved plant iCLIP2 protocol revealed *in vivo* binding of RS31 and SR34a to a subset of pri-miRNAs. Together, these findings expand the repertoire of RBPs implicated in plant miRNA biogenesis and support a role for SR proteins as candidate regulators of pri-miRNA processing.

## Background

MicroRNAs (miRNAs) are noncoding RNAs with a length of 20-24 nucleotides (nt) that regulate the expression of cognate target genes by eliciting mRNA cleavage or translational inhibition [1, 2]. MiRNAs are generated from RNA polymerase II transcripts with internal stem-loop structures, the primary miRNAs (pri-miRNAs). A first cleavage by an RNase III protein releases the stem-loop (precursor miRNA, pre-miRNA) which is further processed to the miRNA/miRNA* duplex. After methylation of the 3’ ends by HUA ENHANCER1 (HEN1), the mature miRNA is loaded into ARGONAUTE 1 (AGO1) while the miRNA* strand is mostly degraded [3–5]. In higher plants, the endonucleolytic cleavages are performed mainly by the RNase III type enzyme DICER-LIKE1 (DCL1) [6]. Along with DCL1, the double-stranded RNA-binding protein (RBP) HYPONASTIC LEAVES 1 (HYL1) and the zinc finger protein SERRATE (SE), collectively designated as the microprocessor complex, are required to ensure efficient and accurate processing [7–9]. More recently, it was found that phase separation of SE promotes pri-miRNA processing by the microprocessor complex [10]. Apart from the microprocessor complex, a suite of proteins affects pri-miRNA processing. DAWDLE (DDL) and PLEIOTROPIC REGULATORY LOCUS 1 (PRL1) stabilize pri-miRNAs and promote miRNA biogenesis [11, 12]. Additional members of the MOS4-ASSOCIATED COMPLEX interact with and promote the activity of DCL1 [12–15]. SMALL1 (SMA1), a homolog of the mammalian DEAD-box pre-mRNA splicing factor Prp28, interacts with the DCL1 complex and enhances the efficiency of miRNA processing [16]. TOUGH (TGH) interacts with pri-miRNAs and promotes their interaction with HYL1 [17].

It is increasingly becoming clear that the secondary structure of the pri-miRNA stem-loop plays a critical role in processing. For example, CHROMATIN REMODELLING 2 (CHR2), the ATPase subunit of the SWI/SNF chromatin remodelling complex, displays RNA helicase activity to unwind the pri-miRNA stem-loops, leading to reduced levels of mature miRNAs [18].

Pri-miRNAs contain a lower stem and flanking single-stranded basal segments below the double-stranded region corresponding to the mature miRNA and its complementary miRNA*. Above the duplex region, there are an upper stem and the terminal loop. The initial cleavage by DCL1 occurs 15-17 nt away from the ssRNA-dsRNA junction. Due to extensive heterogeneity in length and structure of pri-miRNA stem-loops, different modes of plant pri-miRNA processing have been distinguished based on the direction and number of cuts in higher plants [19]. Accordingly, pri-miRNAs can be processed from base-to-loop or from loop-to-base [19, 20]. In the “short base-to-loop” processing mode (*e.g*., pri-miRNA 172 and pri-miR398 families), an imperfectly paired lower stem of ∼15 nt below the miRNA/miRNA* duplex followed by an internal loop determines the site of the initial pri-miRNA cleavage [21]. Subsequent cleavage of the pre-miRNA at a fixed distance of ∼21 nt from the first cut then releases the miRNA/miRNA* duplex. Two pri-miRNA families, pri-miR156 and pri-miR160, are processed by a “short loop-to-base” mechanism where the first cut is guided by an upper stem segment and a small loop.

Besides, further modes of processing exist in which two additional cuts are required to release the mature miRNA (sequential base-to-loop or sequential loop-to-base) [19, 20]. These successive cuts along the stem allow for generation of additional low abundant small RNAs. In the pri-miR159 and pri-miR319 families with their long stem segments the first cut occurs near the loop and three more sequential cuts follow to release the mature miRNA in this “sequential loop-to-base” mechanism.

Transcription of pri-miRNA genes widely responds to endogenous cues and environmental stimuli including abiotic and biotic stress [22]. Processing of pri-miRNAs occurs co-transcriptionally once the secondary structure stem-loop is folded [23]. While for pri-miRNAs processed from loop-to-base all steps occur co-transcriptionally, for pri-miRNAs processed from base-to-loop only the first cleavage is executed co-transcriptionally, while further processing occurs on pri-miRNAs released into the nucleoplasm [23]. Notably, some pri-miRNAs can be processed both co-transcriptionally and post-transcriptionally, depending on environmental conditions. Thus, plants exquisitely fine-tune processing of miRNA precursors into mature miRNAs to adjust the miRNome. Accordingly, numerous RBPs have been implicated in the regulation of pri-miRNA processing.

Interaction of several RBPs with pri-miRNAs has been demonstrated using RNA immunoprecipitation [24, 25]. Among those, *Arabidopsis thaliana* GLYCINE-RICH RNA-BINDING PROTEIN 7 binds to selected pri-miRNAs *in vivo* to impact their processing [24]. Arabidopsis SR proteins RS40 and RS41 interact with HYL1/SE, bind pri-miRNAs, and promote the proper processing (and in intron-containing cases, splicing) of a subset of pri-miRNAs *in vivo* [25].

So far, the regions within the precursor that interact with the RBPs remain unknown. Ultraviolet (UV) crosslinking based techniques such as individual-nucleotide resolution crosslinking and immunoprecipitation (iCLIP) determine *in vivo* binding sites of RBPs with single nucleotide resolution [26]. Irradiation with 254-nm UV light is employed to covalently link RNAs and bound proteins to preserve the native interactions. RNA-protein complexes are recovered from lysates by immunoprecipitation and the associated RNAs are radioactively labelled. After purification of the RNA-protein complexes by denaturing gel electrophoresis the region above the expected molecular weight, containing protein with associated RNAs, is processed for RNA isolation to generate sequencing libraries. During reverse transcription, the remnants of the crosslinked protein stall the reverse transcriptase, causing cDNAs to terminate immediately before the crosslink site. Sequencing these cDNAs enables determination of binding sites at single-nucleotide resolution. Although UV crosslinking-based methods were initially developed for mammalian cell monolayers or translucent tissues, iCLIP has recently been adapted for plants, whose multicellular, opaque tissues and rigid cell walls complicate crosslinking and recovery [27, 28].

In the conventional iCLIP procedure, one bipartite linker is used to prime cDNA synthesis, and the attachment of the linker to the cDNA 3’ end is accomplished *via* intramolecular circularization and relinearization. In iCLIP2, a more efficient ligation of the second linker directly to the cDNA 3’ end is used, as done in the infrared CLIP protocol [29]. Furthermore, the protocol includes a pre-amplification step prior to size selection of the cDNAs and uses bead-based instead of gel-based size selection and employs beads instead of purification by repeated ethanol precipitation that was introduced in eCLIP to avoid sample loss [30]. Recently, the updated iCLIP2 protocol which provides increased sensitivity has been implemented in plants [31–34].

To systematically identify RBPs that may impact pri-miRNA processing, we have established RNA affinity pulldown with *in vitro* transcribed pri-miRNA stem-loops incubated with protein extracts from plant nuclei. Bound proteins were identified by mass spectrometry and categorized according to enrichment relative to controls. Binding of the RBPs to the miRNA stem-loops *in vivo* was validated using plant iCLIP2. Mutants deficient in selected interacting RBPs were monitored for alterations in miRNA biogenesis, employing small RNA northern blot hybridization.

## Methods

### Cloning and *in vitro* transcription of pri-miRNA stem-loops

DNA fragments corresponding to the *MIR398b* stem-loop structure including the T7 promoter were synthesized as a gBlock^TM^ (IDT, Coralville, IA, USA) and subcloned into the pUC19 vector via *Eco*RI and *Xba*I restriction sites (Additional file 1, Table S1). For *in vitro* transcription of *MIR398b*-SL, the plasmid pUC19-T7-MIR398b-SL containing the *MIR398b* stem-loop region downstream of the T7 promoter was linearized with *Xba*I and gel purified using the NucleoSpin^®^ Gel and PCR Clean up kit (Macherey-Nagel, Düren, Germany). For *MIR159a*-SL, the annotated *MIR159a* stem-loop structure including the T7 promoter, the 20-nt Csy4-hairpin and spacer sequences were synthesized as a gBlock^TM^ (IDT, Coralville, IA, USA) and subcloned into the pUC19 vector via *Eco*RI and *Hind*III restriction sites (Additional file 1, Table S1) [35]. To omit *in vitro* transcription of the Csy4 hairpin, the stem-loop region of *MIR159a*-SL was PCR-amplified from the plasmid pUC19-T7-Csy4-MIR159a-SL using primers T7-MIR159a-SL_for and MIR159a-SL_rev. The templates for *in vitro* transcription were purified via the NucleoSpin^®^ Gel and PCR Clean up kit (Macherey-Nagel, Düren, Germany). *In vitro* transcription was performed using the MEGA shortscript^TM^ T7 transcription kit (Invitrogen, Carlsbad, CA, USA) in the presence of biotinylated dUTP according to the manufactureŕs instructions. The *in vitro* transcripts were purified using the RNA Clean & Concentrator-25 Kit (Zymo Research, Freiburg, Germany). The secondary structure of *MIR159a-SL* and *MIR398b-SL* were predicted *in silico* using the RNAfold web server tool [36].

### Plant material

All lines used in this study are in the *Arabidopsis thaliana* Col-0 ecotype. The following mutant and transgenic lines have been described previously: *rs31-1* (SALK_021332; At3g61860) [37, 38], 35S::RS31 overexpressing the RS31 coding region (At3g61860) under control of the Cauliflower Mosaic Virus (CaMV) 35S promoter (RS31-ox) [38], RS31::RS31-GFP, expressing RS31 fused to GFP under the control of the endogenous promoter in the *rs31-1* mutant background [38], *rs31a* line carrying an artificial miRNA against *RS31a* (amiR-31a-E2; At2g46610) [39], *sr34a-1* (GK-803A10; AT3G49430) [34], SR34a::SR34a-GFP, expressing SR34a fused to GFP under the control of the endogenous promoter in the *sr34a-1* mutant background, *sr34-1* mutant (SALK_106067; A1g02840) [40], *sr45-1* (SALK_004132; AT1G16610) [41], *brr2a-2* (*cäö*) carrying a G to A transition in the At1g20960 gene that results in a T895I missense mutation [42], *nuc1-2* (SALK_002764; At1g48920) [43], and *rs40 rs41* double mutant (WiscD-sLox382G12 x SAIL_64_C03; AT4G25500, AT5G52040) [44]. Transgenic plants expressing GFP alone under the control of the CaMV 35S promoter have also been described previously [38].

Arabidopsis seeds were surface-sterilized and sown on half-strength MS (Murashige and Skoog; Duchefa, Haarlem, Netherlands) plates [45]. Plants were grown in 12 h light–12 h dark cycles at 20°C in Percival incubators (CLF, Wertingen, Germany). For preparation of nucleoplasmic extracts, RNA analysis and iCLIP, 14-day-old aerial tissue from Col-0 wild type was harvested at dusk.

### Preparation of nucleoplasmic protein extracts

Five grams of Arabidopsis Col-0 wild-type plant material were ground in liquid N_2_ and resuspended in 5 mL nuclear lysis buffer (nLB buffer) (20 mM Tris-HCl pH 7.4, 25% glycerol, 20 mM KCl, 2 mM Na_2_-EDTA pH 8.0, 2.5 mM MgCl_2_, 250 mM sucrose, 5 mM dithiothreitol (DTT)). All following steps were performed on ice or at 4°C. The samples were filtered through a 100 μm nylon filter and two layers of Miracloth (22–25 μm; Calbiochem). The resulting total lysate was centrifuged at 1500× g for 30 min. The supernatant corresponding to the cytosolic fraction was transferred to a fresh tube. The pellet containing the nuclei was resuspended in 12 mL nuclei resuspension buffer (NRBT) (20 mM Tris-HCl pH 7.4, 25% glycerol, 2.5 mM MgCl_2_, 0.2% Triton X-100) and centrifuged again at 1500× g for 15 min. This washing step was repeated four-to-five times until the supernatant became clear. Finally, the pellet was suspended in 3 volumes (∼600 μL) of protein extraction buffer (PEB+) (20 mM HEPES-KOH pH 7.5, 5% glycerol, 1.5 mM MgCl_2_, 0.2 mM Na_2_-EDTA pH 8.0, 420 mM NaCl, 5 mM DTT, 1x cOmplete protease inhibitor (Roche, San Francisco, CA, USA), as previously described [46]. After adjusting the salt concentration to 150 mM NaCl, the pulldowns were performed. Protein concentrations were determined using the Qubit^TM^ Protein Assay Kit (Thermo Fisher Scientific).

### Immunoblot

The immunoblot analysis of total lysate, cytoplasmic and nucleoplasmic fractions was essentially performed as described [27]. Briefly, primary antibodies were polyclonal antibodies against HISTONE 3 (Agrisera, Vännäs, Sweden, AS10710; rabbit; dilution 1:5000), UDP-GLUCOSE PYROPHOSPHORYLASE (Agrisera AS11 1739; rabbit; dilution 1:5,000 and SERRATE (Agrisera AS09532A; rabbit; dilution 1:1000). The secondary antibody was HRP-coupled anti-rabbit IgG (Sigma-Aldrich, Burlington, MA, USA, A0545; dilution 1:2500). Chemiluminescence was detected using either Stella (Raytest) or the Fusion-FX6 system (Vilber, Germany).

### Biotin-based RNA pulldown

Beads loaded with the *in vitro* transcripts or empty beads as control were incubated with 2 mL of nucleoplasmic proteins (800-1100 µg) for 90 min at 4°C. After three washes in 500 µL protein washing buffer beads were either processed for quality control or for mass spectrometry. For RNA analysis, 5% of the slurry (beads after pulldown, BAP) was heated for 5 min at 95°C in formamide containing buffer. For protein analyses, 95% of the slurry was resuspended in 40 µL SDS sample buffer and subjected to SDS-PAGE. For either silver staining or mass spectrometry, the slurry of samples (E+) and the empty beads (E-) were resuspended in 40 µL LDS sample buffer (Novagen^TM^ NuPAGE^TM^ LDS sample buffer, Thermo Scientific) and incubated for 10 min at 70°C.

### Silver staining

Samples were subjected to SDS-PAGE using 4–12% NuPAGE^TM^ Novex Bis-Tris precast gel (Thermo) run at 100 V in 1x MOPS buffer followed by silver staining as described previously [47].

### Mass Spectrometry and Data Analysis

Proteins were separated on a 4–12% NuPAGE^TM^ Novex Bis-Tris precast gel (Thermo) for 8 min at 180 V in MES buffer (Thermo). After fixation and staining with Coomassie Brilliant Blue (Carl Roth, Karlsruhe, Germany), gels were minced and destained in 50% ethanol/25 mM ammonium bicarbonate followed by dehydration in 100% acetonitrile. Subsequently, samples were treated with 10 mM DTT/50 mM ammonium bicarbonate (Sigma-Aldrich) for 1 h at 56°C and with 50 mM iodoacetamide/50 mM ammonium bicarbonate (Sigma-Aldrich) for 45 min at room temperature in the dark. Gel pieces were dehydrated with pure acetonitrile (VWR) and incubated with 1 μg MS-grade trypsin (Serva) at 37°C overnight. Tryptic peptides were extracted twice with 30% acetonitrile and twice with 100% acetonitrile and the mixture was concentrated in a concentrator (Eppendorf) to a final volume of ca. 100 μL. Peptides were desalted on StageTips [48] and separated on a nanocapillary (New Objective) packed in-house with Reprosil C18 (Dr. Maisch GmbH, Ammerbuch, Germany). The column was attached to an Easy nLC 1000 system (Thermo) operated with a 90 min gradient from 5% to 60% acetonitrile in 0.1% formic acid at a flow of 225 nL/min. The spray capillary was mounted on the nanospray ion source of a Q Exactive Plus mass spectrometer (Thermo). Measurements were performed using HCD fragmentation with a data-dependent Top10 MS/MS spectra acquisition scheme per MS full scan in the orbitrap analyzer.

The raw files were processed with MaxQuant (version 1.5.2.8) [49] and searched against the TAIR10 protein database (35,386 entries). MaxQuant standard settings were used, except label-free quantitation [50], without FastLFQ and match between runs option was activated. Prior to statistical analysis, entries from contaminants, reverse hits and only identified by site were removed. Missing values were imputed by a downshifted compressed beta distribution at the limit of quantitation. Enrichment was calculated as the mean of the 4 replicates and the variation was statistically assessed by Welch *t*-test using R. Logarithmic transformations of the enrichment were plotted against the determined *p* value and proteins with a log_2_ fold change ≥ 1.0 and a *p*-value ≤ 0.05 were classified as significantly enriched.

### Analysis of GO Terms and Protein Domains

Genes enriched in gene ontology (GO) terms and protein domains were analysed in the online ThaleMine database using the default background population and Holm–Bonferroni test correction [51].

### Small RNA gel blots

For RNA analysis, at least ten plants were bulked for each sample per replicate and RNA was isolated using TRIzol® reagent (Invitrogen). For small RNA gel blots, 5 μg of total RNA was fractionated on a 17% polyacrylamide 8 M urea gel in Tris-borate-EDTA buffer, transferred onto GeneScreen membrane by capillary blotting and crosslinked with UV-light (254 nm) at 120 mJ/cm^2^. Hybridization was performed with 5′-biotinylated antisense DNA oligonucleotides (Metabion, Planegg, Germany) in hybridisation buffer (7% SDS, 200 mM Na_2_HPO_4_ pH 7.0, 5 μg/mL salmon sperm DNA) (Additional file 1, Table S2). Hybridization was performed for 12–15 h at 38°C with end-over-end rotation and subsequently rinsed with washing buffer (1× SSC, 0.1% SDS) twice for 15 min at 38°C. The biotin-labelled probes were detected using the Nucleic Acid Detection Module Kit (Thermo Fisher Scientific), according to the manufacturer’s instructions. Chemiluminescence detection was done with Stella (Raytest) or the Fusion-FX6 system (Vilber, Germany) and signals were quantified using ImageJ (v. 1.53, Rasband, W.S., NIH, Bethesda, MD, USA) [95]. A Student’s *t*-test was performed to determine statistical significance (*** *p ≤* 0.001, ** *p ≤* 0.01, * *p ≤* 0.05).

### RT-qPCR

For RT-qPCR, RNA was reverse transcribed using AMV reverse transcriptase (EURX Roboklon, Berlin, Germany). qPCR was performed in a volume of 10 μL with the iTaq SYBR GREEN supermix (BioRad, Hercules, CA, USA) using 45 cycles of 15 s at 95°C and 30 s at 60°C in a CFX96 cycler (BioRad). Cq values were determined and relative expression levels were calculated based on non-equal efficiencies for each primer pair [52]. Data were normalized to *PP2A* (At1g13320) and expressed as mean expression levels of the independent biological replicates each ± standard deviation or as indicated in the figure legend. A Student’s t-test was performed to determine statistical significance (*** *p ≤* 0.001, ** *p ≤* 0.01, * *p ≤* 0.05). Primers are listed in Additional file 1, Table S3.

### Plant iCLIP2

RS31::RS31-GFP, SR34a::SR34a-GFP and 35S::GFP plants were subjected to irradiation with 254 nm UV light (2000 mJ cm^-2^) and above-ground tissues were flash frozen. For each replicate, three grams of plant material were processed, as previously described for plant iCLIP2, and reads were processed according to our pipeline [31, 32].

### MIR annotation file

The annotation of genomic pri-miRNA coordinates from mirEX^2^ [53] were combined with data from Lepe-Soltero et al. (Additional file 2, Table S4) [54]. For each *MIRNA*, the annotations spanning the longest genomic range was retained and transformed into BED and GTF format. The sources, where the annotations originate from, were specified in the naming field of the file, i.e. miRXXX_M2 for mirEX2 or miRXXX_LS for Lepe-Soltero. *MIRNAs* with no counterpart in the other source were marked with oM2 (only mirEX^2^) or oLS (only Lepe-Soltero).

### Motif discovery

The sequence consisting of nine nucleotides at each binding locus was extracted utilizing bedtools getfasta in a strand-specific configuration [55] in conjunction with the TAIR10 genome. The process of motif discovery was divided into two distinct methodologies. The initial methodology employed Expectation-Maximization via STREME 5.4.1 [56] to ascertain significant motifs, while the subsequent methodology focused on k-mer distributions surrounding the designated peak position within the binding site region. Both methodologies utilized an identical set of sequences (for the purpose of motif discovery).

### Differential gene expression

Mapped reads in the BAM format were obtained from previous paired-end RNA-sequencing experiments encompassing three biological replicates each of *A. thaliana* wild type (Col-0), the *rs31-1* loss-of-function mutant, and the RS31-ox line [38]. Transcript quantification was performed using StringTie (v2.2.1), with a reference GTF file containing pri-miRNA annotations (GFF2 format). Quantification was executed using the following parameters: stringtie Sam-ple.bam -p 4 -G primiRNA.gtf -e -b Sample/. Gene-level FPKM (Fragments Per Kilobase of transcript per Million mapped reads) values were extracted from the resulting t_data.ctab files for each replicate and aggregated using a custom R script. Differential expression between genotypes was assessed by performing two-sample t-tests on FPKM values, comparing wild type with both the mutant and overexpressor lines.

## Results

### Identification of pri-miRNA stem-loop interacting proteins via biotin-based pulldown

A growing number of RBPs have been implicated in the complex and dynamic regulation of plant miRNA biogenesis yet understanding the specific regulatory proteins that interact with distinct miRNA precursors remains insufficiently explored. To identify RBPs regulating the processing of miRNA precursors in Arabidopsis, we developed an RNA affinity pulldown based on biotinylated *in vitro* transcripts of pri-miRNA (*MIRNA*) stem-loops (*SL*). The experimental strategy is outlined in Fig. 1A. We chose the *SL* regions of two pri-miRNAs that are processed from loop-to-base and from base-to-loop, respectively. *Pri-miR159a* is processed from loop-to base in a sequential manner to release the mature miR159 [57, 58]. The template for *in vitro* transcription comprised 198 nucleotides downstream of the T7 RNA polymerase promoter (Additional file 1, Table S1, Additional file 3, Fig. S1A). In contrast, *pri-miR398b* is processed in base-to-loop direction to release the mature miR398. To include all structural determinants for accurate processing, the 134 nucleotide *MIR398b-SL* transcript contains the ∼15-nt lower stem region which specifies the position of the first DCL1 cleavage site (Additional file 1, Table S1, Additional Additional file 3, Fig. S1B). For RNA affinity pulldown, the biotinylated *MIR159a-SL* and *MIR398b-SL in vitro* transcripts were immobilized on streptavidin beads.

**Fig. 1.**
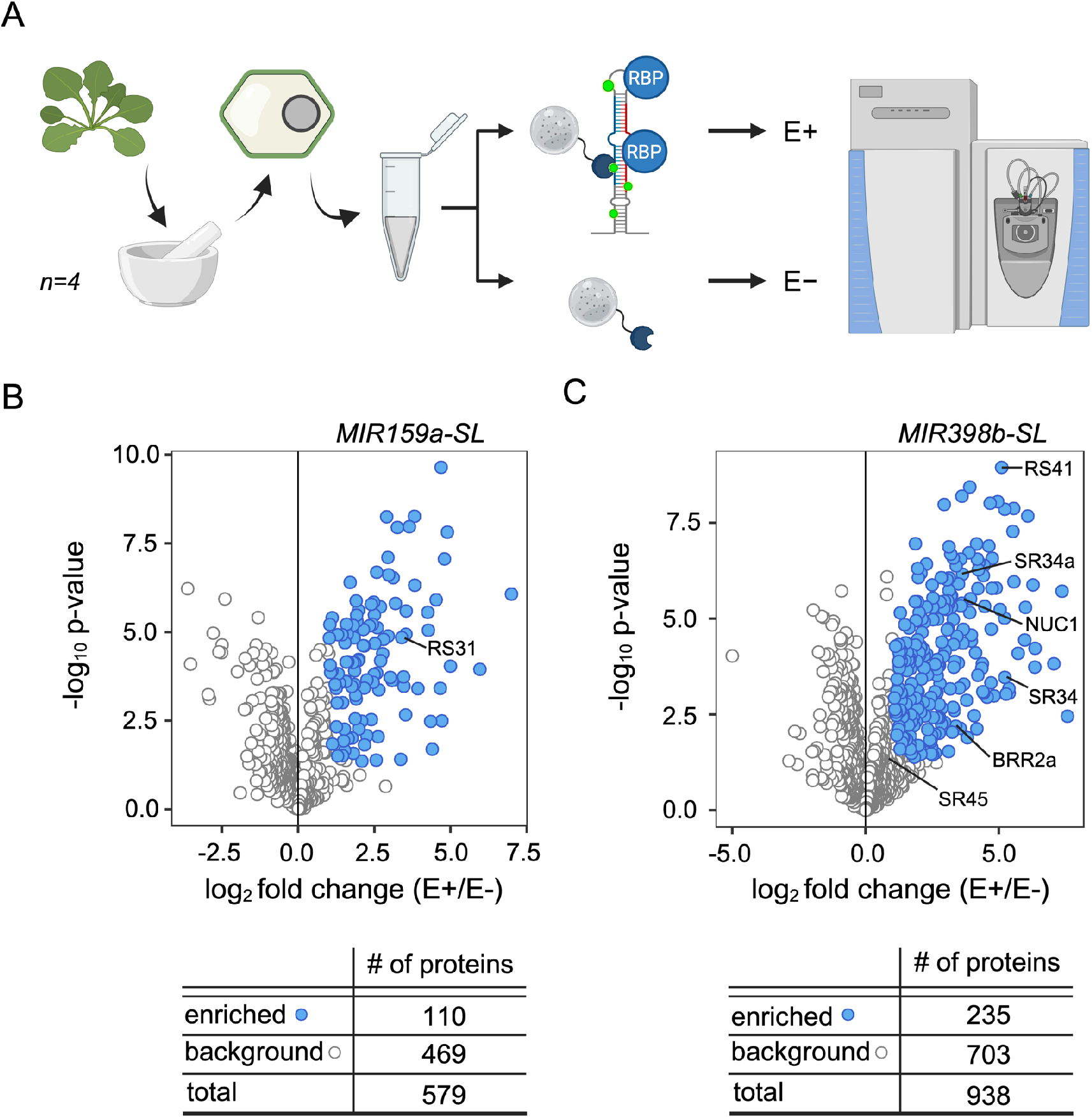
Biotin-based pulldown to identifies candidate proteins interacting with *MIRNA* stem-loops. **(A)** Experimental strategy: Nucleoplasmic protein extracts were prepared from four independent batches of Arabidopsis plants and incubated with biotinylated *MIRNA* stem-loops transcripts immobilized on streptavidin beads. Eluted proteins (E+) were identified by mass spectrometry and enrichment was determined relative to eluates from empty beads (E-) (Welch *t*-test, n=4). **(B-C)** Volcano plots show proteins recovered with the *MIR159a-SL* **(B)** and the *MIR398b-SL* **(C)**. Proteins with a log_2_ fold change > 1 and a p-value ≤ 0.05 (Welch *t*-test, n=4) are defined as interactors, whereas all other proteins are considered background. Proteins selected for further analysis are indicated.

As pri-miRNA processing occurs entirely in the nucleus in higher plants, both *MIR159a*-*SL* and *MIR398b-SL in vitro* transcripts were incubated with nuclear extracts to identify candidate interactors. Therefore, nuclei were isolated, and extracts enriched in nucleoplasmic proteins were prepared without lysing nuclei, as outlined in Additional file 3, Fig. S2A. The enrichment of nucleoplasmic proteins and the purity of the extracts was demonstrated by immunoblots using compartment-specific antibodies (Additional file 3, Fig. S2B) (Additional file 4, Fig. S18: Uncropped blots to Fig. S2B).

The bead immobilized *in vitro* transcripts of *MIR159a-SL* and *MIR398b-SL*, respectively, were then incubated with the nucleoplasmic proteins. Denaturing gel electrophoresis before the pulldown (input) and after the pulldown (supernatant and eluate) confirmed the integrity of the RNAs during the incubation (Additional file 3, Fig. S3A, B) (Additional file 4, Fig. S19: Uncropped gels to Fig. S2A, B). After extensive washing of the beads, bound proteins were heat-eluted in LDS sample buffer. Comparison of the proteins eluted from beads loaded with either *MIR159ab-SL* or *MIR398b-SL* (E+) and proteins eluted from empty beads (E-) by SDS-PAGE followed by silver staining confirmed the co-precipitation of proteins interacting with the *MIRNA-SLs* (Additional file 3, Fig S3C, D) (Additional file 4, Fig. S19: Uncropped gels to Fig. S2C, D).

The remainder of the eluates were subjected to MS in four biological replicates. Proteins which are significantly enriched relative to eluates from empty beads with a log_2_ fold change ≥ 1.0 and a p-value ≤ 0.05 were classified as interactors (Fig. 1B, C) (Additional file 5, Table S5 and S6). Proteins not meeting these criteria were not considered further as significantly enriched candidates. In the four preparative pulldowns for the *MIR159a-SL*, we retrieved 579 proteins. 110 of these were significantly enriched relative to proteins retrieved with empty beads (E-) (Fig. 1B, Additional file 5, Table S5). For the *MIR398b* stem-loop, 938 proteins were retrieved in total with 235 being significantly enriched (Fig. 1C, Additional file 5, Table S6).

### Categorization of the proteins interacting with the *MIRNA* stem-loops

To functionally categorize the proteins enriched in the pulldown assays, we conducted a systematic gene ontology (GO) term analysis of the interactors’ molecular identities. Among the candidate interactors of *MIR159a-SL* and *MIR398b-SL*, 52% and 60%, respectively, were annotated with RNA-associated molecular function GO terms (Fig. 2A, top). Accordingly, the interactors were also enriched for RNA-associated GO terms in Biological Process (BP) (Additional file 6, Table S7 and S8). This is consistent with potential functions as regulators of miRNA biogenesis. Importantly, two thirds of the interactors were present among the RNA binding proteomes (RBPome) in Arabidopsis determined by RNA interactome capture using oligo(dT) pull-down or purification based on phase extraction (Fig. 2A, bottom, Additional file 7, Table S9) [47, 59–64]. This provides independent support for RNA-binding activity of many of the identified proteins *in vivo*.

**Fig. 2.**
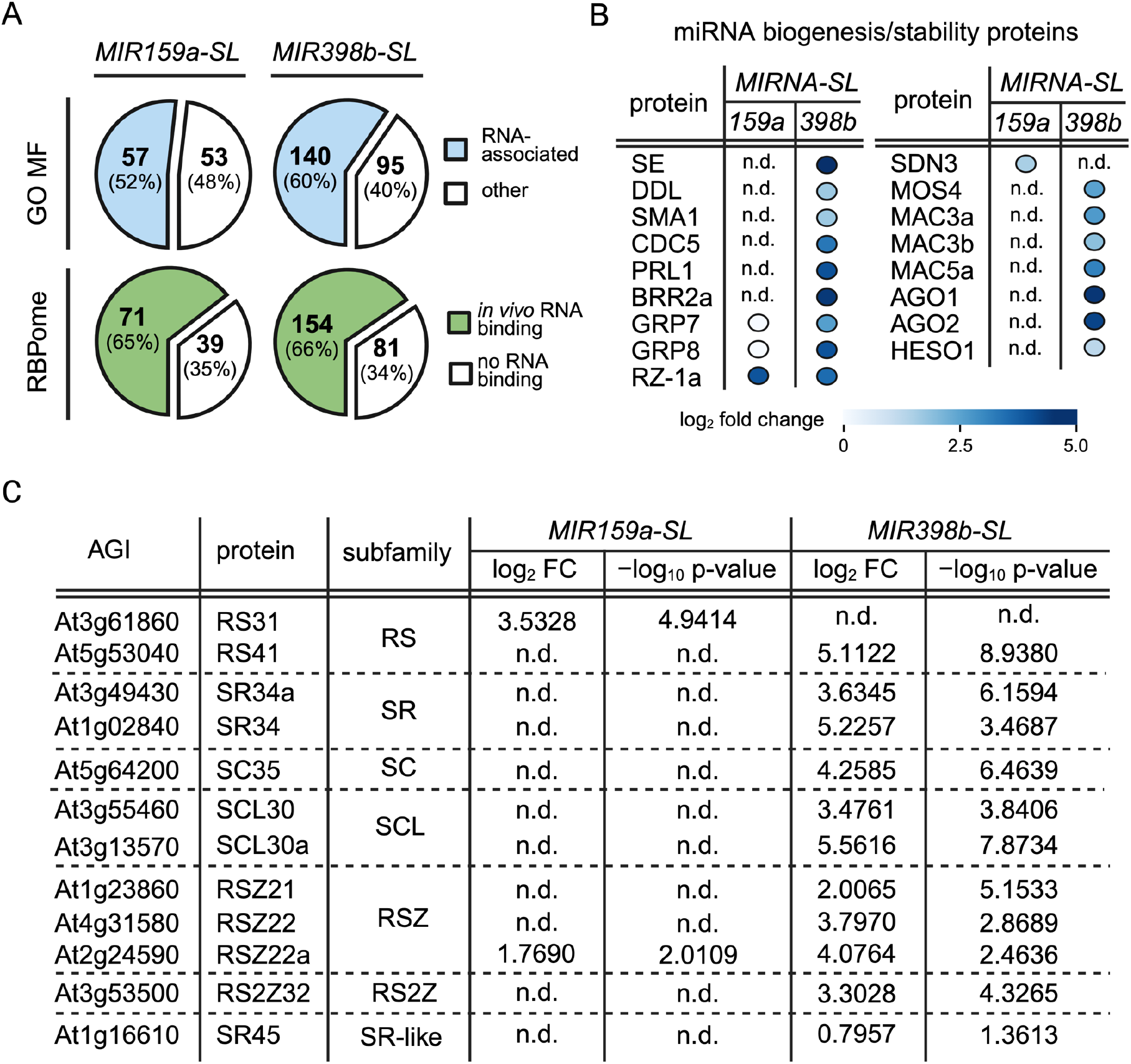
Functional classification and properties of candidate *MIRNA-SL* interactors. **(A)** Pie charts displaying the proportion of enriched proteins that are associated with RNA-binding-related molecular function (MF) Gene Ontology (GO) terms (top) and overlap with the Arabidopsis RNA-binding proteomes (RBPomes) determined by RNA interactome capture (bottom) for *MIR159a-SL* interactors and *MIR398b-SL* interactors. **(B)** Proteins associated with *MIRNA* processing/stability enriched in the pulldowns with *MIR159a-SL* and *MIR398b-SL*. **(C)** SR proteins identified as *MIR159a-SL* or *MIR398b-SL* interactors. SR-like protein SR45 was detected but did not meet the fold-change cutoff. n.d. – not detected.

Furthermore, among the protein domains enriched in the interactors characterized RNA-binding and nucleotide-binding domains are predominantly found (Additional file 8, Table S10 and S11).

To further validate our purification strategy, we tested whether we co-purified proteins known to be involved in miRNA biogenesis (Fig. 2B). For *MIR159a-SL*, no components of the miRNA core processing complex (defined as DCL1, HYL1, SE) were found. However, among the interactors were RZ-1a and SDN, proteins that have already been associated with miRNA biogenesis and stability, respectively [35, 65]. Additionally, the glycine-rich RNA binding proteins GRP7 and GRP8, for which direct *in vivo* binding to pri-*miR159a* has already been confirmed, are also among the identified, albeit not significantly enriched proteins [24].

For *MIR398b-SL*, SE of the core processing complex, AGO1, AGO2, and numerous proteins associated with pri-miRNA processing were significantly enriched (Fig. 2B). In particular, the BRR2a RNA helicase, a component of the U5 small nuclear ribonucleoprotein (U5 nRNP), recently shown to interact with pri-miRNAs, ensuring proper folding for processing was among the interactors [66]. Additionally, another component of U5 snRNP, GAMETOPHYTIC FACTOR 1 (GFA1, At1g06220), was enriched (Additional file 5, Table S6).

Notably, we identified several SR proteins. Although RS40 and RS41 have been implicated in pri-miRNA processing, the impact of other SR and SR-like proteins in pri-miRNA processing is not well understood (Fig. 2C). The SR protein family in plants is expanded compared to mammals. Arabidopsis harbors 18 SR proteins classified into six subfamilies and two SR-like proteins, SR45 and SR45a [67]. Pulldown of *MIR159a-SL* led to enrichment of RS31, a member of the plant-specific RS subfamily characterized by two RRMs (RS31, RS31a, RS40, and RS41). Furthermore, RSZ22a (RSZ subfamily with similarity to human SRSF7 harbouring a single RRM and one zinc knuckle) was also enriched.

In the pulldown with *MIR398b-SL*, RS41 (RS subfamily), and the SR subfamily members SR34 and SR34a (orthologous to human SRSF1 proteins with an RRM and a pseudo-RRM), were identified as interactors. We further detected enrichment of the RSZ subfamily proteins RSZ21, RSZ22, and RSZ22a, and the plant-specific RS2Z32 protein (RS2Z subfamily; single RRM and two zinc knuckles). Among single-RRM SR proteins, SC35 (an SRSF2 orthologue) and the plant-specific SCL30 and SCL30a were enriched. The SR-like protein At-SR45 showed a log_2_ fold change of 0.7957. As it did not meet our significance criterion, it was not classified as an interactor.

### Impact of interactors on pri-miRNA and mature miRNA levels

To assess whether the identified *MIR159a-SL* and *MIR398b-SL* interactors influence miRNA biogenesis, we first analyzed pri-miRNAs and mature miRNA levels in selected SR protein mutants, as several SR proteins have recently been implicated in this process (Fig. 3).

**Fig. 3.**
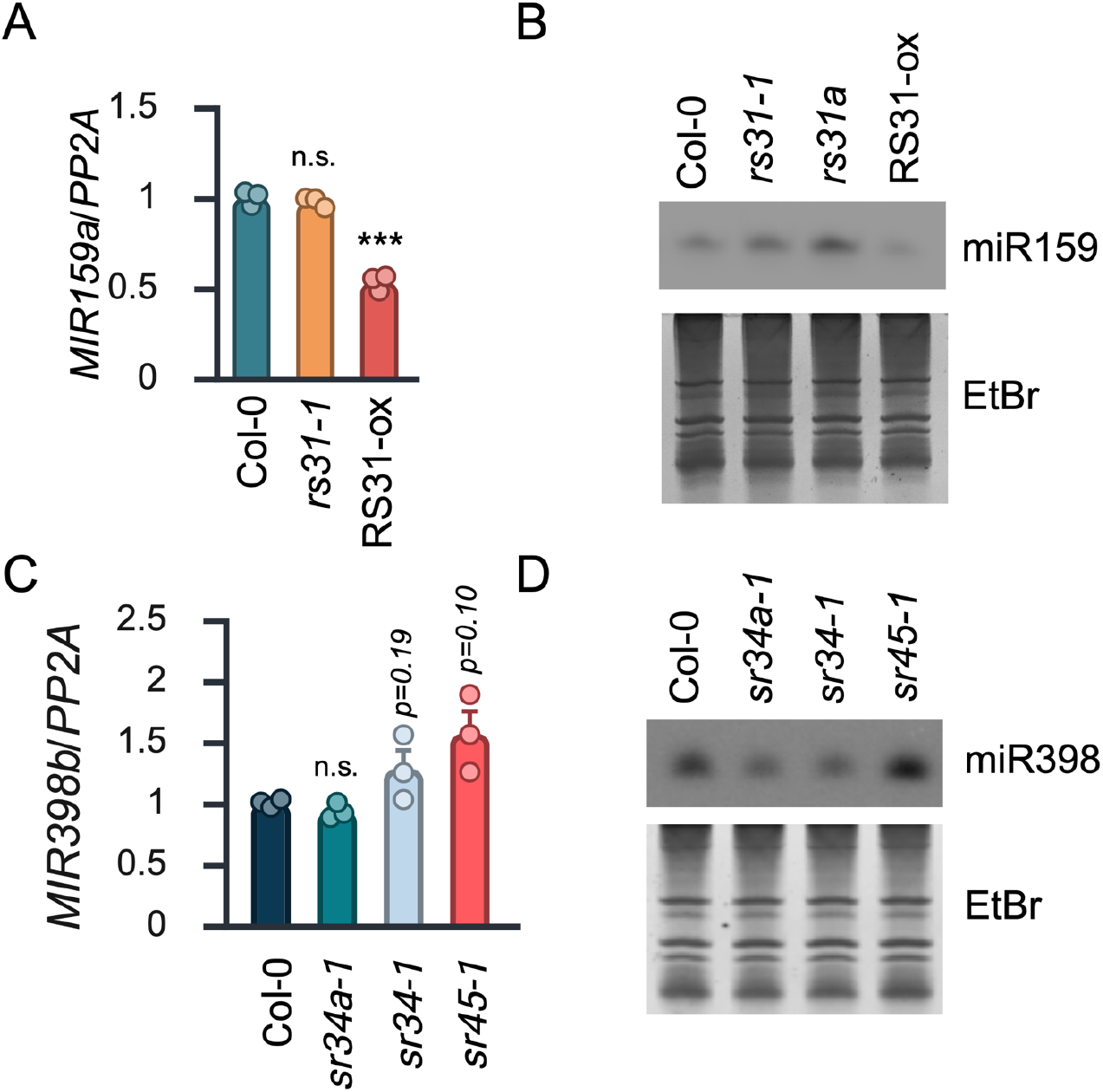
Impact of SR proteins on miR159 and miR398 levels. **(A)** RT-qPCR analysis of *MIR159a* levels in Col-0, *rs31-1*, *rs31a* and RS31-ox lines. **(B)** RNA gel blot analysis of miR159 in Col-0, *rs31-1*, *rs31a* and RS31-ox lines. Ethidium bromide staining of the gel serves as a landing control. **(C)** RT-qPCR analysis of *MIR398b* levels in Col-0, *sr34a-1*, *sr34-1,* and *sr45-1*. **(D)** RNA gel blot analysis of mature miR398 in Col-0, *sr34a-1*, *sr34-1,* and *sr45-1*. For RT-qPCR results, data are based on three biological replicates and shown as mean ± SD. Student’s *t*-test was performed to determine statistical significance (*** *p ≤* 0.001, ** *p ≤* 0.01, * *p ≤* 0.05, n.s., not significant).

For *MIR159a-SL*, RS31 was identified as an interactor. We analyzed the *rs31-1* mutant [37] and plants constitutively overexpressing RS31 (RS31-ox) [68]. As some peptides shared between the paralogs RS31 and RS31a and therefore cannot be unambiguously assigned by mass spectrometry, we also included the *rs31a* knockdown line carrying an artificial miRNA (amiR) construct against *RS31a* (amiR-31a-E2) [39].

*Pri-miR159a* levels were unchanged in *rs31-1* but significantly reduced in RS31-ox plants relative to wild type (Fig. 3A). By contrast, mature miR159 levels were significantly increased in the *rs31-1* mutant and the amiR *rs31a* knockdown line, whereas they were modestly but not significantly (p=0.09) decreased in RS31-ox (Fig. 3B, Additional file 3, Fig S4A) (Additional file 4, Fig. S20: Uncropped gels and blots to Fig. 3B and Fig. S4A), These results suggest an inverse relationship between RS31/RS31a abundance and mature miR159 accumulation. Because the amiR-*rs31a* construct is based on the *pri-miR159a* backbone, the resulting high level of *pri-miR159a* signal precluded reliable quantification of the endogenous *pri-miR159a* levels in the amiR-*rs31a* knockdown line (Additional file 3, Fig 4B) [39].

**Fig. 4.**
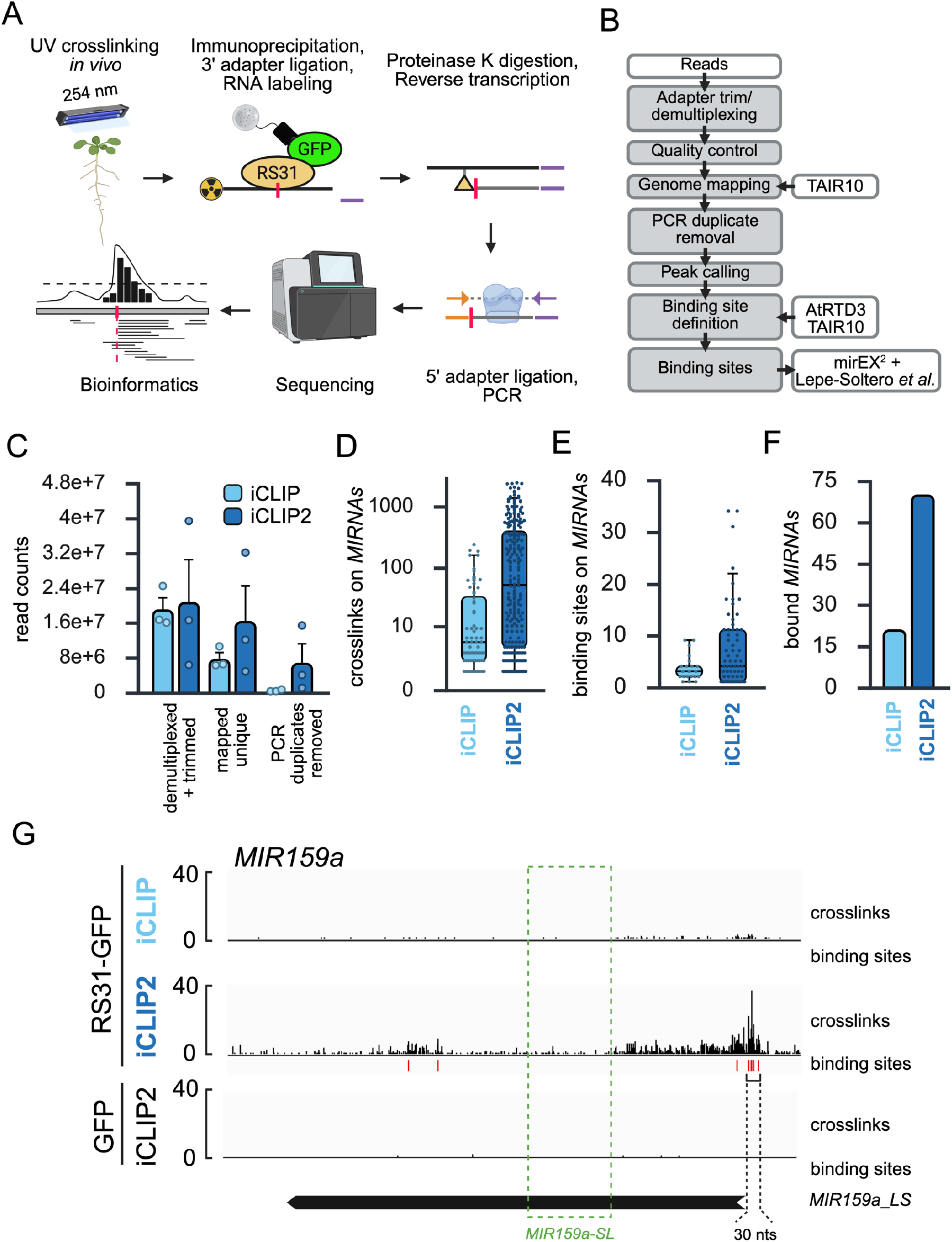
Plant iCLIP2 identifies RS31-GFP binding sites on *MIRNAs*. **(A)** Outline of the iCLIP2 procedure. **(B)** Bioinformatics workflow. **(C)** Number of reads after demultiplexing, adapter removal, trimming and quality control, mapped reads, uniquely mapped reads, and reads after deduplication. **(D, E)** Number of RS31-GFP crosslink and binding sites on *MIRNAs* in iCLIP and plant iCLIP2. **(F)** Number of RS31-GFP *MIRNA* targets in iCLIP and plant iCLIP2. **(G)** Crosslink sites and significant binding sites of RS31-GFP on *MIR159a* as determined by iCLIP and iCLIP2. For comparison, crosslink sites and significant binding sites of GFP are displayed.

We also mined available transcriptome data of *rs31-1* mutant and RS31-ox plants for differential expression of *MIRNAs* [38]. Because *MIRNAs* remain incompletely annotated, likely due to the highly variable length and poorly defined transcript boundaries, we generated a custom annotation file for putative *MIRNAs* by combining the annotation data from mirEX^2^ and Lepe-Soltero et al. [53, 54] (Additional file 2, Table S4).

Using this annotation file, we detected expression of 326 putative *MIRNA* transcripts across the data set. Of these, 207 were expressed in wild-type plants (FPKM ≥ 1), 210 in the *rs31-1* mutant, and 199 in RS31-ox (Additional file 9, Table S12). Pairwise comparisons of FPKM values revealed that, in the *rs31-1* mutant, four *MIRNA* transcripts showed significantly higher abundance and two showed significantly lower abundance compared to wild type at nominal significance threshold of p ≤ 0.05 (Additional file 3, Fig 4C, Additional file 9, Table S12). In RS31-ox plants, eight *MIRNA* transcripts showed higher abundance and 15 showed lower abundance relative to wild type using the same criterion (Additional file 3, Fig 4C, Additional file 9, Table S12). In contrast to the RT-qPCR analysis, *pri*-*miR159a* did not show a significant change in either *rs31-1* or RS31-ox in the RNA-seq data set. This may be due to the sequence similarity within the *MIR159* family and the resulting high variability of *MIR159a* FPKM values among the three wild-type replicates (Additional file 3, Fig 4E, Additional file 9, Table S12).

Next, we analyzed the level of *MIR398b* and mature miR398 in the respective mutants of the *MIR398b-SL* interactors SR34a and SR34. While the level of *pri-miR398b* was unchanged in both *sr34a-1* and *sr34-1* compared to the wild-type (Fig. 3C), the level of mature miR398 was moderately but significantly reduced in both mutants (Fig. 3D, Additional file 3, Fig. S5A) (Additional file 4, Fig. S21: Uncropped gels and blots to Fig. 3D and S5A).

Although SR45 did not meet our significance threshold as an interactor of *MIR398b-SL*, its direct *in vivo* binding to a miR398 precursor was previously confirmed via RNA immunoprecipitation and sequencing (RIP-seq) [69]. Consequently, we evaluated the levels of mature miR398 and *pri-miR398b* in *sr45-1* but found no significant differences compared to the wild type, despite slight increases in *pri-miR398b* (Fig. 3C) and miR398 levels (Fig. 3D, Additional file 3, Fig. S5A) (Additional file 4, Fig. S21: Uncropped gels and blots to Fig. 3D and S5A)..

The plant-specific RS subfamily protein RS41 was among the most strongly enriched *MIR398b-SL* interactors and showed one of the lowest p-values. It is known that RS41 and RS40 directly bind to a subset of miRNA precursors *in vivo* and act together with HIGH OSMOTIC GENE EXPRESSION 5 (HOS5) to facilitate the biogenesis of miRNAs [25]. To elucidate the effect of RS41 protein on miR398 expression, we employed the *rs40 rs41* double mutant owing to the significant homology and redundancy of both RS proteins as previously described [25]. Contrary to the unchanged level of *pri-miR398b* (Additional file 3, Fig. S5B), the level of mature miR398 was significantly reduced in the *rs40 rs41* double mutant compared to the wild-type (Additional file 3, Fig. S5C) (Additional file 4, Fig. S21: Uncropped gels and blots to Fig. S5C).

Together, these results indicate that distinct SR proteins influence the accumulation of selected mature miRNAs, often without corresponding changes in the levels of their cognate pri-miRNAs, consistent with functions at or downstream of pri-miRNA processing.

To further assess the functional relevance of identified interactors of *MIRNA-SLs* we selected additional proteins containing classical RNA-binding domains for analysis. Among the interactors of *MIR398b-SL*, we identified the helicase BRR2a. In the *brr2a-2* (*cäö*) mutant, in which threonine at position 895 is substituted by isoleucine *pri-miR398b* and mature miR398 levels were decreased, suggesting that BRR2a affects *pri-miR398b* and mature miR398, in addition to its previously described effects on other miRNAs [66] (Additional file 3, Fig. S6A-C) (Additional file 4, Fig. S22: Uncropped gels and blots to Fig. S6A-C). Furthermore, in the *nuc1-2* mutant defective in a nucleolin-like protein (NUC1), levels of *pri-miR398b* and of mature miR398 were significantly reduced (Additional file 3, Fig. S6A-C) (Additional file 4, Fig. S22: Uncropped gels and blots to Fig. S6A-C).

Because miRNA levels vary widely in response to different environmental stimuli [66], we tested whether BRR2A or NUC1 affects *pri-miR398b* and mature miR398 accumulation after exposure to salt and abscisic acid (ABA), a key hormone in plant stress responses. For *brr2a-2* and *nuc1-2*, the levels of *pri-miR398b* were stronger reduced compared to wild-type upon exposure to 200 mM NaCl than under control conditions (Additional file 3, Fig. S6D) (Additional file 4, Fig. S22: Uncropped gels and blots to Fig. S6D). Mature miR398 levels were also strongly reduced in *brr2a-2* and *nuc1-2* under salt stress (Additional file 3, Fig. S6E/F) (Additional file 4, Fig. S22: Uncropped gels and blots to Fig. S6E/F). In response to ABA, *pri-miR398b* levels were modestly, but not significantly reduced to 50% in *brr2a-2* (p = 0.12) whereas mature miR398 levels were not affected (Additional file 3, Fig. S6G-I) (Additional file 4, Fig. S22: Uncropped gels and blots to Fig. S6G-I). Similarly, neither *pri-miR398b* nor mature miR398 differed significantly between *nuc1-2* and wild-type upon ABA treatment. These results suggest that pri-miRNA-associated proteins, including BRR2A and NUC1, can modulate miRNA expression in response to changing environmental conditions, such as salt stress.

### Identification of RS31 binding sites on *MIRNAs in vivo*

To investigate the interaction between *MIRNA* transcripts and SR proteins *in vivo* and to test whether RS31 directly contacts pri-miRNAs, we employed iCLIP-based approaches (Fig. 4A). First, we re-analyzed our previously published RS31 iCLIP data set [68]. To identify RS31 binding to pri-miRNAs, our custom annotation file for potential *MIRNAs* (Additional file 2, Table S4) was integrated in the bioinformatics pipeline (Fig. 4B). The coordinates of the identified binding sites were then compared with the *MIRNA* annotation file and candidate RS31 bound pri-miRNAs were extracted. In this workflow, crosslink sites were first identified at nucleotide resolution, and binding sites were subsequently defined computationally from the crosslink-site distribution according to the established pipeline.

However, this methodology produced only limited evidence for RS31 binding to pri-miRNAs, as relatively few crosslinks were detected and, consequently, binding sites were identified at only a small number of *MIRNA* loci (Additional file 10, Table S13). This is likely due, at least in part, to the low abundance of *MIRNA* transcripts in Arabidopsis [27, 32]. We therefore performed RS31 iCLIP using our recently established more sensitive plant iCLIP2 protocol [32]. The protocol incorporates the greatly improved efficiency of library preparation originally introduced by König and colleagues [70].

Plants expressing the RS31-GFP fusion protein under control of its native promoter in the *rs31-1* background were subjected to UV crosslinking. RS31-GFP and associated RNAs were precipitated from the lysates using GFP-Trap^®^ beads. RNAs were radioactively labeled, and RNA-protein complexes were separated by denaturing gel electrophoresis, blotted onto a nitrocellulose membrane and subjected to autoradiography (Additional file 3, Fig. S7A) (Additional file 4, Fig. S23: Uncropped autoradiograms and gels to Fig. S7A). The indicated membrane regions were excised and subjected to library preparation for three biological replicates. Libraries were amplified by preparative PCR using nine cycles (Additional file, 3, Fig. S7B) (Additional file 4, Fig. S23: Uncropped autoradiograms and gels to Fig. S7B). As a control, we used transgenic plants expressing GFP alone under the control of the CaMV 35S promoter (Additional file 3, Fig. S7C/D) (Additional file 4, Fig. S24: Uncropped autoradiograms and gels to Fig. S7C/D). After sequencing, reads were processed according to our bioinformatics workflow supplemented with the *MIRNA* annotation file (Fig. 4B) [32].

Direct comparison between the previous iCLIP dataset with the plant iCLIP2 dataset showed a substantially higher average number of uniquely mapped reads after processing the raw reads. After removal of the PCR duplicates generated during library amplification, plant iCLIP2 retained an average of around 10 million reads, compared with 0.4 million reads in iCLIP (Fig. 4C, Additional file 10, Table S13). This was accompanied by almost one order of magnitude increase in the number of crosslink sites detected by plant iCLIP2 compared with iCLIP (Fig. 4D, Additional file 10, Table S14). Consequently, more binding sites on pri-miRNAs were identified using plant iCLIP2 (Fig. 4E, Additional file 10, Table S14 and S15). In total, our modified plant iCLIP2 pipeline identified 68 candidate RS31 pri-miRNA targets compared with 21 in the previous iCLIP dataset (Fig. 4F, Additional file 10, Table S14 and S15).

iCLIP has been shown to be biased to highly expressed transcripts [27]. To test whether we can also find binding sites on pri-miRNAs with low expression levels using our plant iCLIP2 protocol, we monitored expression levels of pri-miRNAs in the RNA-seq data of wild-type plants and compared those with the expression levels of bound pri-miRNAs [38]. Pri-miRNAs detected in iCLIP and plant iCLIP2 displayed similar high median levels, although in plant iCLIP2 more pri-miRNAs with lower expression levels were also detected (Additional file 3, Fig. S8). Overall, our improved plant iCLIP2 protocol enables more efficient identification of RBP-binding sites on short-lived pri-miRNAs than iCLIP.

As proof-of-concept, we checked whether *pri-miR159a* is among the plant iCLIP2 targets and thus directly bound by RS31 *in vivo.* The direct comparison between iCLIP and plant iCLIP2 displayed an increased number of *MIR159a* crosslink sites in our improved plant iCLIP2 data set (Fig. 4G). In contrast to iCLIP, three RS31 binding sites were identified on the annotated *MIR159a* using plant iCLIP2, confirming *pri-miR159a* as a direct target of RS31. Surprisingly, no binding sites could be called within the *MIR159s-SL* region, instead two binding sites were identified upstream and one downstream of the stem-loop. Nevertheless, individual crosslink sites were identified within the stem-loop region, although they did not meet the criteria for binding-site calling in our pipeline. This suggests that RS31 may contact the *pri-miR159a* stem-loop transiently or at low frequency *in vivo*, which could explain why RS31 co-purified with the *MIR159a-SL* in the *in vitro* RNA affinity pulldown. In addition, we observed strong enrichment of crosslink sites and four RS31 binding sites in a region just 30 nucleotides upstream of the annotated *MIR159a* sequence. This suggests that the current annotation of *MIR159a* may be incomplete at the 5’ end. Thus, RS31 appears to bind most prominently upstream of the annotated *MIR159a* stem-loop, while also making weaker or transient contacts within the stem-loop region.

Next, we analyzed the sequence characteristics of the identified binding sites. Given the limited number of binding sites on *MIRNAs*, we performed motif discovery using all transcriptome-wide binding sites to obtain a more robust result (Additional file 11, Table S16). Consistent with earlier findings indicating that RS31 preferentially binds to CU-rich motifs, STREME identified a similar CU-rich motif in the plant iCLIP2 datasets derived from RS31-GFP plants (Additional file 3, Fig. S9A) [69]. A parallel search for k-mer motifs (trimers) enriched within the binding sites confirmed enrichment of CU-rich sequence combinations (Additional file 3, Fig. S9B). The presence of CU-rich motifs within the *MIR159a-SL* is consistent with the possibility that RS31 transiently contacts the *MIR159a* stem-loop region, which may explain its co-purification in the *in vitro* RNA-affinity pulldown (Additional file 3, Fig. S9C).

Cross-referencing differentially abundant *MIRNAs* with the plant iCLIP2 data showed that two RS31-bound pri-miRNAs were moderately, but significantly, up-regulated in *rs31-1 vs.* wild-type (Additional file 3, Fig. S10A). Conversely, six RS31-bound pri-miRNAs were significantly down-regulated and three bound pri-miRNAs were up-regulated in RS31-ox compared with wild-type (Additional file 3, Fig. S10 B). Among these, *pri-miR156c* showed reciprocal changes in *rs31-1* and RS31-ox. A closer look showed nine consecutive RS31 binding sites upstream of the *MIR156c* stem-loop at the 5’ end of the annotated precursor (Additional file 3, Fig. S10C). In addition, there are eight closely spaced binding sites in region of the 3’ base of the annotated stem-loop.

Interestingly, *pri-miR398c* was among the RS31-bound pri-miRNA transcripts that were significantly down-regulated transcripts in RS31-ox (Additional file 3, Fig. S10B). Like *pri-miR398c*, *pri-miR398b* was also down-regulated in RS31-ox, although not significantly (p=0.07). The annotated *MIR398b* and *MIR398c* stem-loop sequences are almost identical. However, RS31 was not co-purifed in the *MIR398b-SL* affinity pull-down. Closer examination of the RS31-binding site distribution on *pri-miR398c* (Additional file 3, Fig. S10 D) and *pri-miR398b* (Additional file 3, Fig. S11A) revealed that binding sites were exclusively localized down-stream of the annotated stem-loops and there were no CU-rich RS31-binding motifs within the annotated *pri-miR398c* and *pri-miR398b* stem-loops. Notably, many additional binding sites were located downstream of the annotated *MIR398c* and *MIR398b* regions, suggesting that annotations of these pri-miRNA transcripts may be incomplete. Three consecutive binding sites were located close to the base of the *pri-miR398b* stem-loop (Additional file 3, Fig. S11A/B). According to the short base-to-loop processing mechanism, this region is important for proper processing of *pri-miR398b* by DCL1 [21]. An imperfectly paired lower stem of ∼15 nucleotides below the miRNA/miRNA* duplex, followed by an internal loop, determines the initial DCL1 pri-miRNA cleavage site. Subsequent cleavage of the pre-miRNA at a distance of ∼21 nucleotides from the first cut then releases the miRNA/miRNA* duplex. To test whether altered RS31 abundance affects mature miR398 accumulation, we performed small RNA gel blots. Mature miR398 levels showed reciprocal but no significant trend, with increased levels in *rs31-1* and reduced levels in RS31-ox (Additional file 3, Fig. S11C) (Additional file 4, Fig. S26: Uncropped gels and blots to Fig. S11C). We further validated the significant decrease of *pri-miR398c* in RS31-ox and the absence of significant change in the *rs31-1* mutant by RT-qPCR (Additional file 3, Fig. S11D). In accordance with the RNA-seq data, *pri-miR398b* levels were reduced in RS31-ox, albeit not significantly (p=0.07) (Additional file 3, Fig. S11D).

Together, these data show that plant iCLIP2 significantly improves the *in vivo* detection of crosslink sites and binding sites on pri-miRNA transcripts *in vivo* and enables identification of a substantially broader set of *MIRNA* transcript targets than the previous iCLIP protocol.

### Identification of SR34a binding sites on *MIRNAs in vivo*

To identify *in vivo* binding of SR34a to pri-miRNAs, we performed plant iCLIP2 on transgenic plants expressing SR34a-GFP under control of its native promoter in the *sr34a-1* mutant background. This line had previously been used to determine the transcriptome-wide binding landscape of SR34a in germinating seeds [34]. Two-week old plants grown in 12 h light/12 h dark cycles at 20°C were subjected to UV crosslinking. SR34a-GFP and associated RNAs were immunoprecipitated from crude lysates using GFP-Trap^®^ beads followed by radioactive labeling and denaturing gel electrophoresis (Fig. 5A) (Additional file 4, Fig. S25: Uncropped autoradiograms and gels to Fig. 5A). The radioactive smear above the expected size of the SR34a-GFP fusion protein, which was strongly reduced after treatment with a high concentration RNase I, indicates efficient RNA-protein crosslinking and recovery of SR34a-GFP-associated RNAs.

**Fig. 5.**
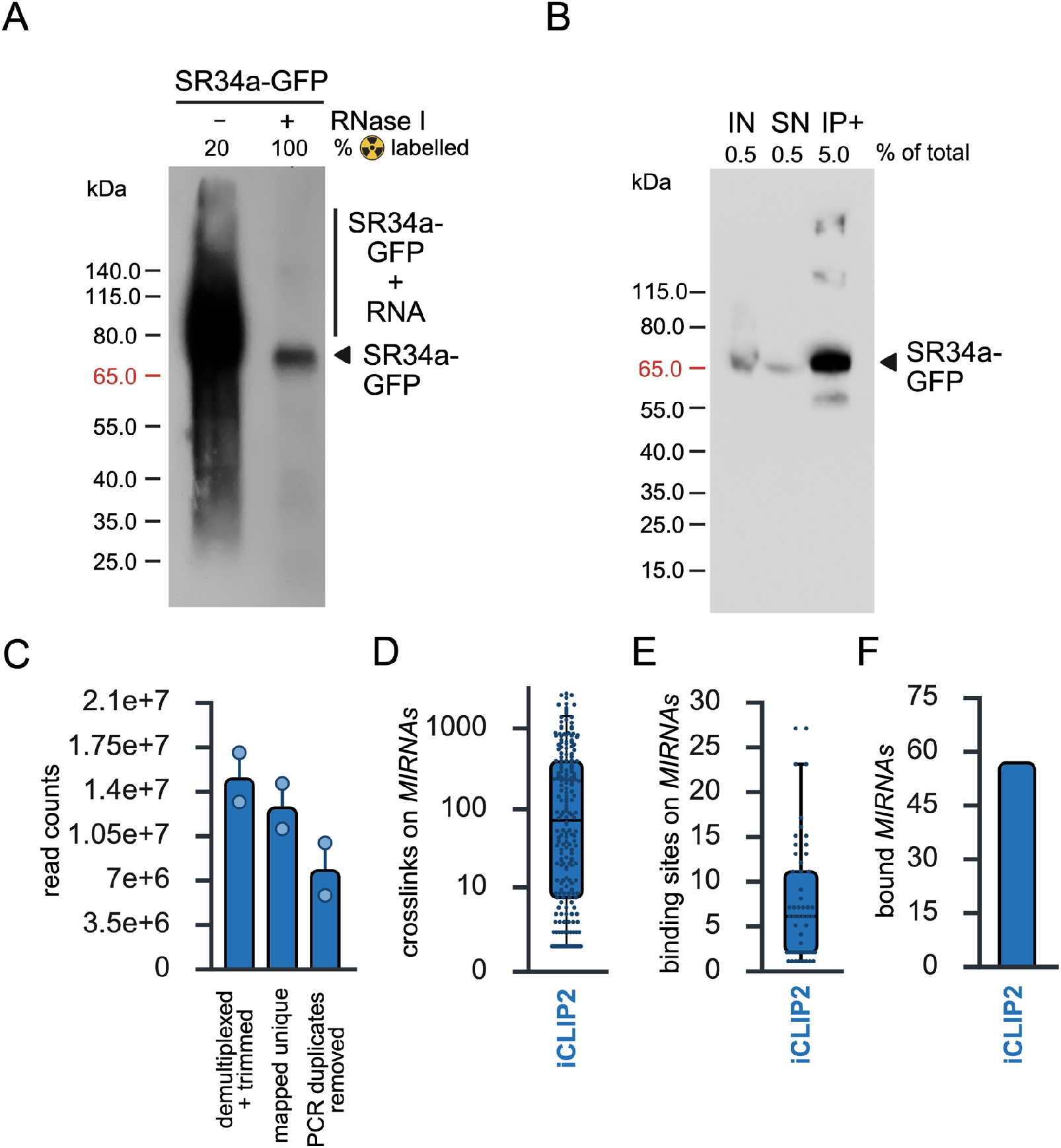
Plant iCLIP2 of SR34a-GFP. **(A)** Autoradiogram of RNA-protein complexes from *SR34a::SR34a-GFP sr34a-1* plants after UV crosslinking and immunoprecipitation with GFP-Trap^®^ beads. Samples qwere treated with RNase I as indicated. The radioactive smear above the expected size of SR34a-GFP corresponds to SR34a-GFP crosslinked to RNA. The arrowhead marks the expected position of SR34a-GFP after RNase treatment. **(B)** Immunoblot analysis of *SR34a::SR34a-GFP sr34a-*plants. Aliquots of the input fraction (IN, 0.5%), supernatant (SN, 0.5%) and the immunoprecipitated fraction (IP+, 5%) were separated by NuPAGE and probed with the α-GFP antibody. **(C)** Number of reads retained after demultiplexing, adapter removal, trimming and quality control, mapped reads, uniquely mapped reads, and reads after removal of PCR duplicates. **(D, E)** Number of SR34a-GFP crosslink sites (D) and binding sites (E) identified on *MIRNAs* by iCLIP and plant iCLIP2. **(F)** Number of SR34a-GFP *MIRNA* targets identified by plant iCLIP2.

In parallel, an immunoblot analysis confirmed efficient precipitation of the SR34a-GFP from the lysate (Fig. 5B) (Additional file 4, Fig. S25: Uncropped autoradiograms and gels to Fig. 5B). We then performed plant iCLIP2 in two biological replicates (Additional file 3, Fig. S12A) (Additional file 4, Fig. S27: Uncropped autoradiograms and gels to Fig. S12A). After PCR-cycle optimization, libraries were amplified using ten cycles (Additional file 3, Fig. S12B) (Additional file 4, Fig. S27: Uncropped gels to Fig. S12B).

Plant iCLIP2 statistics are shown in Fig. 5C-F and Additional file 12, Table S17. Overall, the number of reads retained after processing and removal of PCR duplicates was in a range similar range to that obtained for RS31-GFP. This led to the identification of a total of 65,775 crosslink sites (Fig. 5D, Additional file 12, Table S18) and 412 binding sites (Fig. 5E, Additional file 12, Table S18) distributed across *57* pri-miRNAs bound by SR34a-GFP (Fig. 5F, Additional file 12, Table S19). Motif using STREME and k-mer enrichment confirmed the predominant GCU motif previously identified for SR34a-GFP in germinating seeds (Additional file 3, Fig. S13A,B) [34].

Because SR34a co-purified with the *MIR398b-SL* in the affinity pulldown, we first checked whether SR34a-GFP also binds to the *pri-miR398b in vivo*. Indeed, a total of four binding sites were identified on the annotated *MIR398b* region, all located downstream of the annotated stem-loop (Fig. 6). An additional binding site was found just four nucleotides downstream of the annotated *MIR398b* region. A similar binding pattern with a binding site located downstream of the stem-loop and outside the annotated precursor region, was observed for *MIR398c* (Additional file 3, Fig. S14A). Importantly, although mature miR398 levels were significantly reduced in *sr34a-1* (Fig. 3D, Additional file 3, Fig. S5A) (Additional file 4, Fig. S21: Uncropped gels and blots to Fig. 3D and S5A), the level of *pri-miR398b* and *pri-miR398c* levels did not differ significantly between *sr34a-1* and wild-type (Fig. 3C, Additional file 3, Fig. S14B).

**Fig. 6.**
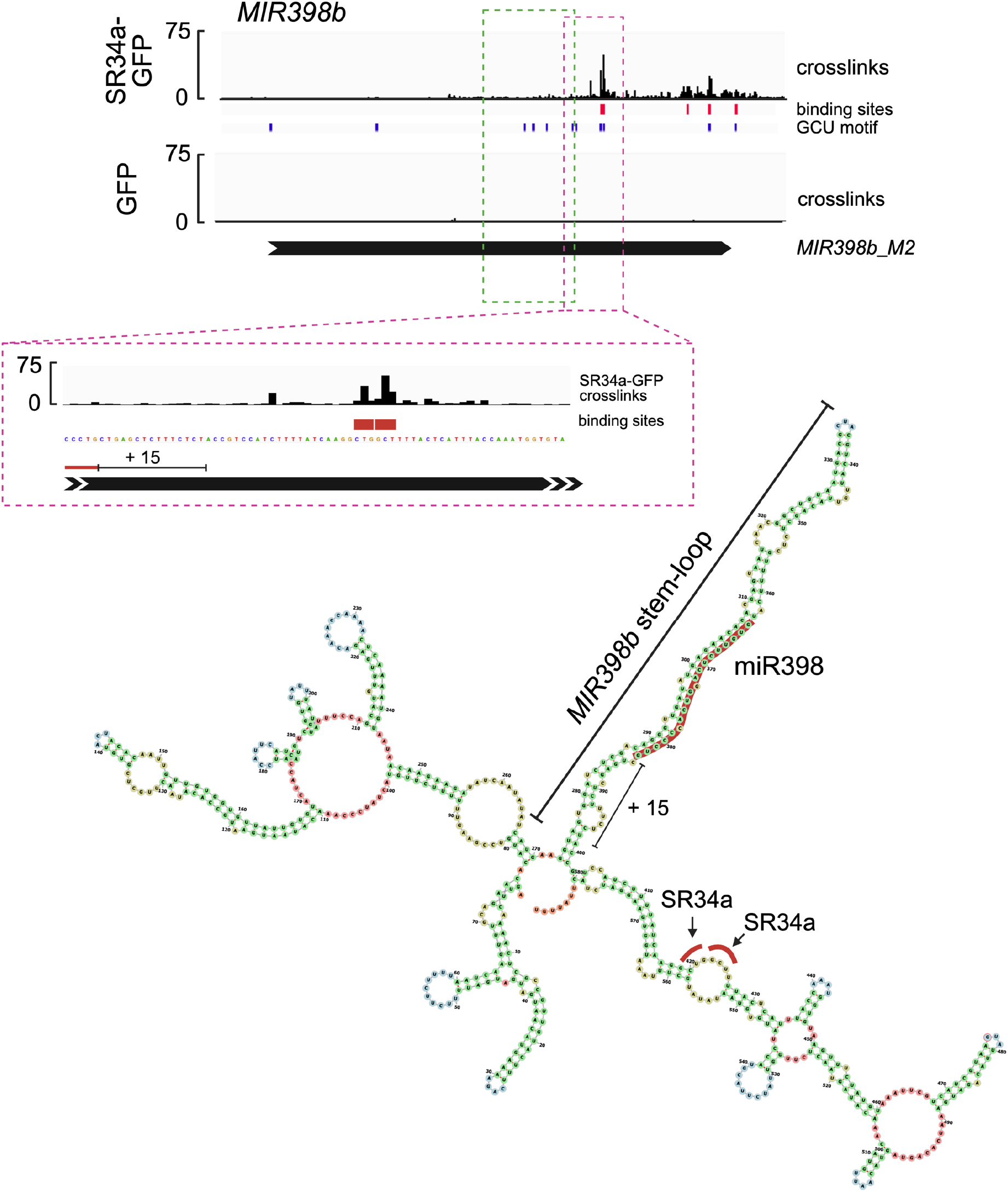
SR34a-GFP binding sites on *pri-miR398b*. Top: Crosslink sites and significant SR34a-GFP binding sites on *MIR398b* as determined by plant iCLIP2. The GFP control track is shown for comparison. Red boxes mark identified binding sites, and blue boxes indicate GCU motifs within the annotated *MIR398b_M2* region shown as a black arrow. The green dashed box marks the region of the annotated *MIR398b* stem-loop region. The magenta dashed box highlights SR34a-GFP binding sites downstream of the base of the *MIR398b* stem-loop region. Bottom: *In silico* predicted secondary structure of *MIR398b*. Red bars indicate the binding sites near the base of the stem-loop. The +15 marks the region important for correct processing of *pri-miR398b*.

### Comparative analysis of RS31-GFP and SR34a-GFP *MIRNA* targets

Comparison of the pri-miRNAs directly bound by RS31-GFP and SR34a-GFP revealed 48 common pri-miRNA targets for the two SR proteins, corresponding to 70.6 % of RS31-GFP targets and 84.2 % of SR34a-GFP targets (Additional file 3, Fig. S15). This suggests that RS31 and SR34a associate with a largely shared subset of pri-miRNA transcripts. In contrast, 20 pri-miRNAs were identified exclusively as RS31-GFP targets and nine pri-miRNAs as SR34a-GFP targets, corresponding to 29.4 % and 15.7 % of their respective target sites.

Moro and colleagues have described loop-to-base (LTB) and base-to-loop (BTL) processing modes for more than one hundred evolutionarily conserved miRNA precursors. About one third of these precursors are processed in the LTB or sequential (s)LTB mode (i.e., *MIR779*, *MIR159* and *MIR319* families). Among pri-miRNAs processed in the BTL direction, members of the *MIR169* family and *MIR402* are processed in the sequential (s)BTL mode. The predominant processing mode was known for 31 of the 77 pri-miRNA transcripts identified as targets of either RS31-GFP or SR34a-GFP (Additional file 3, Fig. S15). Notably, neither RS31 nor SR34a showed an obvious preference for binding pri-miRNAs of a specific processing type, as all processing types are represented in expected proportions.

Next, we asked whether RS31 and SR34a preferentially bind specific regions of the miRNA precursors. For this analysis, the annotated *MIRNAs* were divides into regions upstream and downstream of the stem-loop, the stem-loop itself, and a base region defined as 25 nucleotides upstream and downstream of the stem-loop base (Additional file 3, Fig. S16A). Because the number of crosslinks and consequently the number of identified binding sites can correlate with the pri-miRNA abundance and may not directly reflect the binding affinity, we scored whether at least one binding site was present in each region rather than counting the total number of binding sites per region. Thus, individual pri-miRNAs could be assigned to more than one region. As previously described, RS31 binding was enriched towards the 5’ region of transcripts [38]. Consistently, RS31 binding was detected upstream of the stem-loop for 46 of 68 RS31-bound pri-miRNAs, downstream of the stem-loop for 25, within the stem-loop for 15, and at the base of the stem-loop for 10 (Additional file 3, Fig. S16B). A separate analysis of pri-miRNAs processed by BTL or LTB mechanisms showed a comparable distribution, although the number of pri-miRNAs in these subsets was lower (Additional file 3, Fig. S16B). A similar regional distribution was observed for SR34a (Additional file 3, Fig. S16B). Thus, despite belonging to different SR protein subfamilies, RS31 and SR34a showed broadly similar regional binding distributions on pri-miRNAs. For both RS31 and SR34a binding was detected most frequently upstream of the stem-loop, suggesting that these SR proteins preferentially associate with flanking regions of pri-miRNA transcripts rather than with the stem-loop itself, regardless of processing type. However, no clearly defined general pattern of pri-miRNA binding was apparent.

Among the common evolutionarily conserved pri-miRNA targets that are processed in the BTL direction was *pri-miR167a*. In a region approximately 500-1000 nucleotides downstream of the annotated *MIR167a* stem-loop, one RS31 and seven SR34a binding sites were detected (Additional file 3, Fig. S17A). In addition, four RS31 and five SR34a binding sites were located in an approximately 40-nucleotide CU-rich region only 15 nucleotides upstream of the base of the stem-loop. Strikingly, RS31 and SR34a binding sites partially overlapped in this region.

In contrast, SR34a and RS31 showed distinct binding patterns on the BTL-processed *pri-miR171b* and *pri-miR396b* (Additional file 3, Fig. S17B,C). For *pri-miR171b,* only SR34a showed a significant binding site at the base of the stem-loop, whereas significant RS31 binding sites were located mainly at the 5’ end of the annotated precursor. Conversely, for pri-*miR396b,* significant binding sites in the stem-loop base region were detected only for RS31.

Overall, RS31 and SR34a showed a substantially overlapping set of pri-MIRNA targets, indicating that these SR proteins associate with a largely shared subset of pri-miRNA transcripts. However, neither protein showed an obvious preference for pri-miRNAs with a specific processing mode. Instead, binding by both proteins was detected most frequently upstream of the stem-loop, whereas binding within the stem-loop or at the stem-loop base was less common. Individual examples further showed that RS31 and SR34a can bind either overlapping or distinct regions of the same precursor. Thus, RS31 and SR34a appear to associate preferentially with pri-miRNA flanking regions, but their precise binding patterns are target-specific.

## Discussion

MiRNAs are essential post-transcriptional regulators that modulate gene expression by affecting mRNA stability and translation. By targeting transcription factors and other regulatory proteins, miRNAs can contribute to fine-tuned rather than binary regulation, thereby supporting robustness and adaptability in gene expression. In this context, RBPs involved in miRNA processing are likely to add specificity and regulatory versatility to miRNA production.

Here, we established a biotin-based pulldown strategy to identify candidate RBPs interacting with pri-miRNA SLs. The stem-loops of *pri-miR159a* processed from loop-to-base and *pri-miR398b* processed from base-to-loop were selected as representative models.

Using this approach, we identified 110 significantly enriched proteins for *MIR159a-SL* and 235 for *MIR398b-SL* relative to an empty bead control, which we define as candidate interactors. The suitability of the approach is supported by the recovery of pri-miRNA processing components and RBPs previously implicated in pri-miRNA processing. In addition, the presence of several identified proteins in current Arabidopsis RNA interactomes is consistent with RNA association *in vivo*. Nevertheless, enrichment in pulldown assays should be interpreted cautiously, as it does not by itself establish direct regulatory activity *in vivo*. Notably, the number of interactors identified for *MIR398b-SL* was considerably higher than in our previous pulldown using the cleavage-deficient CRISPR/Cas nuclease Cys4* [35].

Among the enriched proteins, several RRM domain-containing proteins belonged to the SR-rich protein family. Because some mammalian SR proteins have been implicated in the regulation of miRNA biogenesis [71–73], we first examined whether RS31 and its close paralog RS31a influence the abundance of the bound precursor *pri-miR159a* and the corresponding mature miR159. Mature miR159 levels were increased in the *rs31-1* mutant and in the *rs31a* knockdown line, whereas they were reduced in RS31-ox plants. These reciprocal changes are consistent with a negative effect of RS31 on mature miR159 accumulation. However, *pri-miR159a* levels remained largely unchanged in the *rs31-1* mutant, indicating that precursor abundance and mature miRNA accumulation may not be affected in a strictly parallel manner. Transcriptome-wide datasets from *rs31-1* and RS31-ox plants further suggest that RS31 may influence pri-miRNA abundance more broadly, as additional pri-miRNAs were differentially expressed upon RS31 misexpression.

A related pattern emerged for the SR subfamily proteins SR34a and SR34, both of which were purified with *MIR398b-SL*. SR34a and SR34 are homologues of the animal SR protein SRSF1, which modulates the processing of specific miRNAs in mammalian cells [72]. Although *pri-miR398b* levels were unchanged in *sr34a-1* and *sr34-1*, mature miR398 levels were slightly reduced in both mutants. This observation suggests that SR34a and SR34 may contribute to miR398 accumulation, although the modest effect size warrants a cautious interpretation.

By contrast, no significant differences in steady-state levels of either *pri-miR398b* or mature miR398 were detected in *sr45-1* mutants, which is consistent with the non-significant enrichment of SR45 protein in the pulldown. However, previous RIP-seq experiments have shown that SR45 binds to *pri-miR398c in vivo* [69]. Thus, a regulatory influence of SR45 on miR398 expression cannot be excluded, particularly if such an effect is context-dependent or specific to particular *MIR398* family members.

Previous work showed that RS40 and RS41 affect the processing of a subset of pri-miRNAs through direct binding [25]. Although miR398 was not previously identified as differentially regulated by RS40 and RS41, our data suggest that mature miR398 expression is altered in *rs40 rs41* mutants. This discrepancy may reflect differences in growth conditions or harvest time, especially because miR398 expression is diurnally regulated and reaches maximum expression at dusk [74].

Beyond SR proteins, the RNA helicase BRR2a was also recovered in the *MIR398b-SL* pulldown. The significantly reduced *pri-miR398b* and miR398 levels observed in *brr2a-2* mutants are consistent with previous evidence that BRR2a modulates pri-miRNA secondary structure to improve the efficiency and fidelity of processing from pri-miRNA to mature miRNA [66]. Loss of BRR2a function may therefore impair the structural suitability of multiple pri-miRNAs and reduce miRNA accumulation.

NUC1, a nucleolin-like nucleolar protein, promotes early pre-rRNA processing and ribosome biogenesis in plants. Loss of NUC1 in *nuc1-2* mutants alters rRNA precursor accumulation and causes nucleolar disorganization accompanied by developmental defects [75]. The co-purification of NUC1 with *MIR398b-SL*, together with the significantly reduced levels of *pri-miR398b* and mature miR398 in *nuc1-2* mutants, suggests that NUC1 may also contribute to miR398 biogenesis. The reduction in miR398 levels in *brr2a-2* and *nuc1-2* mutants was more pronounced under salt stress than under regular growth conditions. This finding suggests that regulatory effects that appear weak or difficult to detect under standard conditions may become more evident when plants experience specific environmental stimuli.

Across several RBP mutants, the observed differences in steady-state pri-miRNA or mature miRNA abundance were relatively small, as has also been reported in previous studies [17, 24, 25]. Such modest changes are not unexpected for miRNAs, which often act as modulators of gene expression. Moreover, analyses of whole plant tissue may obscure effects that occur only in specific cell types, and functional redundancy among related proteins may further limit the detection of changes in individual mutants, as observed for RS40 and RS41 [25]. These considerations should be taken into account when interpreting the magnitude of the observed effects.

Pulldown-based approaches have also been used in other organisms to identify RBPs associated with miRNA precursors. In mammals, for example, a global inventory of RBPs interacting with 72 pre-miRNA hairpins was assembled using pulldown with biotinylated antisense oligonucleotides across eleven cell lines, leading to the identification of 180 RBPs implicated in miRNA biogenesis [76]. Such inventories provide valuable candidate regulators, but they do not resolve the precise binding sites through which individual RBPs may influence processing.

Mapping the binding sites of RBPs that are not part of the core processing machinery is therefore important for distinguishing direct from indirect regulatory effects, identifying sequence or structural features that contribute to substrate recognition, and clarifying how individual RBPs may modulate the processing of specific pri-miRNAs. To improve the detection of RBP binding sites on lowly expressed *MIRNAs*, we modified the iCLIP library strategy and developed an optimized plant iCLIP2 workflow. Overall, these optimizations improved the efficiency of library preparation, as indicated by the increased number of uniquely mapped reads after removal of PCR duplicates. To identify RBP binding on poorly annotated *MIRNAs*, we also established a new *MIRNA* annotation file and combined it with the optimized plant iCLIP2 workflow. This strategy enabled the identification of more reproducible binding sites corresponding to RBP interaction sites on pri-miRNA transcripts *in vivo*. However, despite the improved library preparation efficiency, detection remained biased toward relatively highly expressed *MIRNAs*. Consequently, *MIRNA* binding targets are likely still underestimated.

Using this approach, we identified 68 pri-miRNA targets for RS31-GFP and 57 pri-miRNA targets for SR34a-GFP. Among these targets, significant binding sites were detected on *pri-miR159a* for RS31-GFP and on *pri-miR398b* for SR34a-GFP. These findings support direct *in vivo* binding of RS31 to *pri-miR159a* and SR34a to *pri-miR398b* and strengthen the interpretation that the biotin-based *in vitro* RNA pulldown can identify RBPs with regulatory relevance for miRNA biogenesis. At the same time, further functional analyses are required to determine how individual binding events influence precursor processing.

Further analysis of RS31-GFP and SR34a-GFP showed that some identified binding sites are located in sequence regions important for processing pri-miRNAs into mature miRNAs, including sequence and structural features proximal to the miRNA/miRNA* duplex [21]. In base-to-loop processed pri-miRNAs, the miRNA/miRNA* duplex contains a lower stem of approximately 15 nucleotides followed by a substantial bulge, and these lower-stem features are important for accurate processing. Members of the *MIR156* and *MIR398* families in particular exhibited RS31-GFP binding sites close to the lower stem of the respective pri-miRNAs and showed differential expression in RS31 gain- and loss-of-function mutants. Together, these observations are consistent with a model in which selected RBPs bind near processing-relevant structural regions and thereby contribute to the regulation of specific miRNA precursors.

The binding-site analyses further indicate that RS31- and SR34a-associated sites are not confined to the stem-loop regions themselves but are also detected prominently in sequences upstream and downstream of annotated pri-miRNA stem-loops. This observation also supports the need for extended *MIRNA* annotations when analyzing RBP binding to miRNA precursors. This broader distribution suggests that the regulatory contribution of these RBPs may extend beyond direct effects on the processing of the miRNA/miRNA* duplex within miRNA precursors. However, the functional relevance of binding in these flanking regions remains to be determined. In addition, the comparatively limited detection of binding sites within paired stem regions should be interpreted with caution, because UV crosslinking is generally less efficient in double-stranded RNA regions. Thus, reduced recovery of plant iCLIP2 signals in paired regions does not necessarily exclude interactions with such structures.

A large fraction of pri-miRNA targets was shared between RS31 and SR34a, with 48 common targets corresponding to 70.6 % of RS31-bound and 84.2 % of SR34a-bound pri-miRNAs. This extensive overlap raises the possibility that RS31 and SR34a contribute to combinatorial or partially redundant regulation of common miRNA precursors. The occurrence of overlapping or closely positioned binding sites is also consistent with the possibility of competitive binding at selected regions. Nevertheless, whether such competition occurs *in vivo* and whether it has a relevant effect on miRNA biogenesis remains unresolved. Addressing this question will require higher-order mutant combinations and targeted functional analyses that can distinguish individual, redundant, and potentially competitive contributions of these RBPs.

## Conclusion

This study combines biotin-based pri-miRNA stem-loop pulldown with optimized plant iCLIP2 to identify candidate RBPs associated with plant miRNA precursors and validate selected interactions *in vivo*. The results reveal that selected SR-rich proteins, together with BRR2a and NUC1, can influence pri-miRNA and mature miRNA accumulation, often in a precursor-specific and context-dependent manner. Direct *in vivo* binding of RS31 to *pri-miR159a* and of SR34a to *pri-miR398b* supports the functional relevance of the pulldown approach and suggests that selected RBPs act near processing-relevant structural regions or adjacent flanking sequences. Overall, these findings expand the repertoire of plant miRNA biogenesis regulators and provide a framework for dissecting combinatorial RBP functions in miRNA biogenesis.

## Abbreviations

GO: Gene ontology
iCLIP: individual-nucleotide resolution crosslinking and immunoprecipitation
*MIRNA*: pri-miRNA
miRNA: microRNA
MS: mass spectrometry
nt: nucleotide
RBP: RNA-binding protein
SR: serine-arginine rich

## Supplementary Information

The online version contains supplementary material.

## Additional files

**Additional file 1:**

**Table S1:** gBlocks^®^ and oligonucleotides used for template generation and *in vitro* transcription. The given sequence maps of gBlock_T7-Csy4-MIR159a-SL and gBlock_T7-MIR398b-SL and plasmid map of pUC19 contain detailed information on regulatory elements, restriction sites and primer binding sites.

**Table S2:** Biotinylated DNA-oligos used for miRNA detection by Northern blotting.

**Table S3:** Oligonucleotides used for RT-qPCR.

**Additional file 2:**

**Table S4:** Genomic location of *MIRNAs* compiled from two joined annotation sources which have been used in this study. Coordinates were unified by selecting the longest annotation (in base pairs) among overlapping entries. Annotation sources include mirEX^2^ (M2) [53], Lepe-Soltero et al. (LS) [54], and entries exclusive to either mirEX^2^ (oM2) or Lepe-Soltero et al. (oLS). Each *MIRNA* entry includes chromosomal location, strand orientation, and source attribution.

**Additional file 3:**

**Figure S1:** Secondary structure prediction of **(A)** *MIR159a-SL* and (B) *MIR398b-SL* using RNAfold from the Vienna RNA Websuite [36].

**Figure S2:** Characterization of nucleoplasmic extracts. **(A)** Scheme of the experimental procedure. **(B)** Immunoblot analysis of total lysate (T), cytoplascmic fraction (C), and nuclear extract (N). The indicated amount of the protein was separated by SDS-PAGE and the blots were probed with antibodies against nucleoplasmic SERRATE (top), cytoplasmic UGPase (middle) and chromatin-associated Histon H3 (bottom). The sizes of the molecular weight standards are indicated.

**Figure S3:** Control of the RNA and protein fractions of the RNA affinity pulldowns (PD). The integrity of the *in vitro* transcribed **(A)** *MIR159a*-SL, and **(B)** *MIR398b-SL* in the input (IN), supernatant (SN) and the eluate fraction (E+) was confirmed by urea polyacrylamide gel electrophoreses. Proteins recovered by pulldown with immobilized **(C)** *MIR159a-SL* and (D) *MIR398b-SL* were analysed by SDS gel electrophoresis and silver staining (E+). Proteins eluted from empty beads (E-) served as control.

**Figure S4:** MiRNA levels in *rs31-1* mutant and RS31-ox lines. **(A)** Quantification of miR159 levels in Col-0 wild-type, *rs31-1*, *rs31a* and RS31-ox lines determined by small RNA gel blots (cf. Fig. 3B). MiR159 levels were quantified using ImageJ and levels are expressed relative to wild-type. **(B)** Level of *MIR159a* in *rs31a* determined by RT-qPCR. Shown are the mean ±SD of three biological replicates. A Student’s *t*-test was performed to determine statistical significance (*** *p ≤* 0.001, ** *p ≤* 0.01, * *p ≤* 0.05, n.s., not significant). Transcriptome-wide identification of differentially expressed *MIRNAs* in either **(C)** *rs31-1* or **(D)** RS31-ox compared to wild-type [38]. **(E)** Level of *MIR159a* in *rs31-1* or RS31-ox relative to wild-type.

**Figure S5:** MiRNA levels in *sr34a-1, sr34-1, sr45-1* and *rs40/rs41* mutants. **(A)** Quantification of miR398 levels in Col-0 wild-type, *sr34a-1*, *sr34-1* and *sr45-1* mutant lines determined by small RNA gel blots (cf. Fig. 3D). MiR398 levels were quantified using ImageJ and levels are expressed relative to wild-type. **(B)** Level of *MIR398b* in *rs40/rs41* (*rs40/41*) determined by RT-qPCR. **(C)** A small RNA gel blot of the *rs40/rs41* (*rs40/rs41*) mutant and the corresponding wild-type plants was hybridized with an anti-miR398 probe (left). Ethidium bromide (EtBr) staining served as a loading control. MiR398 signals were quantified using ImageJ (right). Shown are the mean ±SD of three biological replicates. A Student’s *t*-test was performed to determine statistical significance (*** *p ≤* 0.001, ** *p ≤* 0.01, * *p ≤* 0.05, n.s., not significant).

**Figure S6:** MiRNA levels in *brr2a-2* and *nuc1-2* under non-stress and abiotic stress conditions. The levels of *MIR398b* in A, D, and G and mature miR398 in B, E, and H were determined by RT-qPCR and small RNA gel blots, respectively, under (A-C) untreated (non-stress control) conditions or upon treatment with (D-F) 200 mM sodium chloride (NaCl) or (G-I) 100 µM abscisic acid (ABA) for 48 hours (h). Small RNA blots were hybridized with an anti-miR398 probe and ethidium bromide (EtBr) staining served as a loading control. MiR398 signals in C, F, and I were quantified using ImageJ. Shown are the mean ±SD of three biological replicates. A Student’s *t*-test was performed to determine statistical significance (*** *p ≤* 0.001, ** *p ≤* 0.01, * *p ≤* 0.05, n.s., not significant).

**Figure S7:** Plant iCLIP2 of RS31-GFP. **(A)** Autoradiograms of RNA-protein complexes from *RS31::RS31-GFP rs31-1* plants after UV crosslinking and immunoprecipitation in three replicates. For technical reasons, the sample of the first replicate was split into 2 lanes after elution in LDS sample buffer. The regions containing the crosslinked RNAs used for library generation are indicated (red dashed boxes). A control treatment with high RNase I was previously described [45]. **(B)** Gel electrophoresis of amplified plant iCLIP2 cDNA libraries for RS31-GFP with increasing PCR cycles. P3Solexa and P5Solexa denote primers used for library amplification. **(C)** Autoradiogram of RNA-protein complexes from *35S::GFP* control plants after UV crosslinking and immunoprecipitation in two replicates. **(D)** Gel electrophoresis of amplified *35S::GFP* control libraries with increasing PCR cycles.

**Figure S8:** Shown is the average expression level (FPKM) of all *MIRNAs* expressed in Col-0 wild-type (FPKM ≥ 0) compared to the average expression level of bound *MIRNAs* identified for RS31-GFP by iCLIP or plant iCLIP2.

**Figure S9:** Sequence motifs enriched at the RS31-GFP binding sites. **(A)** Significantly enriched motifs determined by STREME (left). Density distribution of the identified binding motif around the binding site (right). The centre position (zero) corresponds to the binding site peak. **(B)** Scatterplot displaying trimer counts (x-axis) and trimer z-scores (y-axis). Trimers in the upper right corner are enriched in comparison to a random background correction model. Letters in upper case mark the peak position of the binding site. **(C)** Crosslinks and significant binding sites (red boxes) of RS31-GFP on *MIR159a* as determined by plant iCLIP2. The blue boxes mark the position of the identified CU-rich binding motifs CUCC and CUUC in the region of the annotated *MIR159a_LS* (black arrow). The green dashed box marks the region of the annotated *MIR159a* stem-loop (*SL)*. Blue arrows highlight the CU-rich binding motifs within the *MIR159a-SL*.

**Figure S10:** Transcriptome-wide identification of RS31-GFP bound and differentially expressed *MIRNAs* in either **(A)** *rs31-1* or **(B)** RS31-ox compared to wild-type. Crosslinks and significant binding sites of RS31-GFP on **(C)** *MIR156c* and **(D)** *MIR398c* as determined by plant iCLIP2. Blue boxes mark the position of the identified CU-rich binding motifs CUCC and CUUC in the region of the annotated *MIRNAs* (black arrow). The green dashed box marks the region of the annotated *MIRNA* stem-loops (*SL*).

**Figure S11: (A)** Crosslinks and significant binding sites (red boxes) of RS31-GFP on *MIR398b* as determined by plant iCLIP2. Blue boxes mark the position of the identified CU-rich binding motifs CUCC and CUUC in the region of the annotated *MIR398b_M2* (black arrow). The green dashed box marks the region of the annotated *MIR398b* stem-loop (*SL)*. The purple dashed box focusses on the RS31-GFP binding sites downstream of the base of the *MIR398b-SL*. **(B)** *In silico* predicted secondary structure of *MIR398b*. Red bars indicate the binding sites near the base of the stem-loop highlighted in (A). +15 denotes a region important for correct processing of *MIR398b*. **(C)** Level of *MIR398b* and *MIR398c* in *rs31-1* and RS31-ox compared to wild-type determined by RT-qPCR. **(D)** Levels of mature miR398 in *rs31-1* and RS31-ox compared to wild-type. After small RNA blots, levels were quantified using ImageJ and levels are expressed relative to wild-type. Shown are the mean ±SD of three biological replicates. A Student’s *t*-test was performed to determine statistical significance (*** *p ≤* 0.001, ** *p ≤* 0.01, * *p ≤* 0.05, n.s., not significant).

**Figure S12:** Plant iCLIP2 of SR34a-GFP. **(A)** Autoradiograms of RNA-protein complexes from *SR34a::SR34a-GFP sr34a-1* after UV crosslinking and immunoprecipitation in two replicates. The regions containing the crosslinked RNAs used for library generation are indicated (red dashed boxes). **(B)** Gel electrophoresis of amplified plant iCLIP2 cDNA libraries for SR34a-GFP with increasing PCR cycles. P3Solexa and P5Solexa denote primers used for library amplification.

**Figure S13:** Sequence motifs enriched at SR34a-GFP binding sites. **(A)** Significantly enriched motifs determined by STREME (left). Density distribution of the identified binding motif around the binding site (right). The centre position (zero) corresponds to the binding site peak. **(B)** Scatterplot displaying trimer counts (x-axis) and trimer z-scores (y-axis). Trimers in the upper right corner are enriched in comparison to a random background correction model. Letters in upper case mark the peak position of the binding site.

**Figure S14: (A)** Crosslinks and significant binding sites of SR34a-GFP on *MIR398c* as determined by plant iCLIP2. Blue boxes mark the position of the identified CCU binding motif in the region of the annotated *MIR398c_M2* (black arrow). The track displaying crosslinks of *35S::GFP* plants are given as a control. The green dashed box marks the region of the annotated *MIR398c* stem-loop (*SL)*. **(B)** RT-qPCR analysis of miR398 in Col-0 wild-type, *sr34a-1,* and *sr34-1*. Shown are the mean ±SD of three biological replicates. A Student’s *t*-test was performed to determine statistical significance (*** *p ≤* 0.001, ** *p ≤* 0.01, * *p ≤* 0.05, n.s., not significant).

**Figure S15:** Comparison of the *MIRNAs* directly bound by RS31-GFP and/or SR34a-GFP *in vivo*. Evolutionarily conserved *MIRNAs* for which a processing mechanism has been described are labelled accordingly (see figure legend).

**Figure S16:** Distribution of binding sites across RS31-GFP and SR34a-GFP *MIRNA* targets. **(A)** Annotated *MIRNAs* were divided into defined regions upstream (blue) and downstream (green) of the stem-loop, the stem-loop itself (black) and a base region (orange). **(B)** Distribution of binding sites on all *MIRNAs* bound RS31-GFP (left) or those, which were either known to be processed in base-to-loop (middle) or loop-to-base direction (right). **(C)** Distribution of SR34a-GFP *MIRNA* binding sites as described for RS31-GFP. Absolute numbers of *MIRNAs* are given in brackets.

**Figure S17:** Crosslinks and significant binding sites (red boxes) of RS31-GFP and SR34a-GFP on *MIRNAs* as determined by plant iCLIP2. Blue boxes mark the position of the identified binding motifs for RS31-GFP and SR34a-GFP. The green dashed box marks the region of the annotated *MIRNA* stem-loops (*SL)*. **(A)** The purple dashed box focusses on the RS31-GFP binding sites downstream of the base of the *MIR167a-SL*. The *in silico* predicted secondary structure of *MIR167a* is shown at the bottom. Red bars indicate RS31-GFP and SR34a-GFP binding sites near the base of the stem-loop highlighted above. +15 denotes a region important for correct processing of *MIR398b*. RS31-GFP and SR34a-GFP binding sites on the annotated **(B)** *MIR171b*, **(C)** *MIR396b,* and **(D)** *MIR157c*.

**Additional file 4:**

**Figure S18:** Uncropped blots to Figure S2B.

**Figure S19:** Uncropped gels to Figure S3A-D.

**Figure S20:** Uncropped gels and blots to Figure 3B and S4A.

**Figure S21:** Uncropped gels and blots to Figure 3D and S5A/C.

**Figure S22:** Uncropped blots and gels to Figure S6A-I.

**Figure S23:** Uncropped autoradiograms and gels to Figure S7A/B.

**Figure S24:** Uncropped autoradiograms and gels to Figure S7C/D

**Figure S25:** Uncropped autoradiogram and gel to Figure 5A/B.

**Figure S26:** Uncropped blots and gels to Figure S11C.

**Figure S27:** Uncropped autoradiograms to Figure S12.

**Additional file 5:**

**Table S5:** Mass spectrometry (MS) data of biotin-based *MIR159a-SL* affinity pulldown.

**Table S6:** Mass spectrometry (MS) data of biotin-based *MIR398b-SL* affinity pulldown.

**Additional file 6:**

**Table S7:** Gene ontology (GO) term analysis of *MIR159a-SL* interactors.

**Table S8:** Gene ontology (GO) term analysis of *MIR398b-SL* interactors.

**Additional file 7:**

**Table S9:** Overlap of proteins enriched in RNA affinity pulldowns with RNA binding proteomes.

**Additional file 8:**

**Table S10:** Protein domain enrichment of *MIR159a-SL* interactors.

**Table S11:** Protein domain enrichment of *MIR398b-SL* interactors.

**Additional file 9:**

**Table S12:** Differential expression analysis of *MIRNAs* in *rs31-1* and RS31-ox compared to wild-type.

**Additional file 10:**

**Table S13:** Plant iCLIP2 read statistics of RS31-GFP replicates after each processing step.

**Table S14:** Crosslinks and binding site coverage of RS31-GFP plant iCLIP2 on *MIRNAs*.

**Table S15:** Binding sites of RS31-GFP on annotated *MIRNAs*.

**Additional file 11:**

**Table S16:** Transcriptome-wide binding sites of RS31-GFP.

**Additional file 12:**

**Table S17:** Plant iCLIP2 read statistics of SR34a-GFP replicates after each processing step.

**Table S18:** Crosslinks and binding site coverage of SR34a-GFP plant iCLIP2 on *MIRNAs*.

**Table S19:** Binding sites of SR34a-GFP on annotated *MIRNAs*.

## Acknowledgement

We thank K. Neudorf for expert technical assistance, Lars Hennig and Claudia Köhler for providing the *brr2a-2* (*cäö*) mutant, A.S.N. Reddy for providing *sr45-1* mutant, Julio Saez-Vasquez for providing *nuc1-2* mutant. We acknowledge support for the publication costs by the Open Access Publication Fund of Bielefeld University and the Deutsche Forschungsgemeinschaft (DFG).

## Authorś contributions

V.B. and J.L. performed RNA pulldowns. A.W. performed small RNA gel blots, RT-qPCRs and bioinformatics analysis of RNA pulldowns. M.S. and F.B. performed MS analysis. M.R. performed immunoblotting and provided advice on RNA pulldowns. T.K. performed small RNA gel blots, RT-qPCRs, iCLIP and plant iCLIP2 experiments. M.L. performed bioinformatics analysis of iCLIP and assembled the corresponding supplementary tables. T.L., P.D. and M.K. provided mutants. M.K. provided RNA-seq data and expertise for data evaluation. T.K. and D.S. designed the experimental concepts and acquired funding. T.K. and D.S. wrote the manuscript. T.K. and A.W. prepared the figures. All authors approved the manuscript.

## Availability of data and materials

Plant iCLIP2 reads have been submitted to Sequence Read Archive (SRA) under the BioProject accession No.: PRJNA1478861.

## Ethics approval and consent to participate

Not applicable

## Consent for publication

Not applicable

