## Additional file 3 for "Identification and binding-site mapping of RNA-binding proteins interacting with microRNA precursors in Arabidopsis"

#### Figure S1

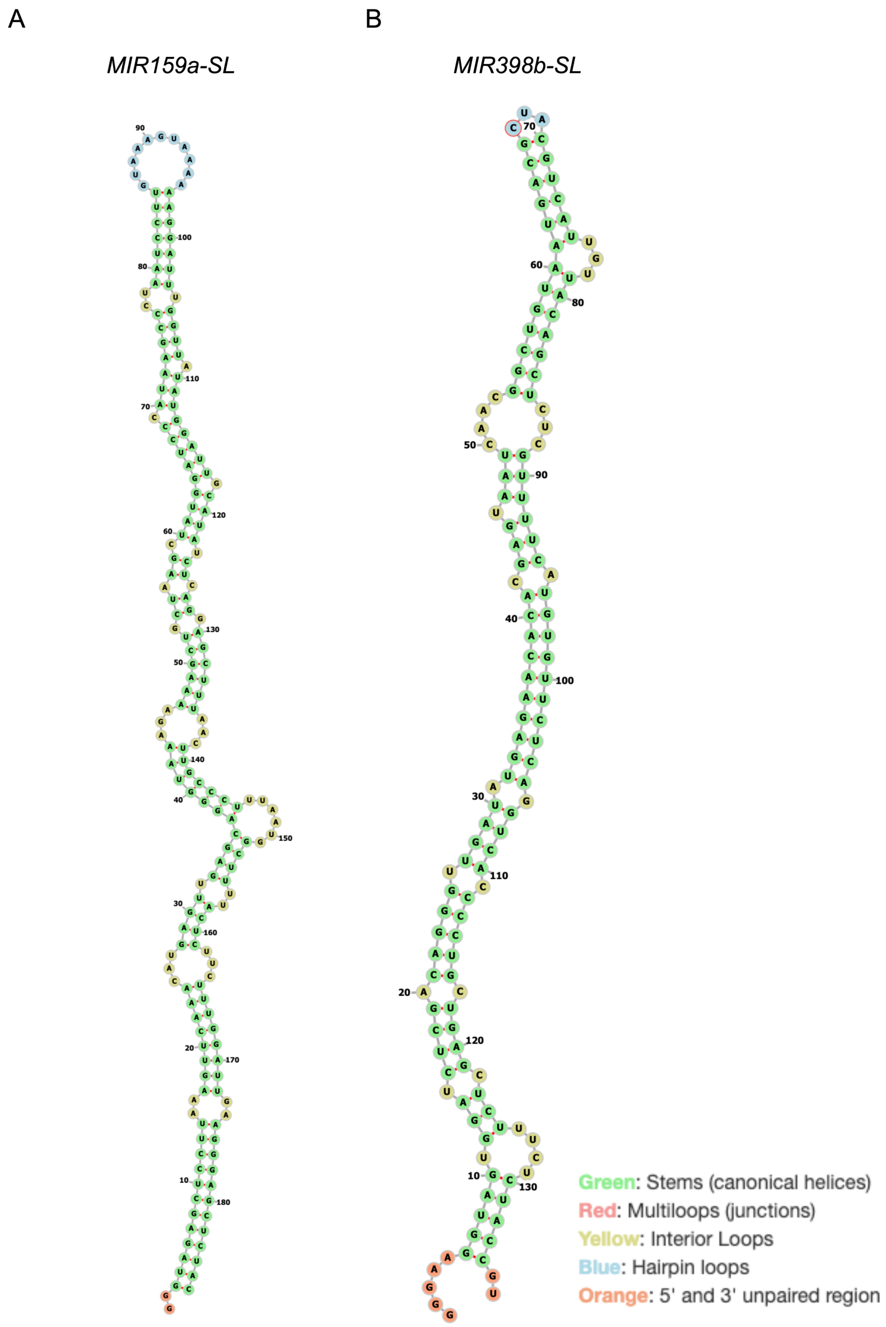


**Figure S1:** Secondary structure prediction of (A) *MIR159a-SL* and (B) *MIR398b-SL* using RNAfold from the Vienna RNA Websuite [36].

#### Figure S2

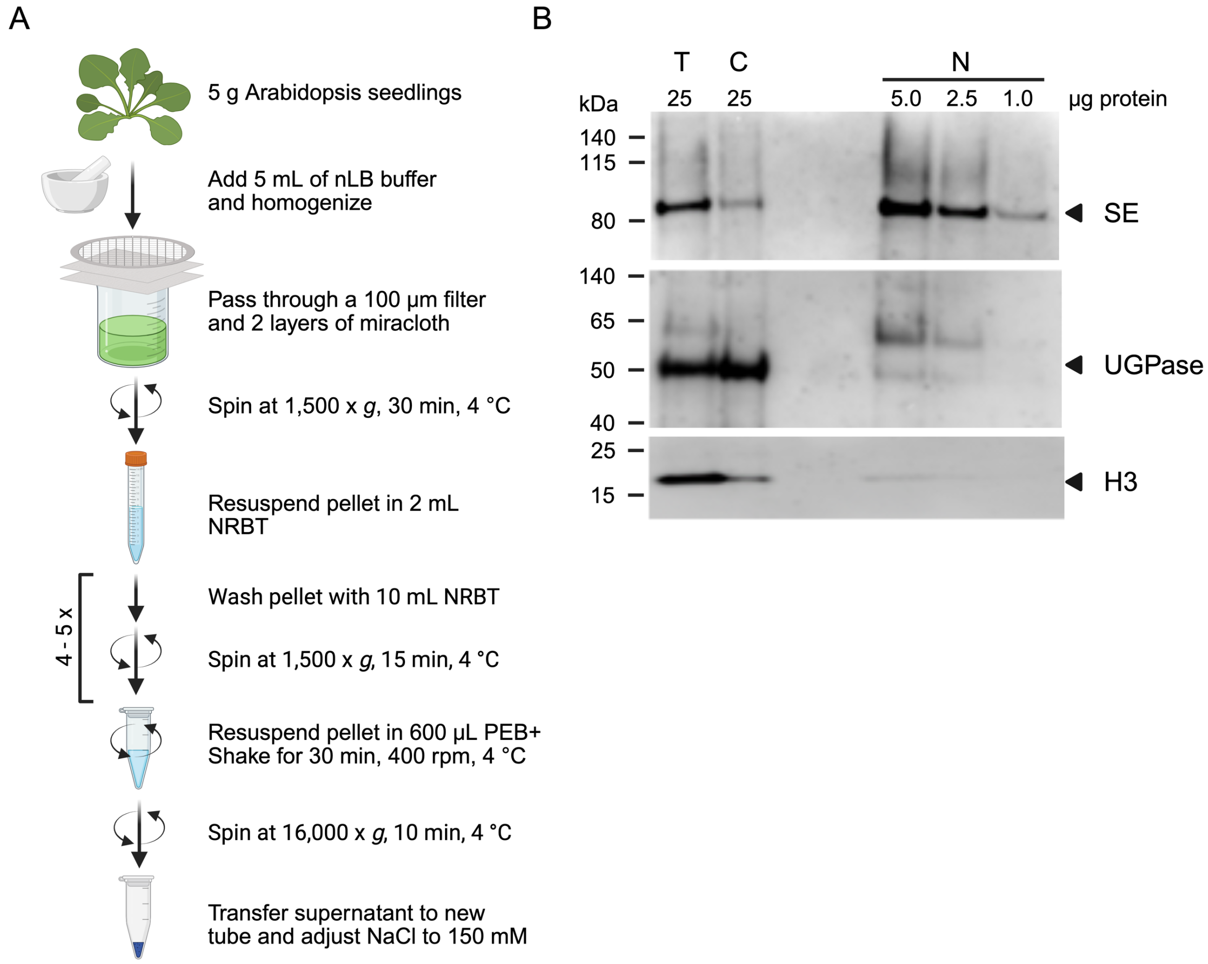


**Figure S2:** Characterization of nucleoplasmic extracts. (A) Scheme of the experimental procedure. (B) Immunoblot analysis of total lysate (T), cytoplascmic fraction (C), and nuclear extract (N). The indicated amount of the protein was separated by SDS-PAGE and the blots were probed with antibodies against nucleoplasmic SERRATE (top), cytoplasmic UGPase (middle) and chromatin-associated Histon H3 (bottom). The sizes of the molecular weight standards are indicated.

#### Figure S3

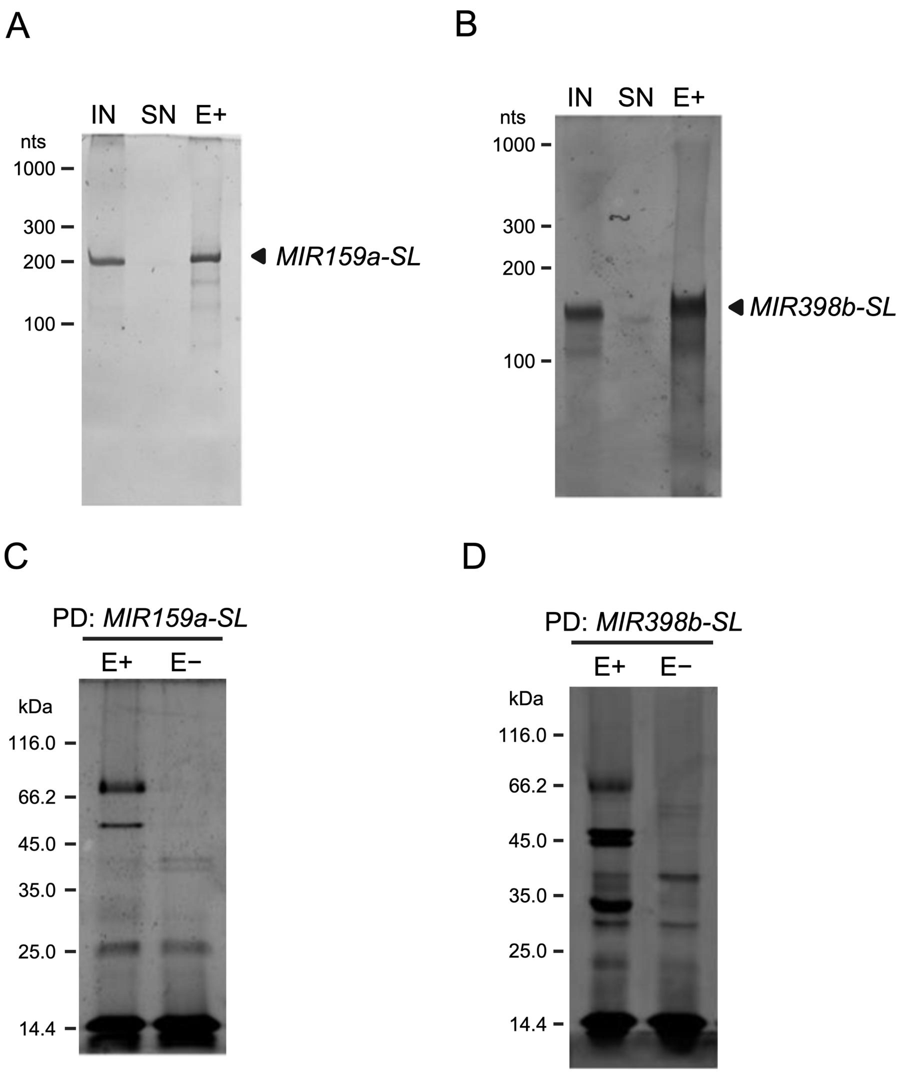


**Figure S3:** Control of the RNA and protein fractions of the RNA affinity pulldowns (PD). The integrity of the *in vitro* transcribed (A) *MIR159a*-SL, and (B) *MIR398b-SL* in the input (IN), supernatant (SN) and the eluate fraction (E+) was confirmed by urea polyacrylamide gel electrophoreses. Proteins recovered by pulldown with immobilized (C) *MIR159a-SL* and (D) *MIR398b-SL* were analysed by SDS gel electrophoreses and silver staining (E+). Proteins eluted from empty beads (E-) served as control.

#### Figure S4

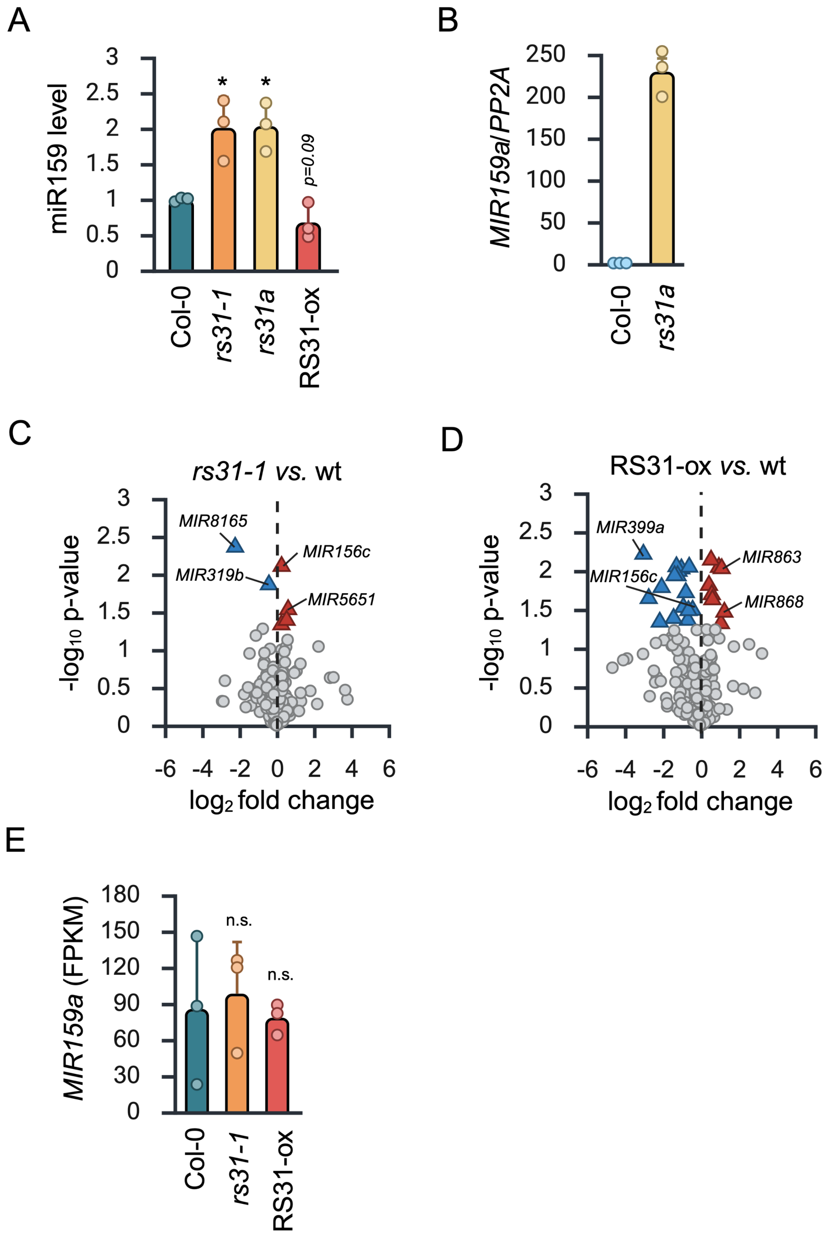


**Figure S4:** MiRNA levels in *rs31-1* mutant and RS31-ox lines. (A) Quantification of miR159 levels in Col-0 wild-type, *rs31-1*, *rs31a* and RS31-ox lines determined by small RNA gel blots (cf. Fig. 3B). MiR159 levels were quantified using ImageJ and levels are expressed relative to wild-type. (B) Level of *MIR159a* in *rs31a* determined by RT-qPCR. Shown are the mean ±SD of three biological replicates. A Student's *t*-test was performed to determine statistical significance (*** $p \leq 0.001$, ** $p \leq 0.01$, * $p \leq 0.05$, n.s., not significant). Transcriptome-wide identification of differentially expressed *MIRNAs* in either (C) *rs31-1* or (D) RS31-ox compared to wild-type [38]. (E) Level of *MIR159a* in *rs31-1* or RS31-ox relative to wild-type.

#### Figure S5

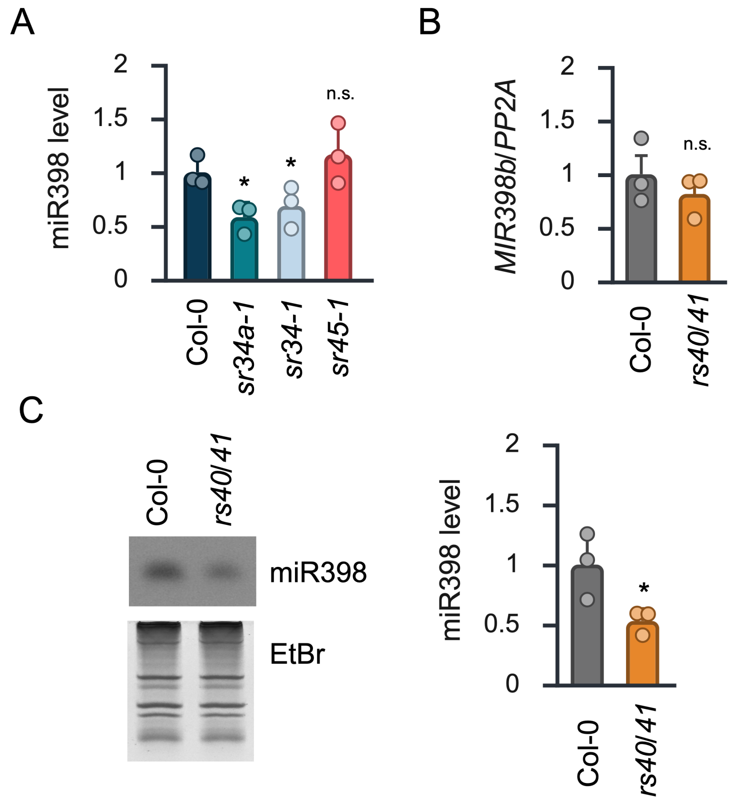


**Figure S5:** MiRNA levels in *sr34a-1, sr34-1, sr45-1* and *rs40/rs41* mutants. (A) Quantification of miR398 levels in Col-0 wild-type, *sr34a-1*, *sr34-1* and *sr45-1* mutant lines determined by small RNA gel blots (cf. Fig. 3D). MiR398 levels were quantified using ImageJ and levels are expressed relative to wild-type. (B) Level of *MIR398b* in *rs40/rs41* (*rs40/41*) determined by RT-qPCR. (C) A small RNA gel blot of the *rs40/rs41* (*rs40/41*) mutant and the corresponding wild-type plants was hybridized with an anti-miR398 probe (left). Ethidium bromide (EtBr) staining served as a loading control. MiR398 signals were quantified using ImageJ (right). Shown are the mean ±SD of three biological replicates. A Student's *t*-test was performed to determine statistical significance (*** $p \leq 0.001$, ** $p \leq 0.01$, * $p \leq 0.05$, n.s., not significant).

#### Figure S6

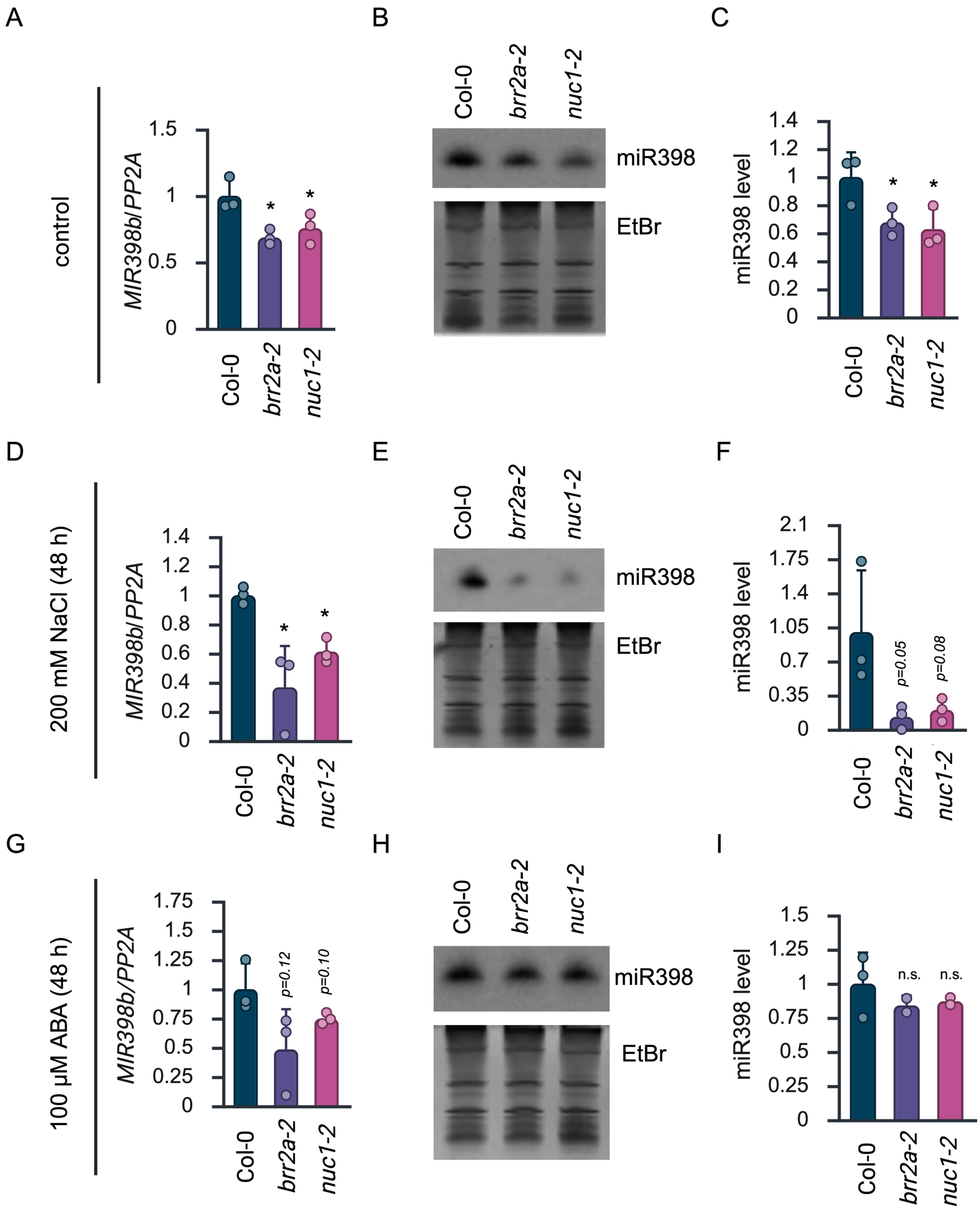


**Figure S6:** MiRNA levels in *brr2a-2* and *nuc1-2* under non-stress and abiotic stress conditions. The levels of *MIR398b* in A, D, and G and mature miR398 in B, E, and H were determined by RT-qPCR and small RNA gel blots, respectively, under (A-C) untreated (non-stress control) conditions or upon treatment with (D-F) 200 mM sodium chloride (NaCl) or (G-I) 100 µM abscisic acid (ABA) for 48 hours (h). Small RNA blots were hybridized with an anti-miR398 probe and ethidium bromide (EtBr) staining served as a loading control. MiR398 signals in C, F, and I were quantified using ImageJ. Shown are the mean ±SD of three biological replicates. A Student's *t*-test was performed to determine statistical significance (*** $p \leq 0.001$, ** $p \leq 0.01$, * $p \leq 0.05$, n.s., not significant).

### Figure S7

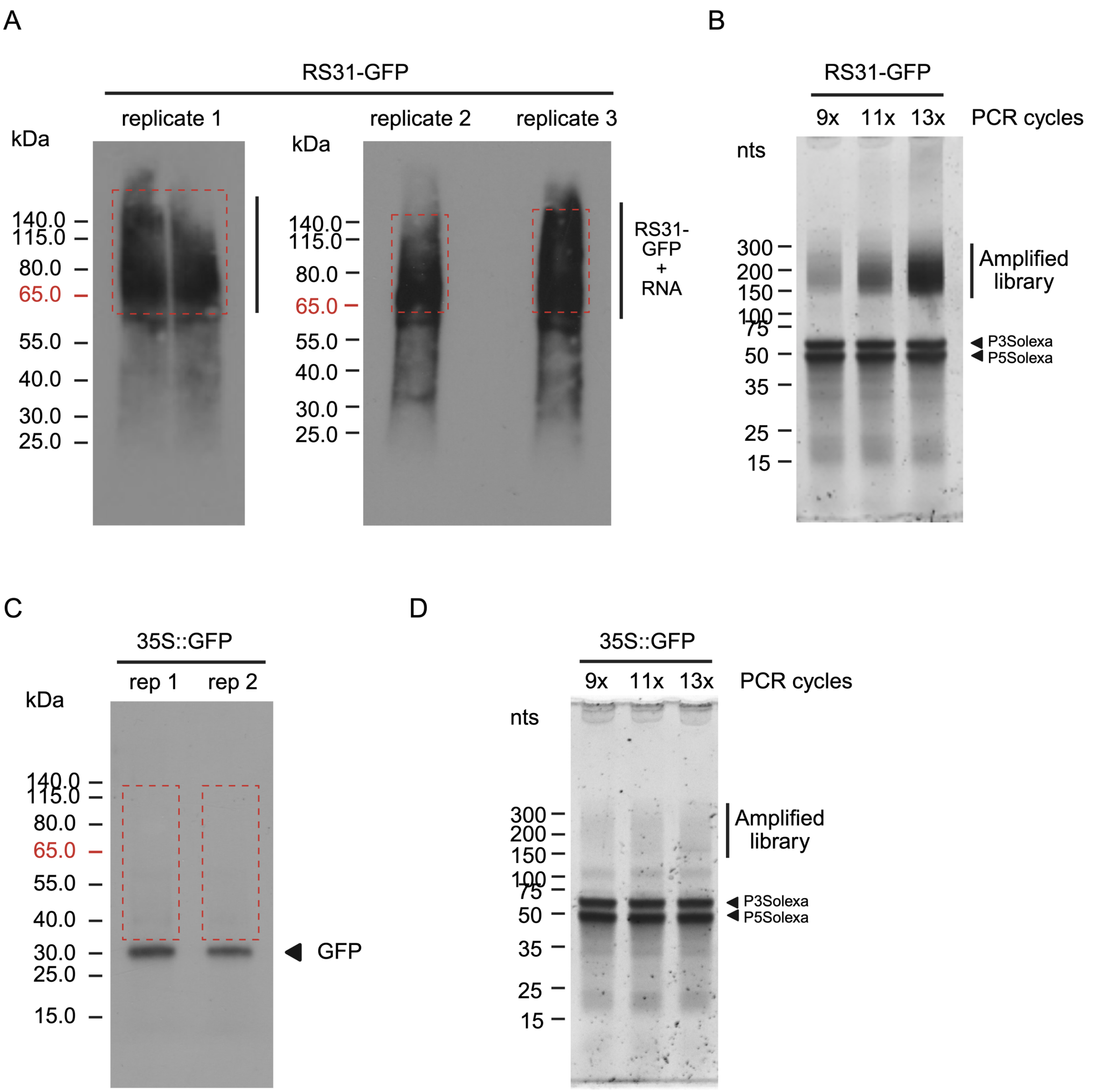


**Figure S7:** Plant iCLIP2 of RS31-GFP. (A) Autoradiograms of RNA-protein complexes from *RS31::RS31-GFP rs31-1* after UV crosslinking and immunoprecipitation in three replicates. For technical reasons, the sample of the first replicate was split into two lanes after elution in LDS sample buffer. The regions containing the crosslinked RNAs used for library generation are indicated (red dashed boxes). A control treatment with high RNase I was previously described [38]. (B) Gel electrophoresis of amplified plant iCLIP2 cDNA libraries for RS31-GFP with increasing PCR cycles. P3Solexa and P5Solexa denote primers used for library amplification. (C) Autoradiogram of RNA-protein complexes from *35S::GFP* control plants after UV crosslinking and immunoprecipitation in two replicates. (D) Gel electrophoresis of amplified *35S::GFP* control libraries with increasing PCR cycles.

#### Figure S8

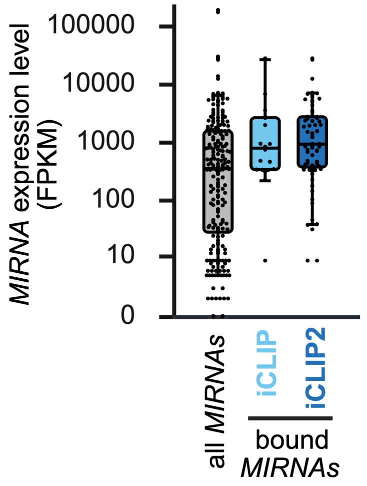


**Figure S8:** Shown is the average expression level (FPKM) of all *MIRNAs* expressed in Col-0 wild-type (FPKM ≥ 0) compared to the average expression level of bound *MIRNAs* identified for RS31-GFP by iCLIP or plant iCLIP2.

#### Figure S9

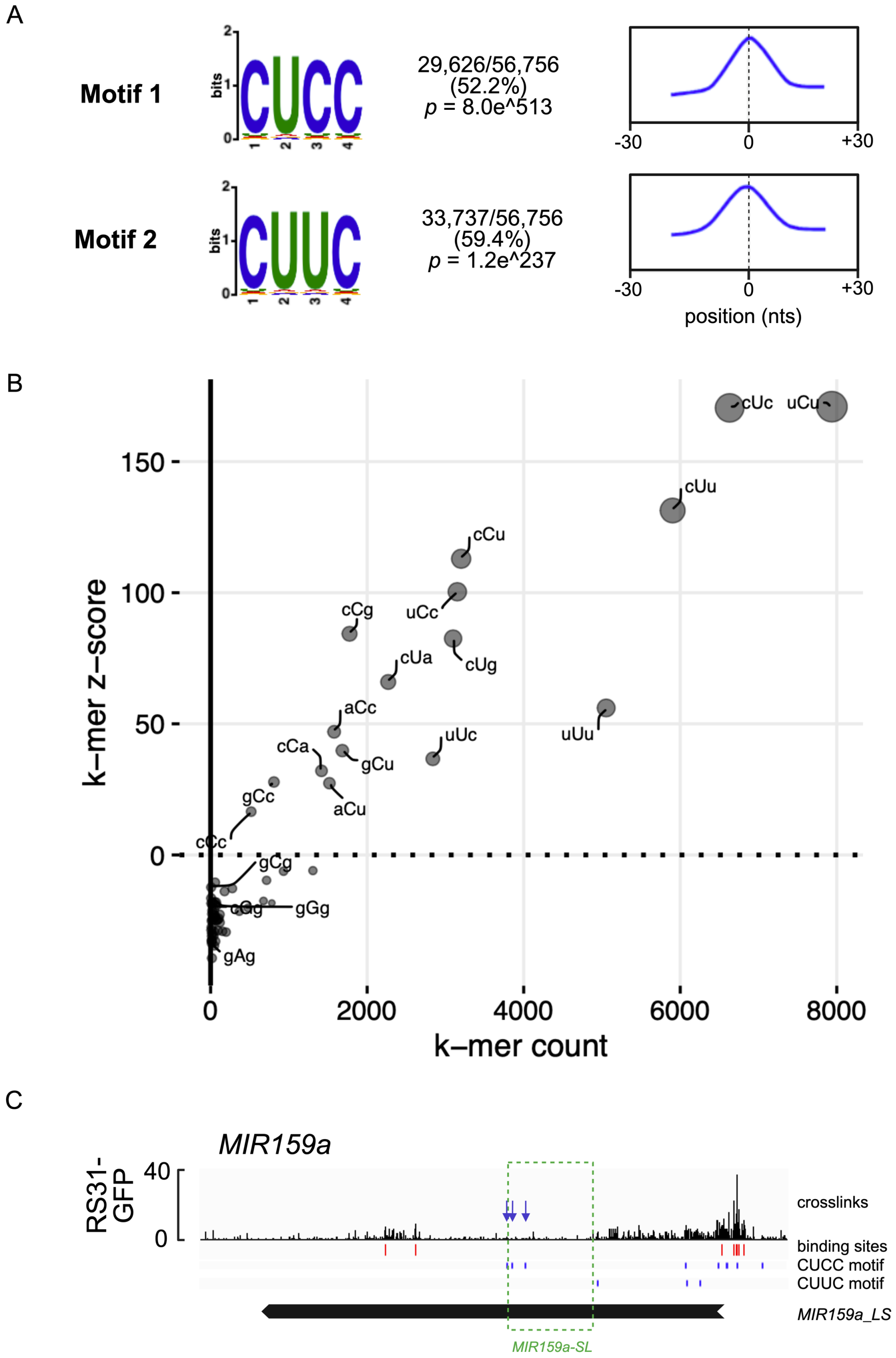


**Figure S9:** Sequence motifs enriched at the RS31-GFP binding sites. (A) Significantly enriched motifs determined by STREME (left). Density distribution of the identified binding motif around the binding site (right). The centre position (zero) corresponds to the binding site peak. (B) Scatterplot displaying trimer counts (x-axis) and trimer z-scores (y-axis). Trimers in the upper right corner are enriched in comparison to a random background correction model. Letters in upper case mark the peak position of the binding site. (C) Crosslinks and significant binding sites (red boxes) of RS31-GFP on *MIR159a* as determined by plant iCLIP2. The blue boxes mark the position of the identified CU-rich binding motifs CUCC and CUUC in the region of the annotated *MIR159a_LS* (black arrow). The green dashed box marks the region of the annotated *MIR159a* stem-loop (*SL*). Blue arrows highlight the CU-rich binding motifs within the *MIR159a-SL*.

#### Figure S10

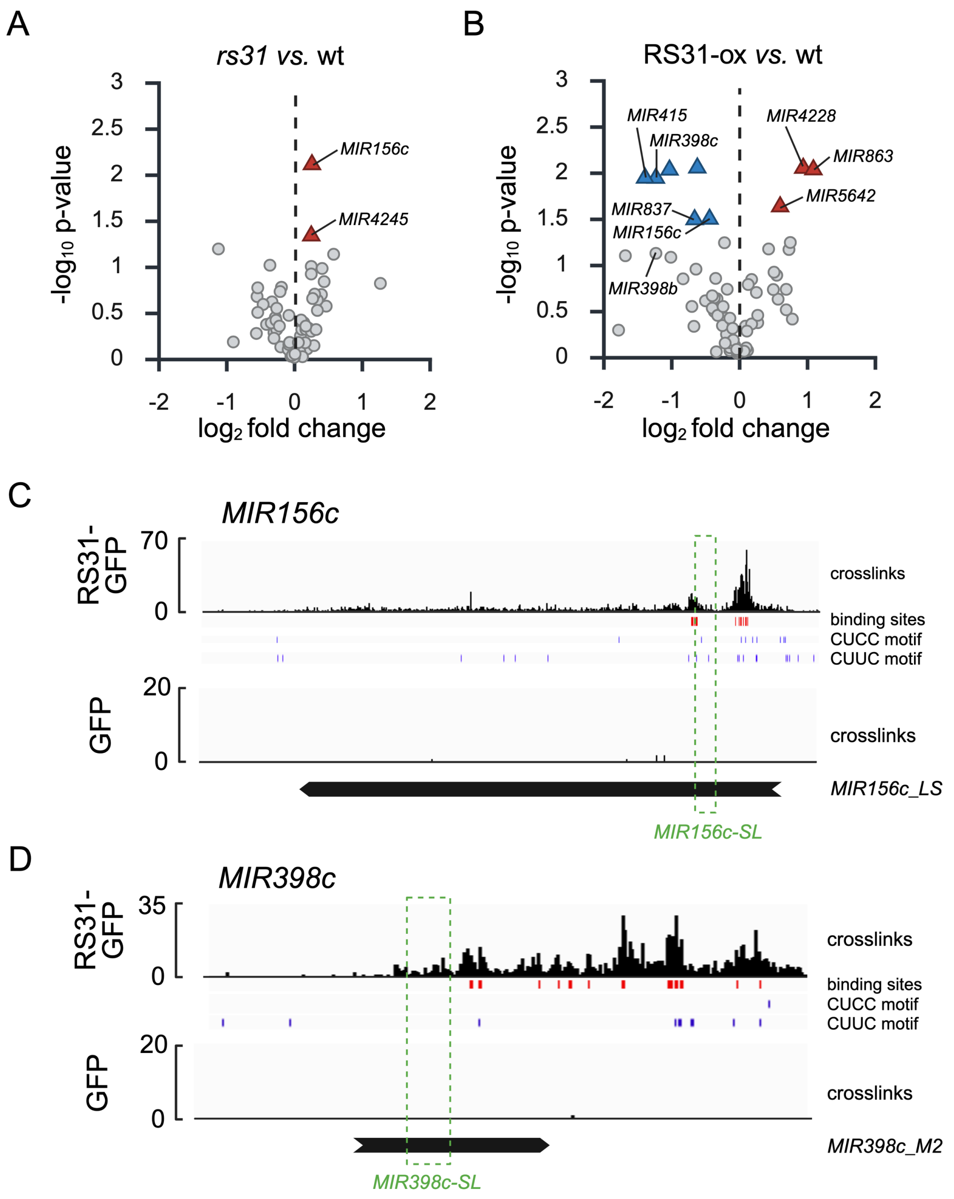


**Figure S10:** Transcriptome-wide identification of RS31-GFP bound and differentially expressed *MIRNAs* in either (A) *rs31-1* or (B) RS31-ox compared to wild-type. Crosslinks and significant binding sites of RS31-GFP on (C) *MIR156c* and (D) *MIR398c* as determined by plant iCLIP2.Blue boxes mark the position of the identified CU-rich binding motifs CUCC and CUUC in the region of the annotated *MIRNAs* (black arrow). The green dashed box marks the region of the annotated *MIRNA* stem-loops (*SL*).

### Figure S11

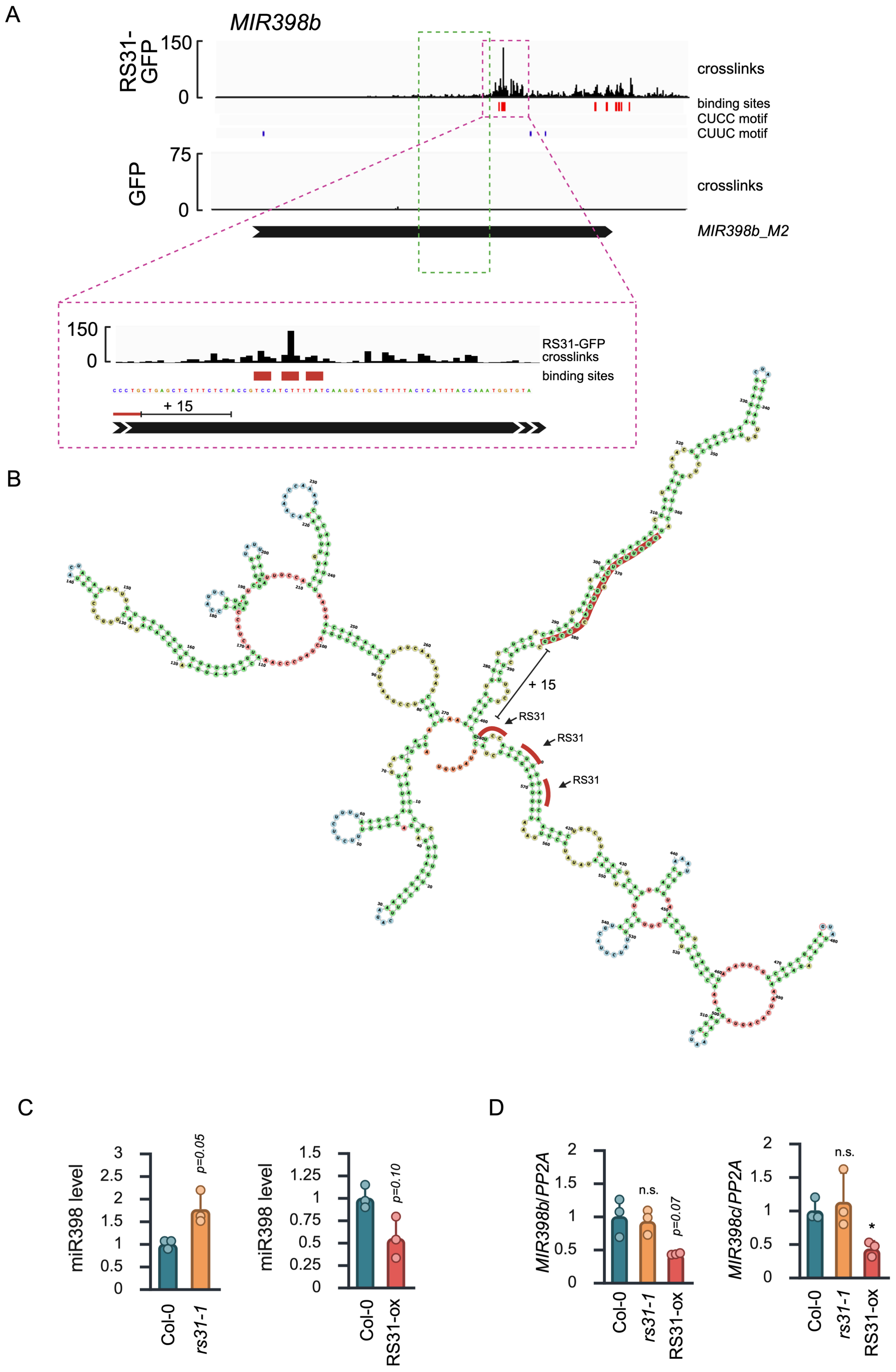
A
MIR398b
RS31-GFP
150
0
crosslinks
binding sites
CUCC motif
CUUC motif
GFP
75
0
crosslinks
MIR398b_M2
150
0
RS31-GFP crosslinks
binding sites
+ 15
B
+ 15
RS31
RS31
RS31
C
miR398 level
Col-0
rs31-1
p=0.05
miR398 level
Col-0
RS31-ox
p=0.10
D
MIR398b/PP2A
Col-0
rs31-1
RS31-ox
n.s.
p=0.07
MIR398c/PP2A
Col-0
rs31-1
RS31-ox
n.s.
*

**Figure S11:** (A) Crosslinks and significant binding sites (red boxes) of RS31-GFP on *MIR398b* as determined by plant iCLIP2. Blue boxes mark the position of the identified CU-rich binding motifs CUCC and CUUC in the region of the annotated *MIR398b_M2* (black arrow). The green dashed box marks the region of the annotated *MIR398b* stem-loop (*SL)*. The purple dashed box focusses on the RS31-GFP binding sites downstream of the base of the *MIR398b-SL*. (B) *In silico* predicted secondary structure of *MIR398b*. Red bars indicate the binding sites near the base of the stem-loop highlighted in (A). +15 denotes a region important for correct processing of *MIR398b*. (C) Level of *MIR398b* and *MIR398c* in *rs31-1* and RS31-ox compared to wild-type determined by RT-qPCR. (D) Levels of mature miR398 in *rs31-1* and RS31-ox compared to wild-type. After small RNA blots, levels were quantified using ImageJ and levels are expressed relative to wild-type. Shown are the mean ±SD of three biological replicates. A Student's *t*-test was performed to determine statistical significance (*** $p \leq 0.001$, ** $p \leq 0.01$, * $p \leq 0.05$, n.s., not significant).

#### Figure S12

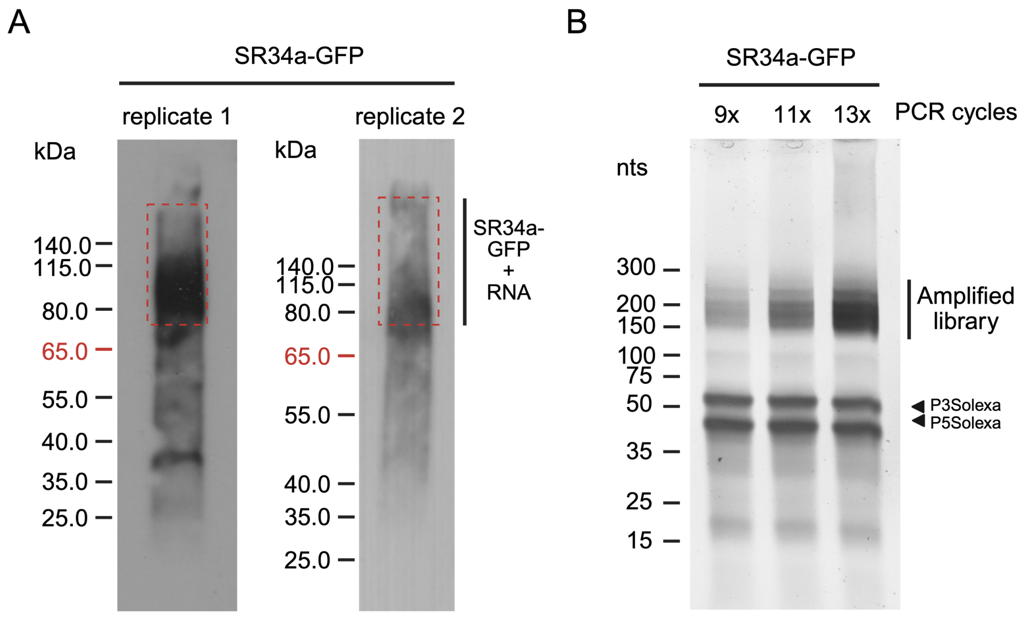


**Figure S12:** Plant iCLIP2 of SR34a-GFP. (A) Autoradiograms of RNA-protein complexes from *SR34a::SR34a-GFP sr34a-1* after UV crosslinking and immunoprecipitation in two replicates. The regions containing the crosslinked RNAs used for library generation are indicated (red dashed boxes). (B) Gel electrophoresis of amplified plant iCLIP2 cDNA libraries for SR34a-GFP with increasing PCR cycles. P3Solexa and P5Solexa denote primers used for library amplification.

### Figure S13

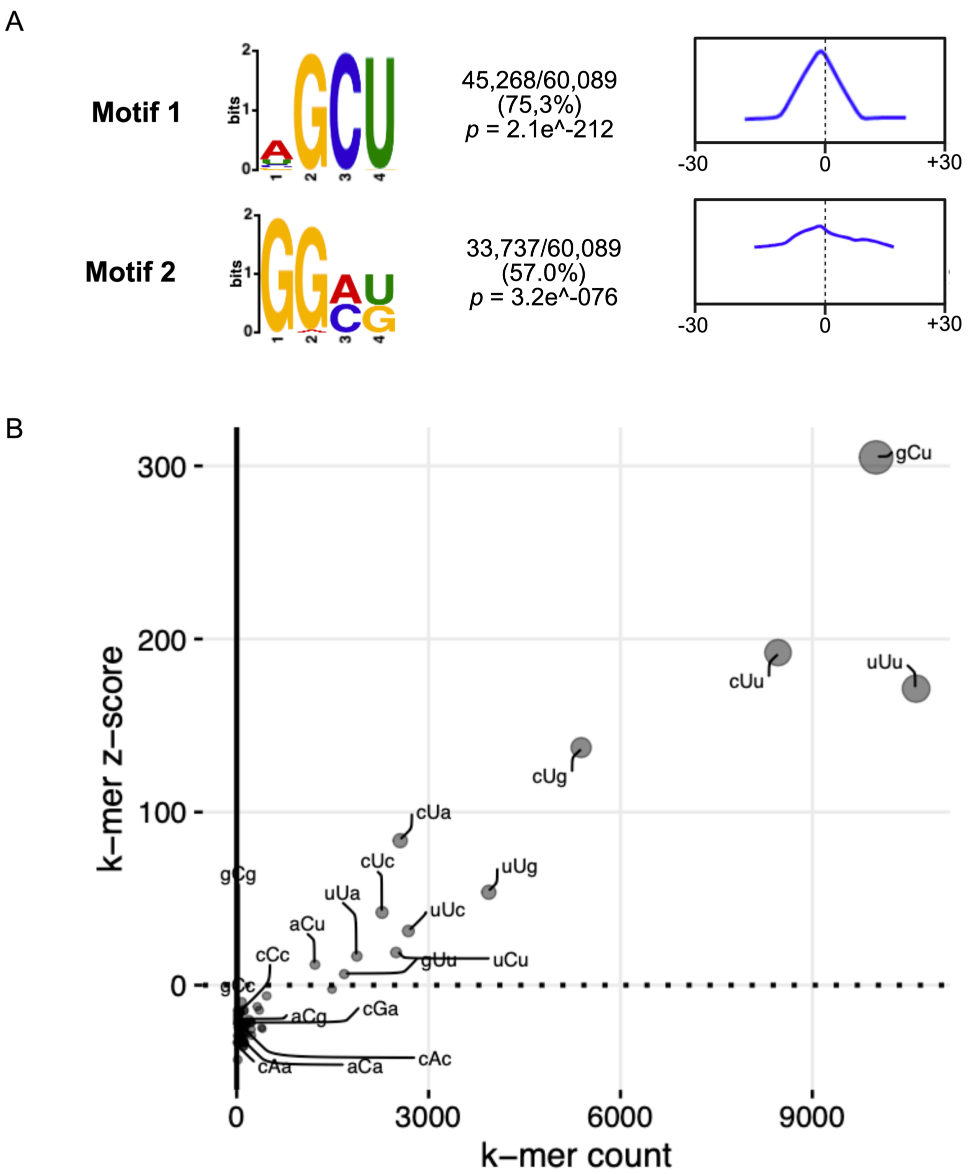


**Figure S13:** Sequence motifs enriched at SR34a-GFP binding sites. (A) Significantly enriched motifs determined by STREME (left). Density distribution of the identified binding motif around the binding site (right). The centre position (zero) corresponds to the binding site peak. (B) Scatterplot displaying trimer counts (x-axis) and trimer z-scores (y-axis). Trimers in the upper right corner are enriched in comparison to a random background correction model. Letters in upper case mark the peak position of the binding site.

#### Figure S14

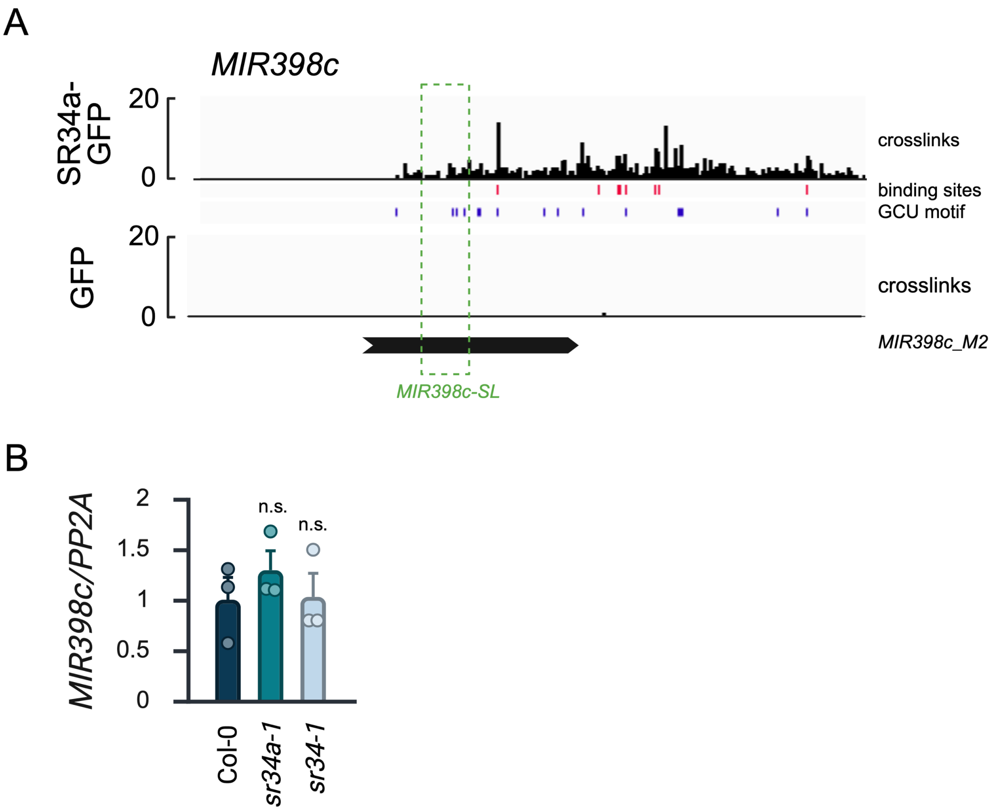


**Figure S14:** (A) Crosslinks and significant binding sites of SR34a-GFP on *MIR398c* as determined by plant iCLIP2. Blue boxes mark the position of the identified CCU binding motif in the region of the annotated *MIR398c_M2* (black arrow). The track displaying crosslinks of *35S::GFP* plants are given as a control. The green dashed box marks the region of the annotated *MIR398c* stem-loop (*SL)*. (B) RT-qPCR analysis of miR398 in Col-0 wild-type, *sr34a-1,* and *sr34-1*. Shown are the mean ±SD of three biological replicates. A Student's *t*-test was performed to determine statistical significance (*** $p \leq 0.001$, ** $p \leq 0.01$, * $p \leq 0.05$, n.s., not significant).

#### Figure S15

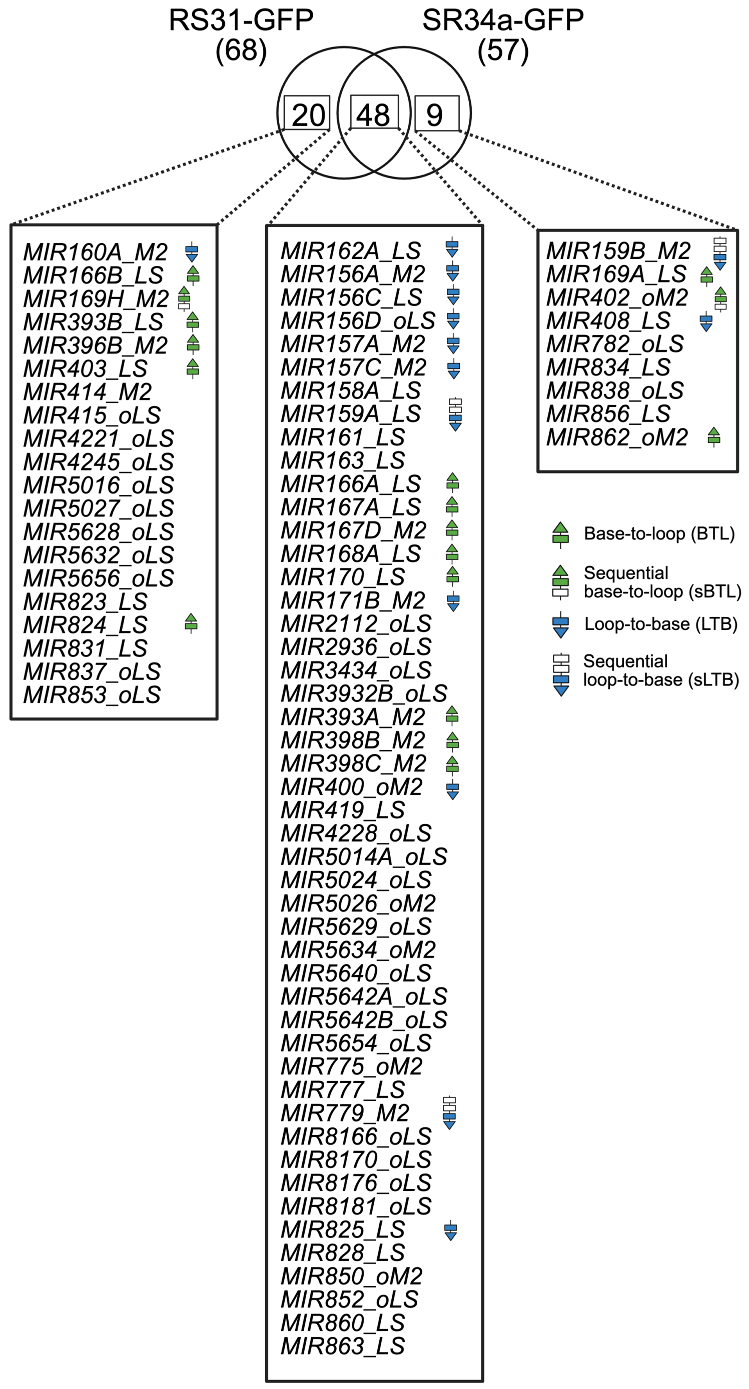


**Figure S15:** Comparison of the *MIRNAs* directly bound by RS31-GFP and/or SR34a-GFP *in vivo*. Evolutionarily conserved *MIRNAs* for which a processing mechanism has been described are labelled accordingly (see figure legend).

#### Figure S16

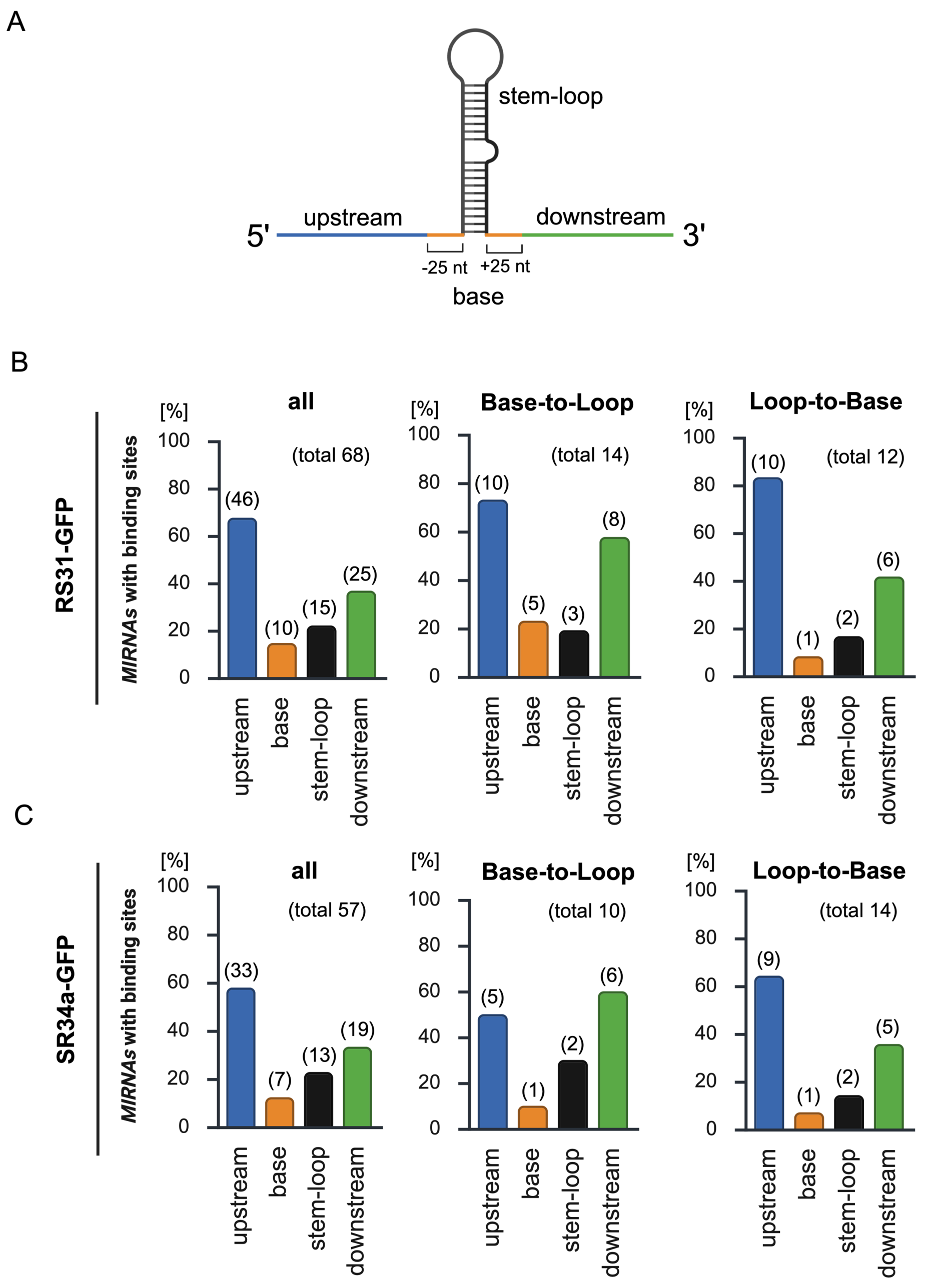


**Figure S16:** Distribution of binding sites across RS31-GFP and SR34a-GFP *MIRNA* targets. (A) Annotated *MIRNAs* were divided into defined regions upstream (blue) and downstream (green) of the stem-loop, the stem-loop itself (black) and a base region (orange). (B) Distribution of binding sites on all *MIRNAs* bound RS31-GFP (left) or those, which were either known to be processed in base-to-loop (middle) or loop-to-base direction (right). (C) Distribution of SR34a-GFP *MIRNA* binding sites as described for RS31-GFP. Absolute numbers of *MIRNAs* are given in brackets.

#### Figure S17

A

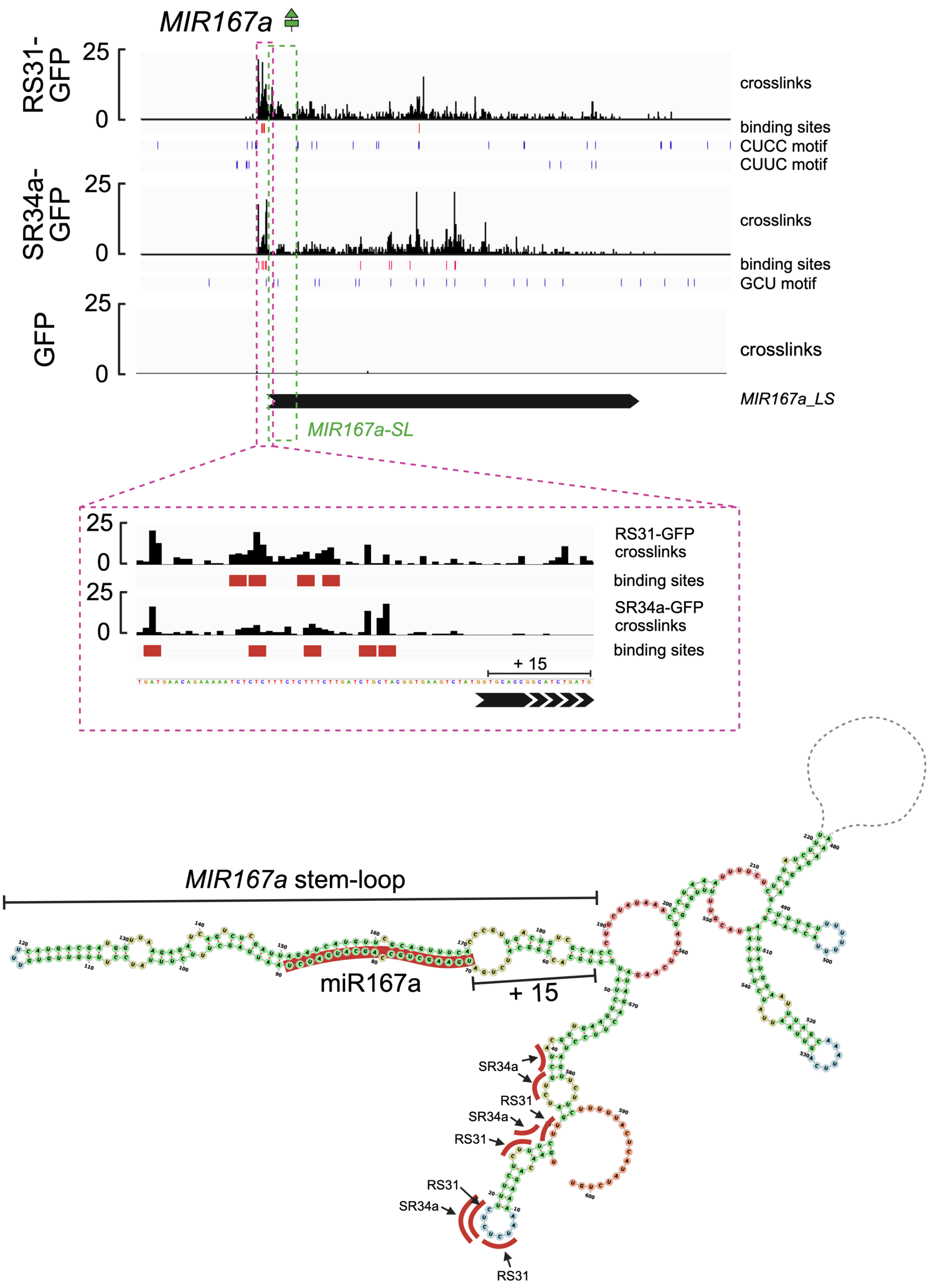

MIR167a
RS31-GFP
25
0
crosslinks
binding sites
CUCC motif
CUUC motif
SR34a-GFP
25
0
crosslinks
binding sites
GCU motif
GFP
25
0
crosslinks
MIR167a_LS
MIR167a-SL
RS31-GFP crosslinks
binding sites
SR34a-GFP crosslinks
binding sites
+ 15
MIR167a stem-loop
miR167a
+ 15
SR34a
RS31
SR34a
RS31
RS31
SR34a
RS31

B

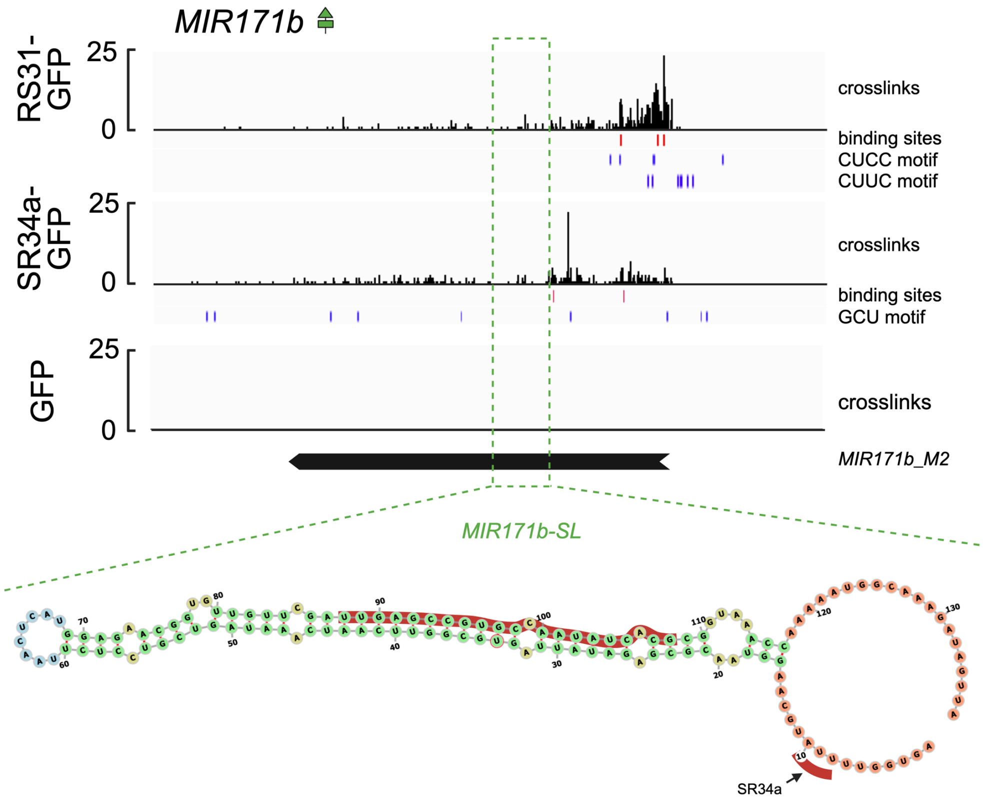


C

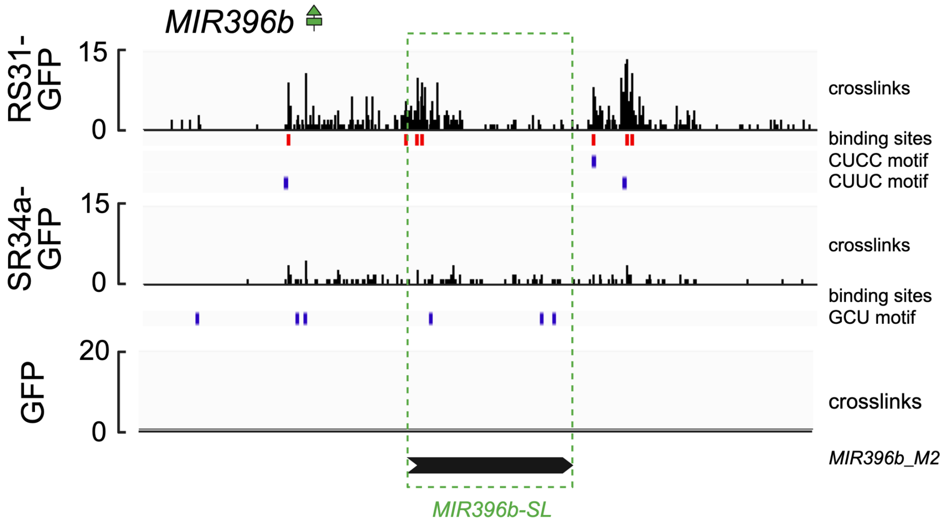


**Figure S17: :** Crosslinks and significant binding sites (red boxes) of RS31-GFP and SR34a-GFP on *MIRNAs* as determined by plant iCLIP2. Blue boxes mark the position of the identified binding motifs for RS31-GFP and SR34a-GFP. The green dashed box marks the region of the annotated *MIRNA* stem-loops (*SL)*. (A) The purple dashed box focusses on the RS31-GFP binding sites downstream of the base of the *MIR167a-SL*. The *in silico* predicted secondary structure of *MIR167a* is shown at the bottom. Red bars indicate RS31-GFP and SR34a-GFP binding sites near the base of the stem-loop highlighted above. +15 denotes a region important for correct processing of *MIR398b*. RS31-GFP and SR34a-GFP binding sites on the annotated (B) *MIR171b*, and (C) *MIR396b.*
