## Additional file 4 for "Identification and binding-site mapping of RNA-binding proteins interacting with microRNA precursors in Arabidopsis"

**Figure S18:** Uncropped blots to Figure S2B.

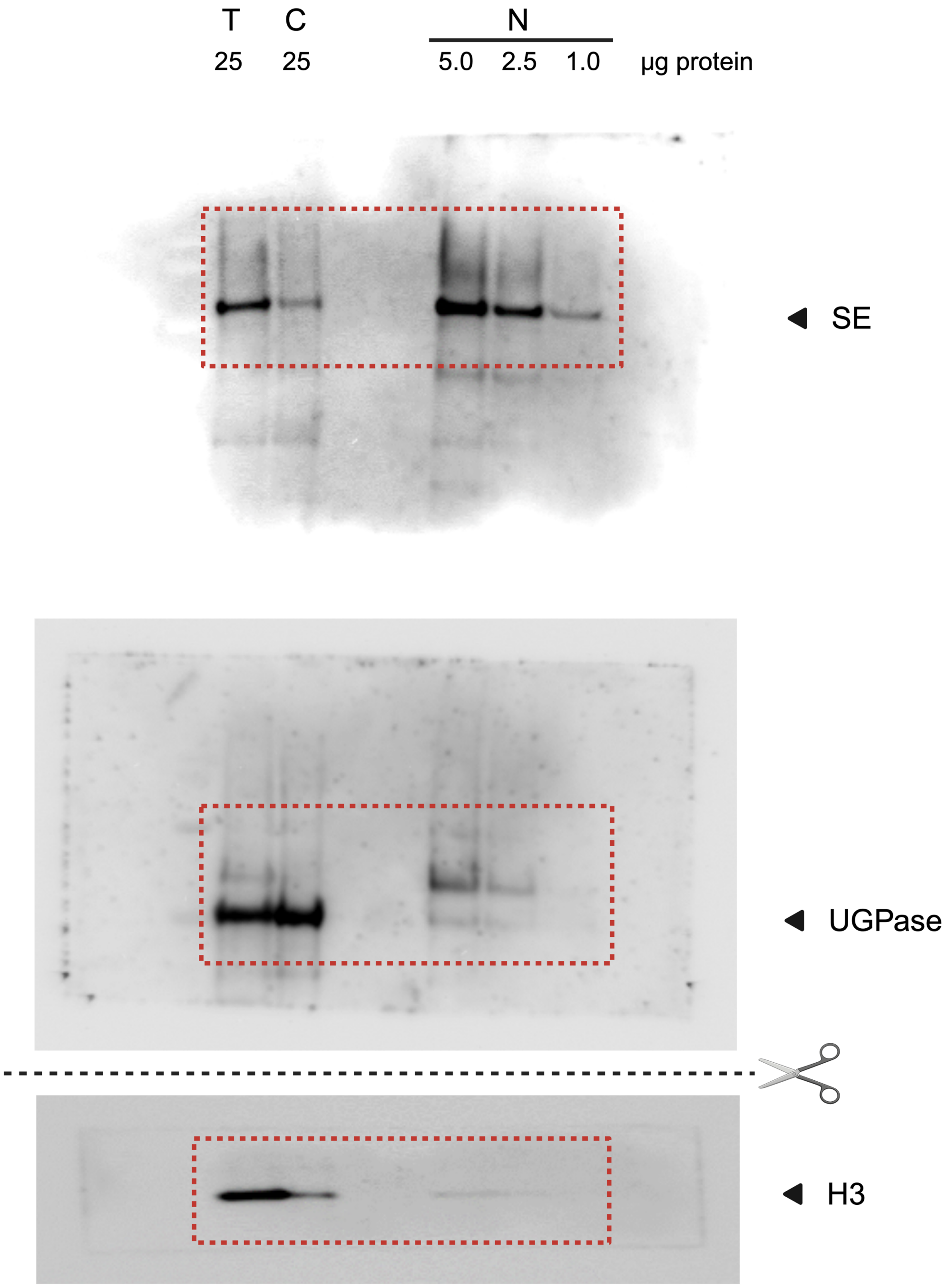


The red dotted box marks the cropped area shown in the main figure.

**Figure S19:** Uncropped gelst o Figure S3A-D.

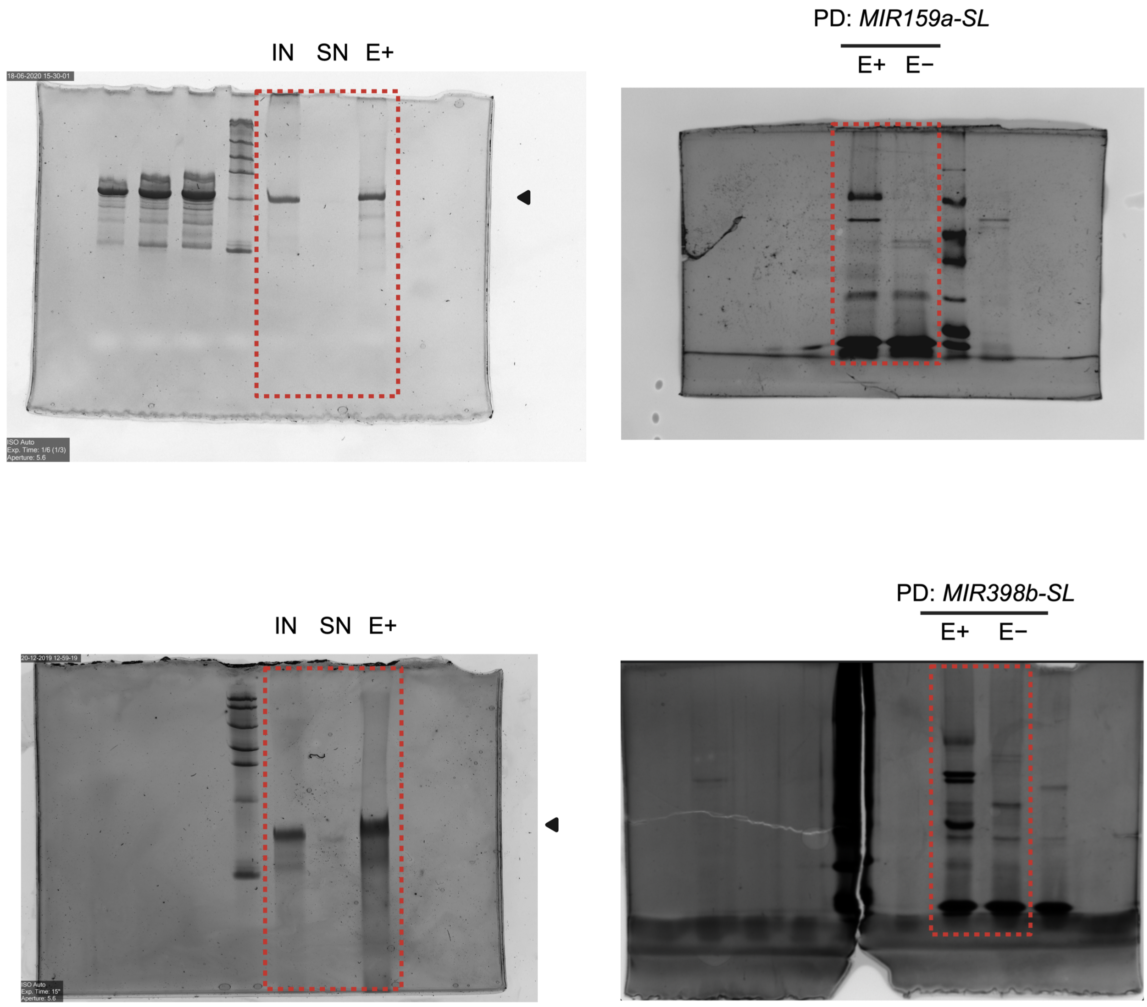


The red dotted box marks the cropped area shown in the main figure.

**Figure S20:** Uncropped gels and blots to Figure 3B and S4A.

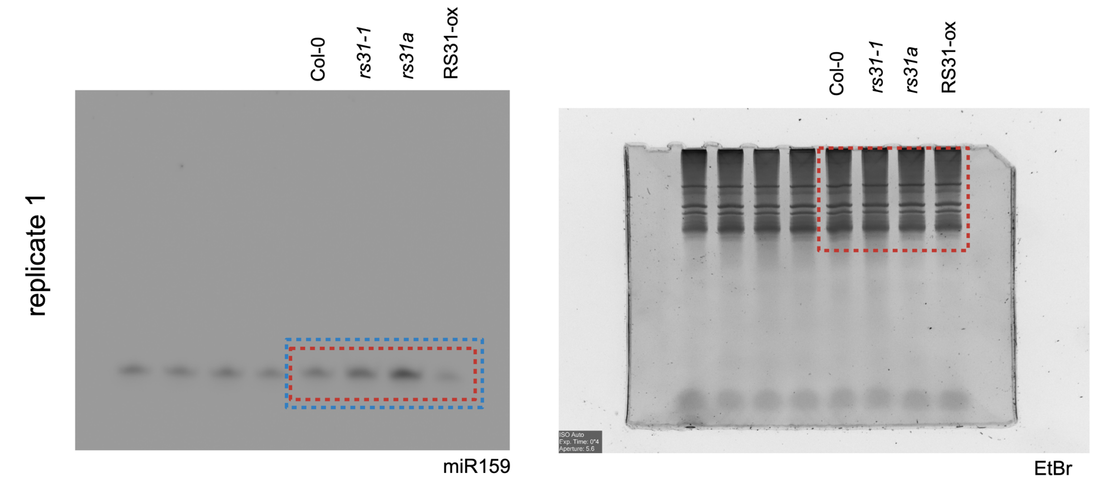


Uncropped gels and blots to Figure S4A.

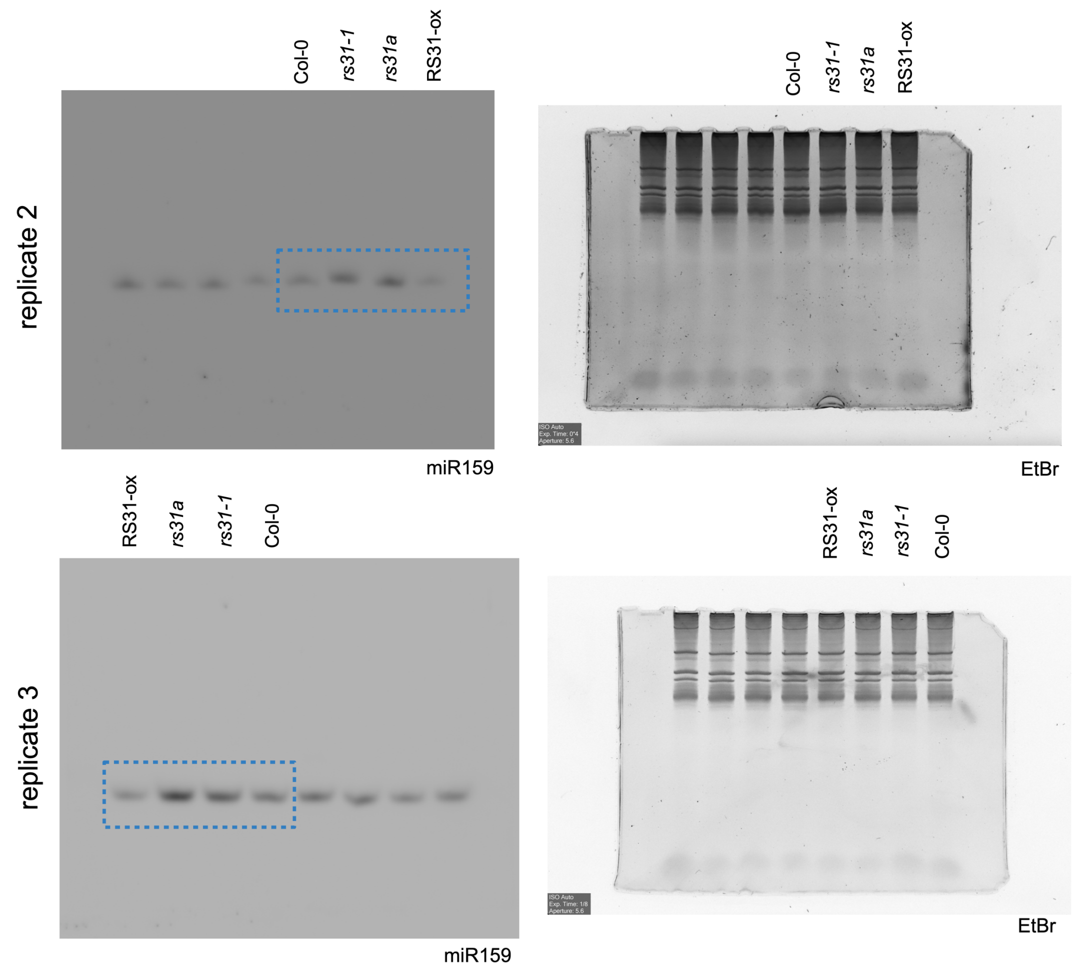


The red dotted box marks the cropped area shown in the main figure. The blue dotted box indicates the area and bands used for quantification with ImageJ.

**Figure S21:** Uncropped gels and blots to Figure 3D and S5A/C.

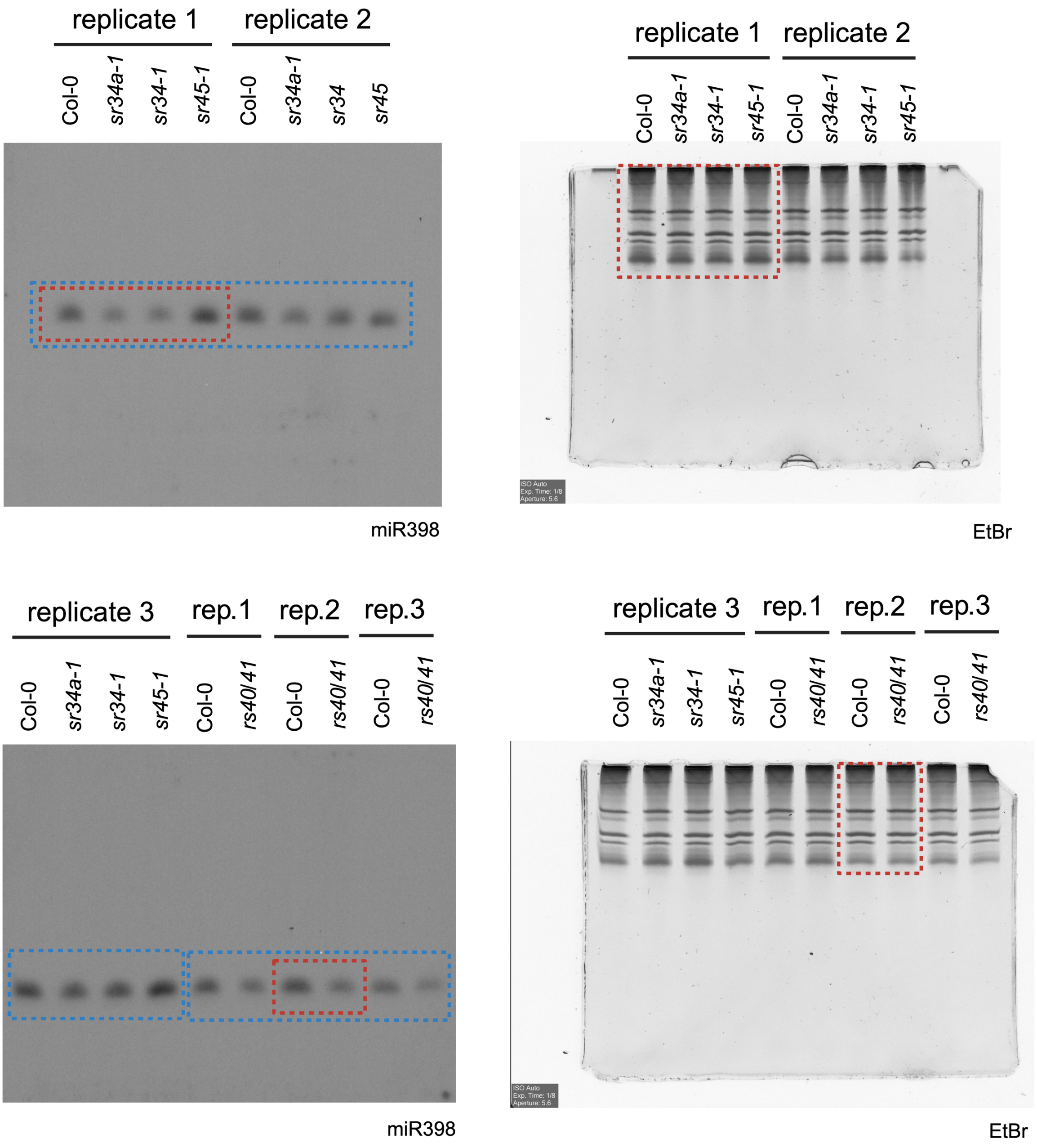


The red dotted box marks the cropped area shown in the main figure. The blue dotted box indicates the area and bands used for quantification with ImageJ.

**Figure S22:** Uncropped blots and gels to Figure S6A-I.

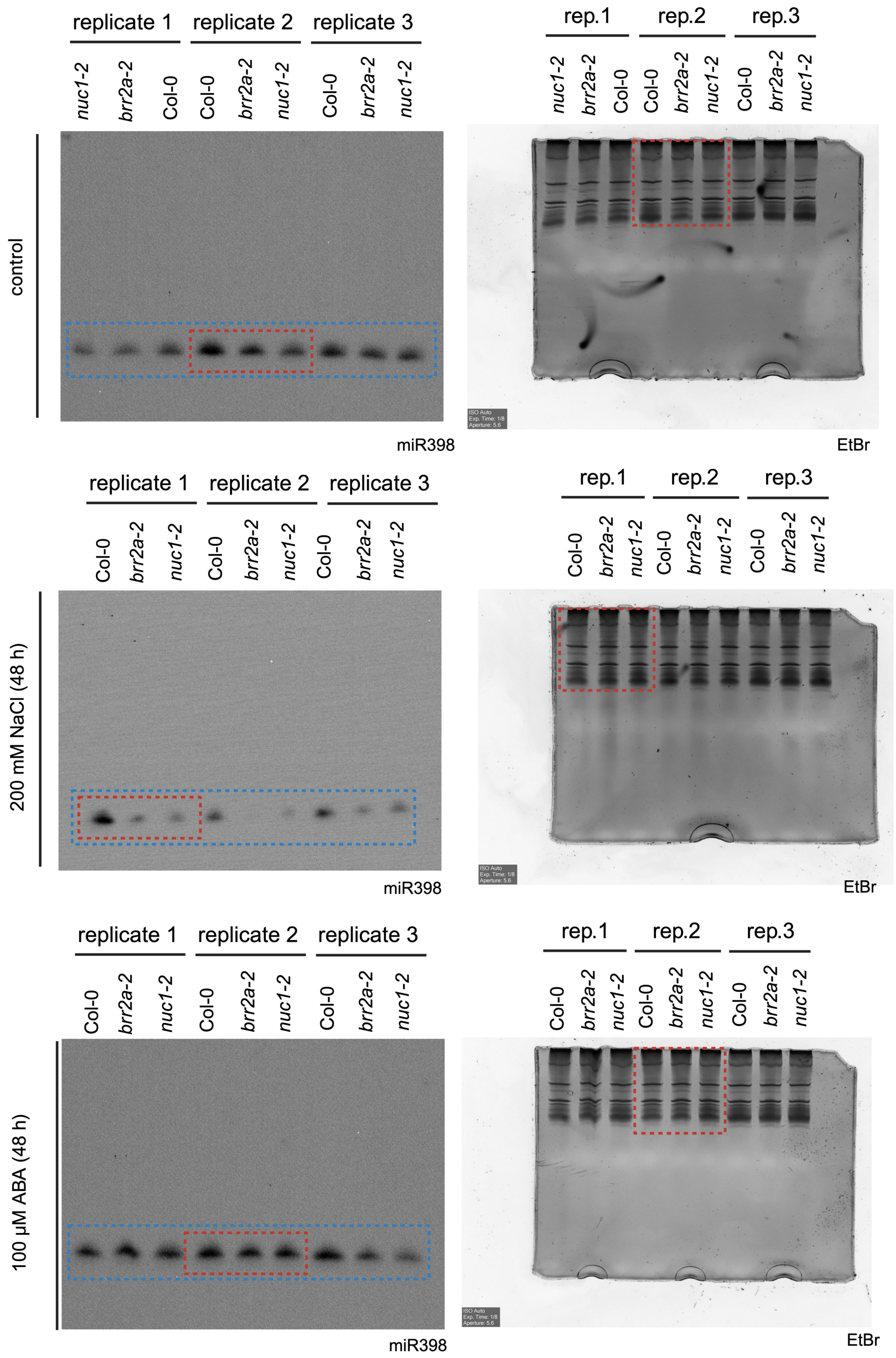


The red dotted box marks the cropped area shown in the main figure. The blue dotted box indicates the area and bands used for quantification with ImageJ.

**Figure S23:** Uncropped autoradiograms and gels to Figure S7A/B.

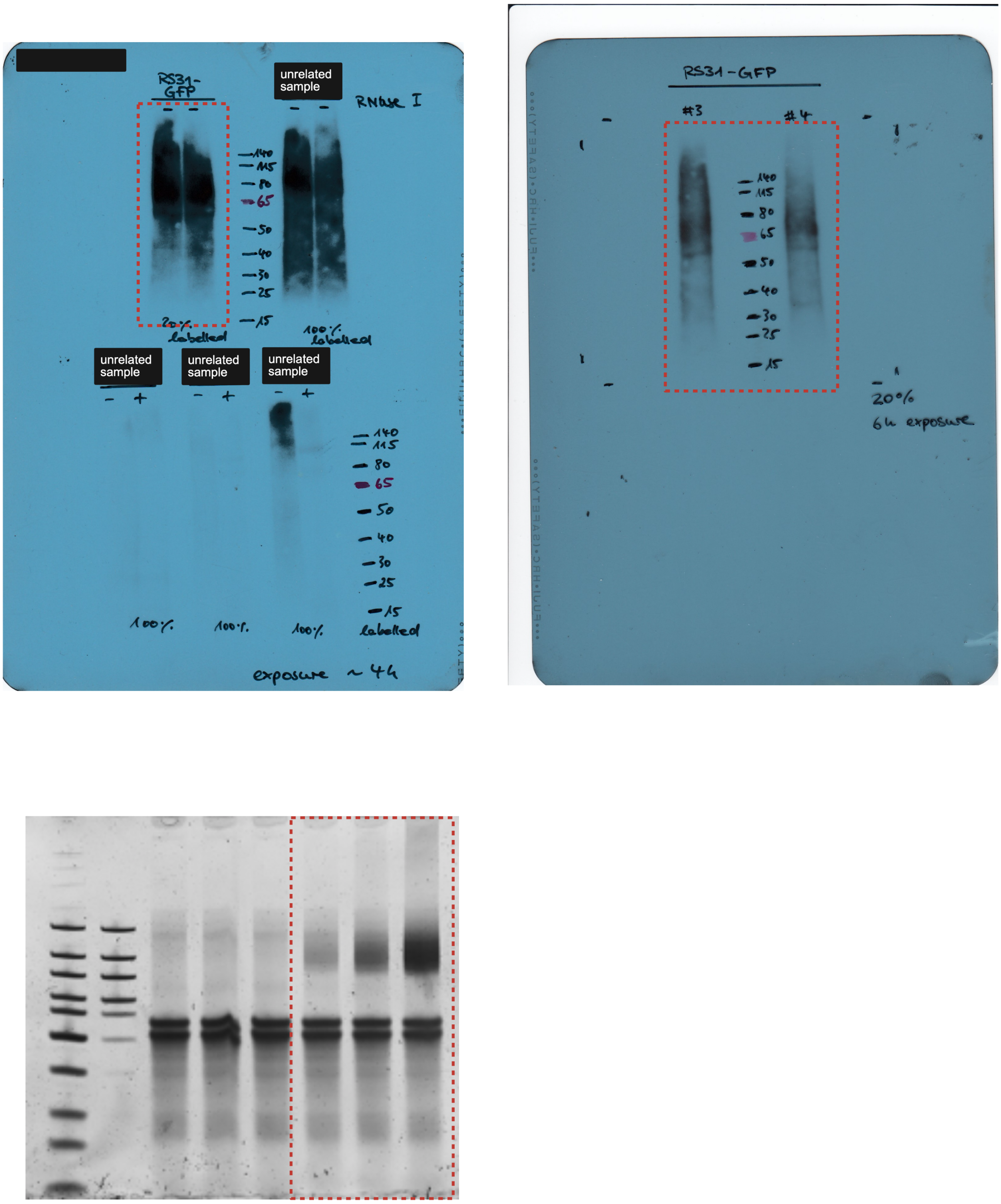


The red dotted box marks the cropped area shown in the main figure.

**Figure S24:** Uncropped autoradiograms and gels to Figure S7C/D.

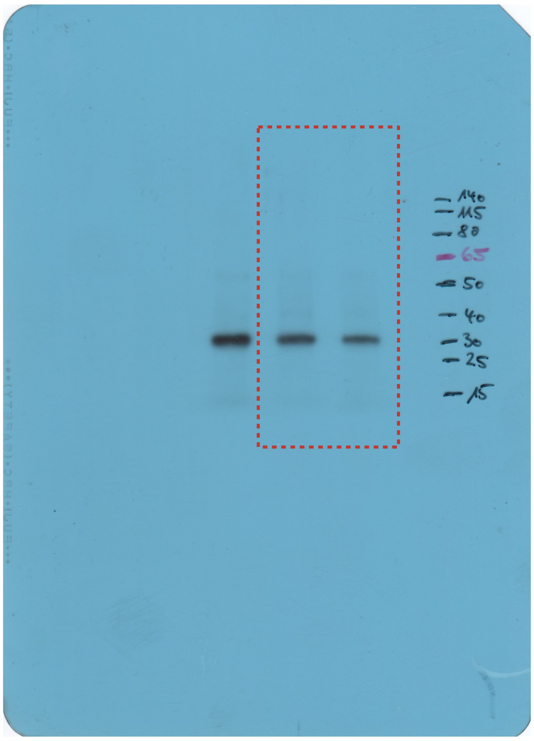


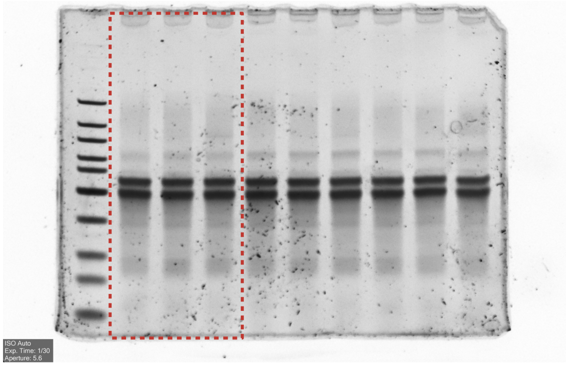


The red dotted box marks the cropped area shown in the main figure.

**Figure S25:** Uncropped autoradiogram and gel to Figure 5A/B.

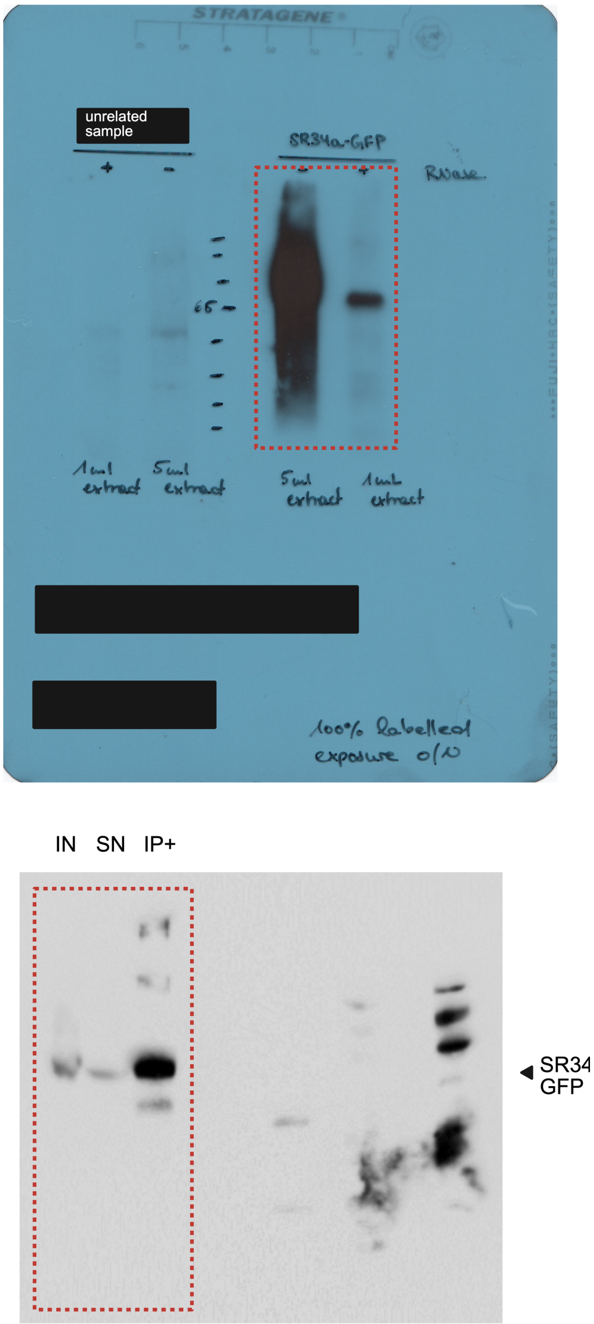


The red dotted box marks the cropped area shown in the main figure.

**Figure S26:** Uncropped blots and gels to Figure S11C.

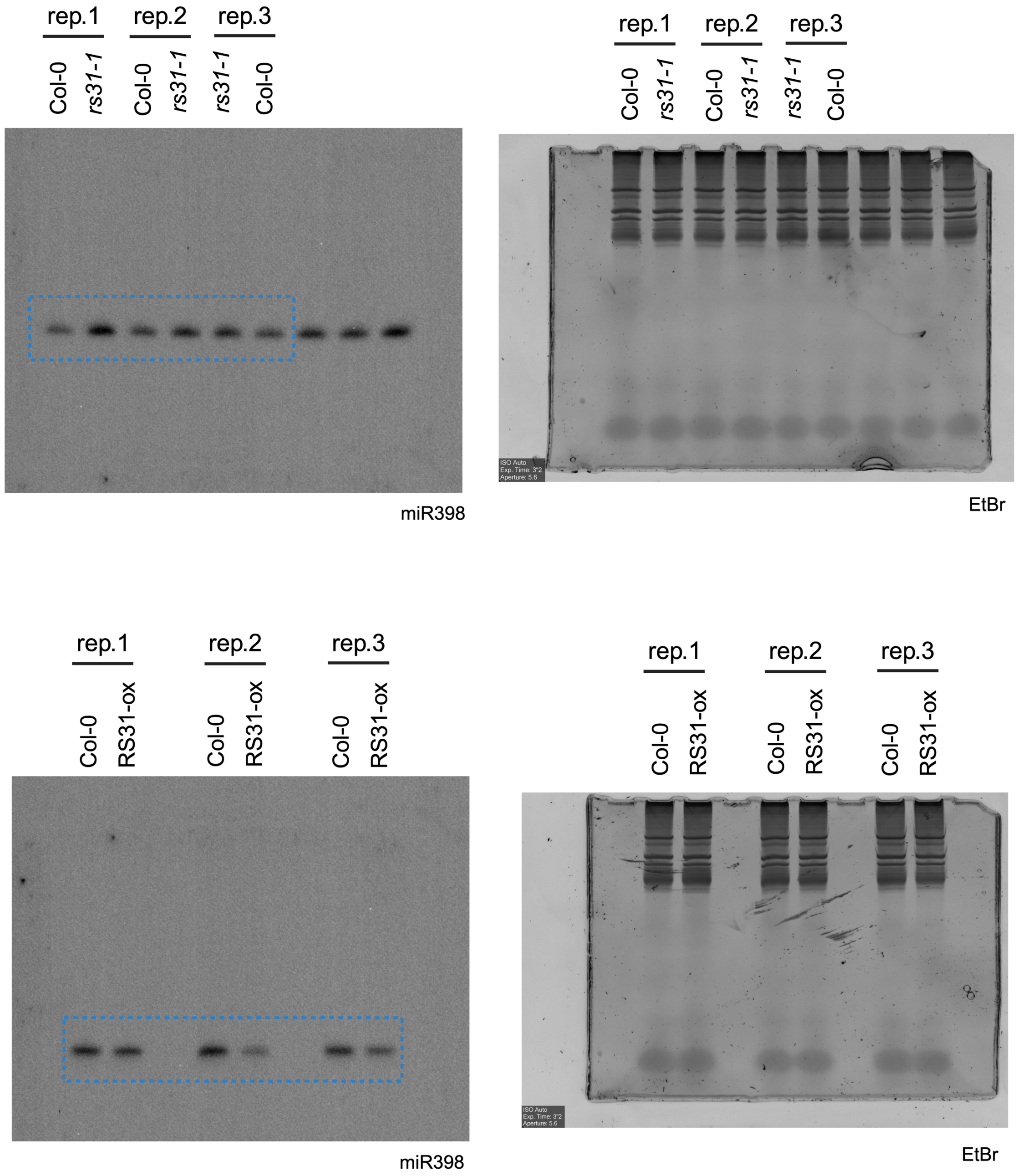


The blue dotted box indicates the area and bands used for quantification with ImageJ.

**Figure S27:** Uncropped autoradiograms to Figure S12.

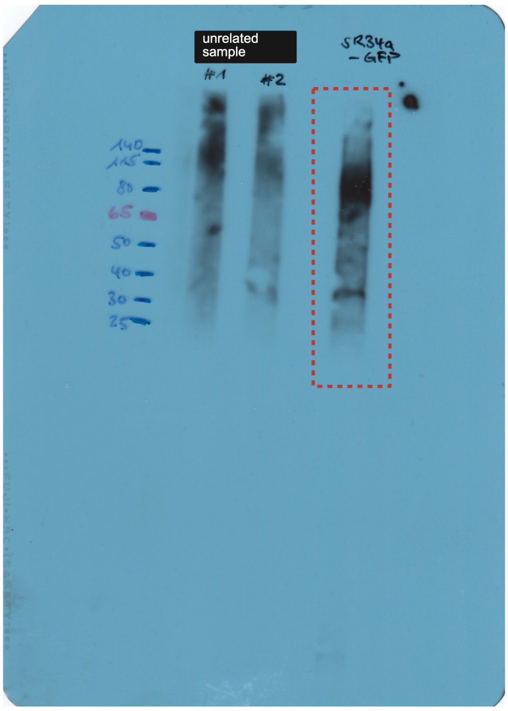


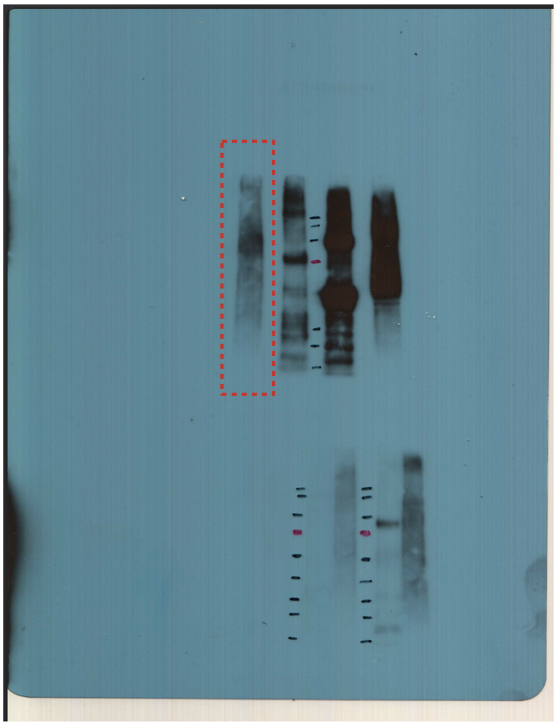

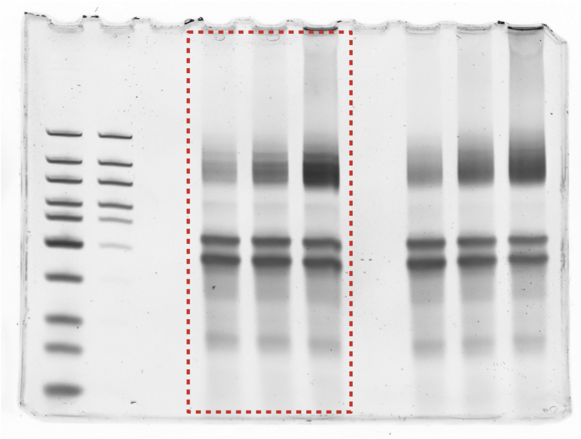

The red dotted box marks the cropped area shown in the main figure.
